# Emulating the gingival-tooth interface during bacterial, fungal, and viral infection in a microphysiological model of the human oral cavity

**DOI:** 10.64898/2026.06.11.731421

**Authors:** Mousa Younesi, Pouria Fattahi, Zhi Ren, Won Dong Lee, Sara Cherry, Hyun Koo, Dan Dongeun Huh

## Abstract

The anatomical complexity and distinctive tissue environment of the human oral cavity pose major challenges to modeling oral infection and host-microbe interactions in preclinical laboratory settings. Here we present a bioengineered oral microphysiological system comprising vascularized human gingival tissue integrated with tooth analogs that together recreate a functional unit of the human oral cavity. We incorporated *Streptococcus mutans* and *Candida albicans* into this system to model cross-kingdom biofilm formation, microbial dissemination, and host-microbial interactions at the gingival-tooth interface. Single-cell RNA sequencing and global metabolomics analysis revealed that fungal colonization induces epithelial-to-mesenchymal transition associated with distinct transcriptional and metabolic signatures. Our platform also allowed us to simulate SARS-CoV-2 infection and examine gingival responses to live-virus challenge. Finally, we integrated the engineered gingival tissue with controlled human saliva flow to show that hyposalivation potentiates the pathogenic capacity of fungal infection. This work demonstrates the potential of oral microphysiological systems as an experimental platform for in vitro modeling and mechanistic investigation of host-microbe interactions under controlled, human-relevant conditions.

## Introduction

The juxtaposition of soft and hard tissues is a fundamental principle of human anatomy that enables seamless structural integration of the specialized building blocks of the human body and their functional coordination required to support complex physiological processes. Among the prominent examples is the gingiva, the soft masticatory mucosa of the oral cavity commonly known as the gum, which is anchored to the mineralized surface of the teeth^1^ (**Fig. 1a**). The interface between these distinct tissue types, termed the gingival-tooth interface, is crucial for maintaining the mechanical stability of the teeth and provides a protective physical and immunological barrier against the constant challenge of dietary, mechanical, and chemical stimuli^1–4^. Sustained perturbation of the oral environment by poor oral hygiene, unhealthy diets, hyposalivation, and other stressors can compromise the microbial balance, increase the abundance and activity of pathogenic species, and promote biofilm formation, which is a major driver of common oral diseases including dental caries, periodontal diseases, and fungal infections^5–8^.

**Fig. 1.**
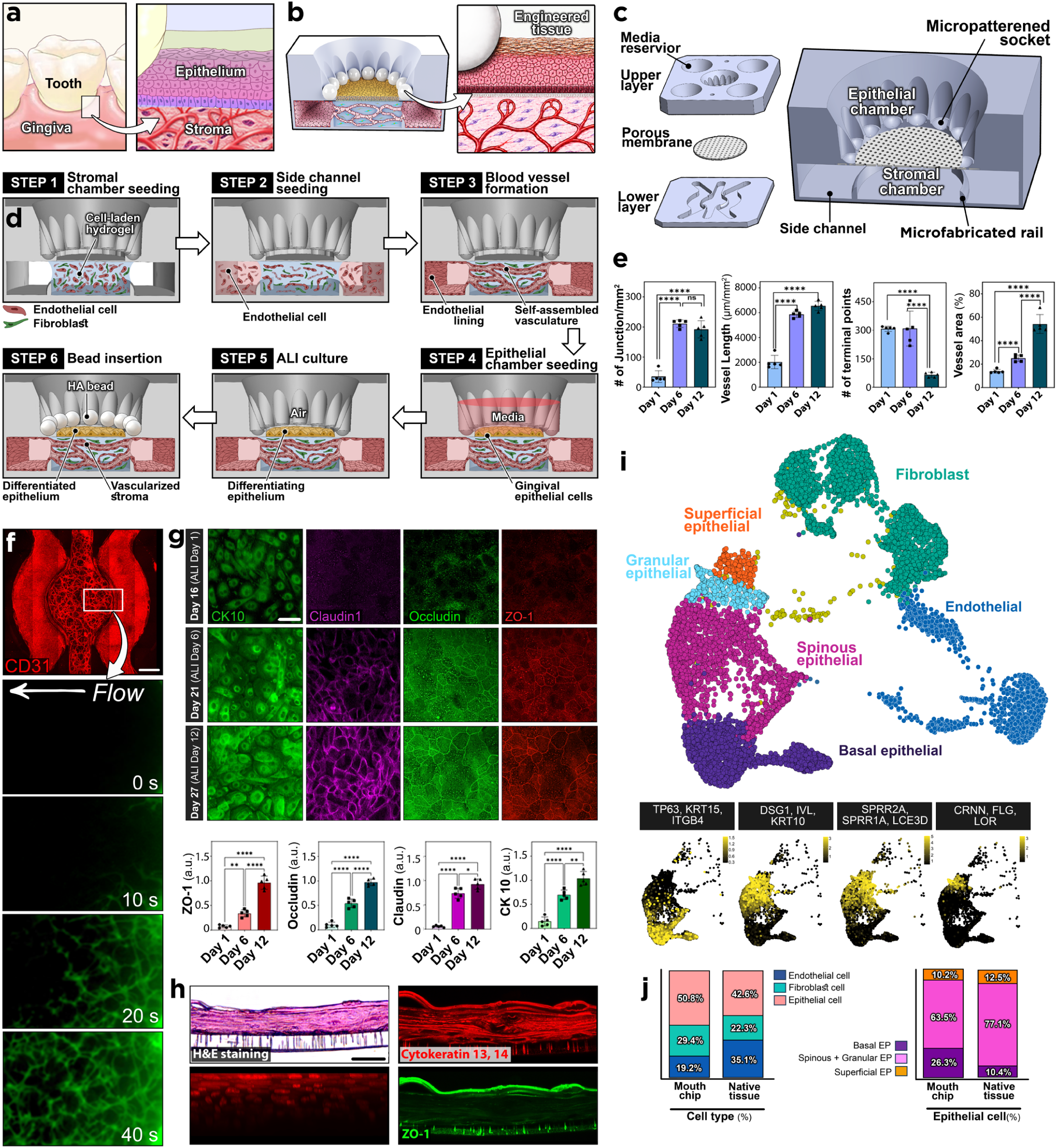
A human oral microphysiological system. **a,b,** Conceptual illustration of the gingival-tooth interface in the human mouth in vivo (**a**) and its translation into a microengineered in vitro model (**b**). **c,** Layer-by-layer and assembled views of the mouth-on-a-chip device, showing the individual components (left) and their integrated configuration (right). **d,** Sequential steps for the production of microengineered gingival tissue. **e,** Time-lapse fluorescence images of progressive blood vessel development in the stromal compartment (top) with quantification of vascular morphology and architecture (bottom). Scale bars, 100 µm. **f,** Time-lapse imaging of vascular perfusion with 70-kDa FITC-dextran (green). Scale bars, 1mm. **g,** Immunofluorescence staining of gingival epithelial differentiation marker (CK-10) and intercellular junctions. Scale bars, 50 µm. **h,** Histological and immunofluorescence analysis of the gingival epithelium harvested from the device. Scale bars, 30 µm. **i,** UMAP visualization of distinct cell types comprising the engineered gingival tissue (top) and heatmap of their cell-type-specific marker expression (bottom). **j,** Relative abundance of epithelial cell types in the mouth-on-a-chip model com-pared to the human gingiva in vivo. Data are presented as mean ± SD. n = 5 biological replicates. Statistical significance was determined by one-way ANOVA with Tukey’s multiple-comparisons test unless otherwise indicated. ns, not significant; *P < 0.05, **P < 0.01, ***P < 0.001 and ****P < 0.0001.

Biofilm-associated oral diseases represent a major global health burden, disproportionately affecting vulnerable populations and presenting persistent challenges for clinical management^9–12^. With increasing efforts to address these challenges, there is a pressing need for robust preclinical approaches to model host-oral microbe interactions and elucidate their contributions to disease pathogenesis in controlled, human-relevant settings. Notably, this need has become increasingly urgent in recent years as clinical evidence reveals the association of oral microbes with systemic diseases, including cardiovascular disease, Alzheimer’s disease, colorectal cancer, and chronic obstructive pulmonary disease^13–17^, suggesting oral health as a critical determinant of systemic wellbeing.

In general, preclinical studies of biofilm-mediated oral diseases require physiologically relevant laboratory models capable of recapitulating the human gingival-tooth interface and simulating host responses to microbial challenges^18–19^. These capabilities are essential for mechanistic investigation of how gingival tissues interact with pathogenic microbes within the oral microenvironment, how such interactions give rise to local inflammation and other adverse effects, how the pathological phenotype evolves over time, and which molecular pathways regulate key disease processes. Equally important is determining whether biofilm-associated infection has the potential to invade the gingival submucosa and disseminate into the bloodstream, thereby contributing to systemic complications. These questions represent a critical knowledge gap that needs to be addressed for the development of more effective preventive strategies and therapeutic interventions in dental medicine^20–23^.

Here we present a bioengineered model of the human oral cavity that enables advanced capabilities for investigating these key questions. This work was motivated by the scarcity of human-relevant preclinical models in oral health research and the continued dominance of animal studies as the primary source of data from which our mechanistic understanding of human oral disease is extrapolated. Conventional in vitro models have relied predominantly on monolayer cultures of human gingival epithelial cell lines, which fail to recapitulate the structural and functional complexity of the native tissue with sufficient fidelity to generate reliable preclinical data^24–25^. To overcome these limitations, three-dimensional (3D) culture techniques have been developed using primary human cells grown in Transwell or similar systems to better approximate the stratified architecture of the gingival epithelium in vivo^26–29^. Recent studies have demonstrated the feasibility of incorporating oral microbes into these models or exposing them to oral care products to evaluate cytotoxic effects^25, 28–30^, laying the foundation for more physiologically relevant in vitro studies.

Building upon these advances, we describe how the principles of 3D culture and tissue engineering can be leveraged in microfabricated devices to i) engineer a physiologically realistic representation of the human gingival-tooth unit and ii) model its integrated responses to bacterial, fungal, and viral infection. Through multi-omics analysis and molecular interrogation, we show that our microengineered system offers a preclinical research platform to generate human-relevant, high-dimensional data for mechanistic investigation of biofilm-driven infection and host defense in the human oral cavity.

## Results

### Model design

At the gingival-tooth interface, the stratified squamous epithelium of the gingiva is in direct contact with the mineralized tooth surface and supported by an underlying connective tissue that contains stromal cells and blood vessels (**Fig. 1a**). To recapitulate this tissue architecture in our microphysiological system (**Fig. 1b**), we created a device made of poly(dimethylsiloxane) (PDMS) that consists of i) an open-top epithelial chamber in the upper layer that contains a circular arrangement of micropatterned sockets designed to mimic dental arch geometry and position tooth surrogates at defined locations, ii) a stromal chamber in the lower layer for forming vascularized connective tissue underneath the epithelium, and iii) a thin, semi-permeable membrane sandwiched between the two chambers (**Fig. 1c, Supplementary Fig. 1**). The lower layer also contains two microchannels partially separated from the stromal chamber by thin, microfabricated rail structures protruding from the bottom surface (**Fig. 1c**). Each compartment in this device is accessible using its own set of fluid access ports, permitting independent control of the cellular microenvironment and compartment-specific sampling of device effluent.

The first step of model construction is to generate the vascularized subepithelial stroma by injecting an extracellular matrix (ECM) hydrogel precursor mixed with human vascular endothelial cells and gingival fibroblasts into the stromal chamber and inducing gelation to form a cell-laden hydrogel construct (**Step 1** in **Fig. 1d**). Next, an additional batch of endothelial cells is seeded into the side microchannels in the lower layer and grown on the channel surfaces (**Step 2** in **Fig. 1d**). This configuration induces vasculogenic self-assembly of endothelial cells in the hydrogel to form interconnected microvessels in the stromal compartment, which then anastomose with the endothelial lining of the side channels to generate a perfusable 3D vascular network (**Step 3** in **Fig 1d**).

To form the gingival epithelium, human gingival epithelial cells are seeded into the epithelial chamber and grown on the membrane surface (**Step 4** in **Fig. 1d**). After confluence, culture medium is removed from the chamber to establish an air-liquid interface (ALI) and drive differentiation into a stratified gingival epithelium (**Step 5** in **Fig. 1d**). Finally, hydroxyapatite (HA) beads, which are used as miniature tooth surrogates, are inserted into micropatterned sockets in the epithelial chamber to complete the construction of the gingival-tooth interface in our microengineered model (**Step 6** in **Fig 1d**).

### Gingival tissue production and characterization

First, we demonstrated the production of vascularized human gingival tissues in our model. In this study, we used primary human gingival epithelial cells and fibroblasts, as well as primary human vascular endothelial progenitor cells derived from CD34+ hematopoietic stem and progenitor cells using an established protocol^31^. The formation of the subepithelial connective tissue was accomplished by endothelial culture with fibroblasts in a fibrin hydrogel scaffold. During this period, vasculogenic activities in the stromal compartment were evidenced by increasing numbers of elongated tubular structures, which progressively joined together to form a 3D network of blood vessels throughout the hydrogel (**Fig. 1e**). Vascularization of the entire tissue construct with a diameter of 4 mm was completed by day 12 (**Fig. 1f**), at which point the vessels became perfusable, as verified by the flow of 70 kDa FITC-dextran from one side channel to the other through the vascular network (**Fig. 1f**).

Following the establishment of the fully vascularized stroma, human gingival epithelial cells were introduced into the device on day 13 and cultured for 2 days to form a confluent monolayer, after which the epithelium was maintained at ALI for additional 10 days (**Extended Figure 1**). Epithelial differentiation during ALI culture was shown by significantly increased expression of CK10, a differentiation marker of gingival epithelial cells^32^, as well as tight junction proteins characteristic of differentiated gingival epithelium, including Claudin1, Occludin, and ZO-1 (**Fig. 1g**). Histological analysis of the membrane-bound intact epithelium retrieved from our de vice revealed stacked epithelial cell layers and the localization of tight junctions at the apical surface, indicating stratification of the epithelium (**Fig. 1h**). Supporting these results, our data demonstrated the beneficial effects of ALI culture on the structural integrity of the epithelial-stromal interface and its barrier function, as evidenced by a progressive increase in electrical resistance between the epithelial and vascular compartments and a concomitant decrease in barrier permeability (**Supplementary Fig. 2**).

For further characterization, we then harvested the microengineered gingival tissues to perform single-cell RNA sequencing (scRNA-seq). UMAP analysis revealed distinct epithelial subpopulations whose gene expression profiles closely matched those of the major epithelial cell types previously identified by scRNA-seq analysis of native human gingival tissues^33–34^, including basal (TP63, KRT15, ITGB4), spinous (DSG1, IVL, KRT10), granular (SPRR2A, SPRR1A,LCE3D), and superficial squamous (CRNN, FLG, LOR) epithelial cells (**Fig. 1i, Supplementary Fig. 3**). These cell types accounted for 36.0 % (basal), 59.0 % (spinous and granular), and 5.0 % (superficial squamous) of the total epithelial population, approximating their relative proportions in the native human gingiva epithelium^35–37^ (**Fig. 1j**).

### In vitro modeling of oral biofilm formation

The mouth harbors diverse microbial species capable of assembling pathogenic biofilms linked to prevalent oral diseases^38–41^ and systemic conditions^42–45^. Among the dominant colonizers, *streptococci* and *Candida* species frequently co-occur and form interkingdom biofilms on tooth and mucosal surfaces^46^. The resulting biofilm-derived products can compromise tissue integrity at sites of colonization, while *Candida* overgrowth drives mucosal and gingival infection and may breach the underlying vascularized stroma to enter the systemic circulation^47^. Motivated by the clinical relevance of these processes, we next evaluated whether our model could support controlled interkingdom biofilm formation at a defined human tooth–gingival interface and enable concurrent measurement of host responses.

To establish an in vitro infection model, we used *Candida albicans* (*C. albicans*) and *Streptococcus mutans* (*S. mutans*), two well-characterized model fungal and bacterial species that form cross-kingdom biofilms in the oral cavity and are associated with pathogenic states through host-microbial interactions^48–52^. HA beads were first coated with saliva to promote physiologically relevant microbial attachment, and then incubated in microbial suspension for 60 minutes to initiate early colonization (**Fig. 2a**). Following insertion of colonized HA beads, quantitative imaging revealed rapid growth and biovolume expansion of both species (**Fig. 2b**), indicating active interkingdom biofilm co-development that recapitulates cooperative assembly and co-adhesion previously reported in vivo^48, 53–55^.

**Fig. 2.**
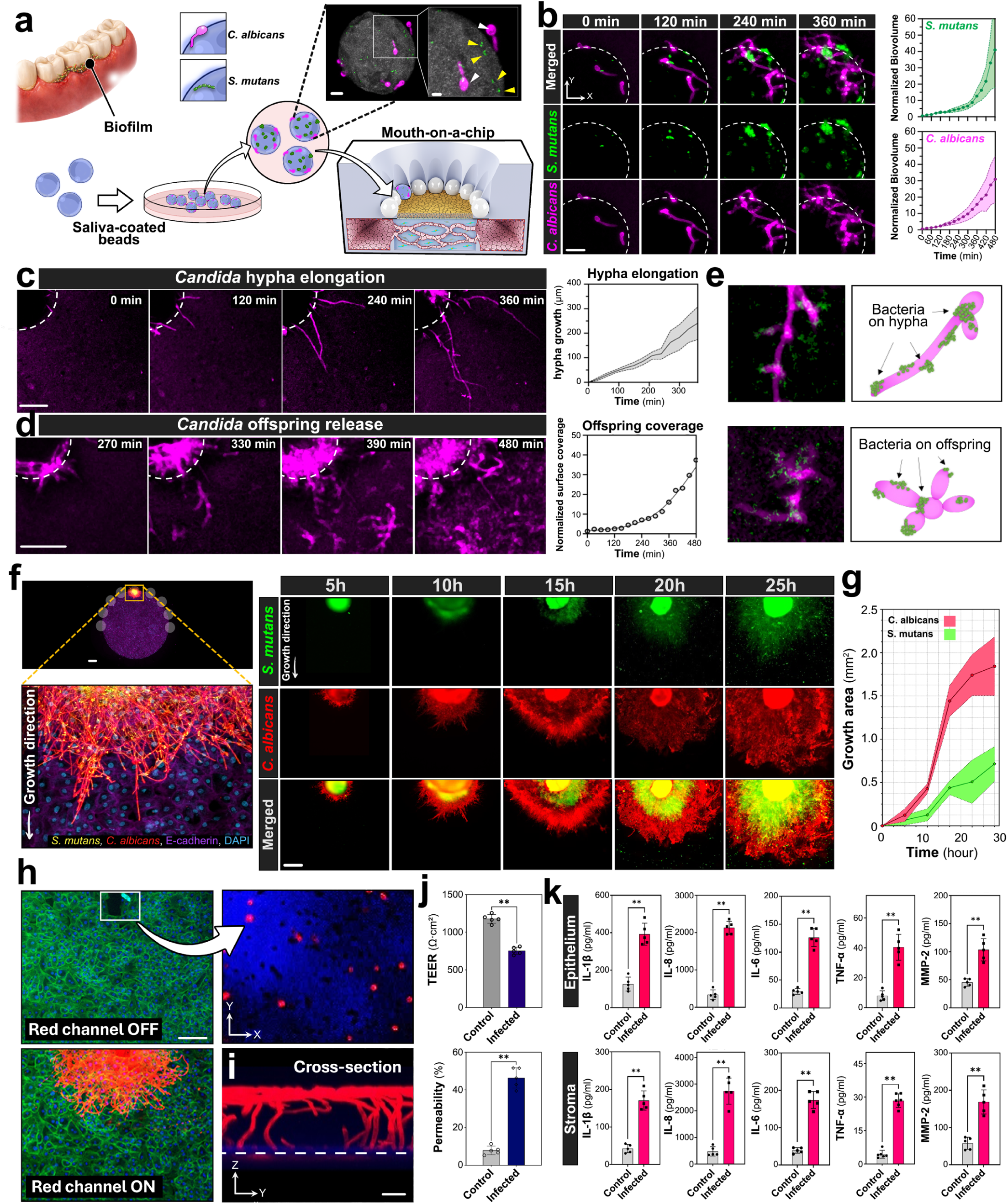
Interkingdom biofilm development in the mouth-on-a-chip. **a,** Preparation of human saliva-coated HA beads colonized with both fungal and bacterial species. The micrographs show *C. albicans* (magenta arrows) and *S. mutans* (yellow arrows) on the bead surface 30 minutes after inoculation. Scale bars, 10 µm. **b,** Fluorescence micrographs and quantification of *C. albicans* and *S. mutans* expansion on the surface of device-incorporated HA bead over time. The white dashed line shows the outline of the bead. Scale bars, 20 μm. **c,d,** Time-lapse imaging and quantification of fungal hypha growth and offspring release and spreading. The white dashed line shows the outline of the beads. Scale bars, 50 μm. **e,** Representative images of *S. mutans* adhesion to *C. albicans* hypha and off-spring. **f.** Higher-magnification view of bacterial-fungal network spreading on the gingival epithelium. Scale bar, 500 µm. **g,** Time-lapse imaging and quantification of the interkingdom biofilm expansion on the epithelial surface. Scale bars, 500 µm. **h,** Fluorescence images of the infected epithelium. The white box in the top left image indicates a region of epithelial displacement, and its magnified view shows the membrane surface (blue) and fungal hypha (red dots). Left scale bar, 100 μm **i.** Confocal micrograph showing the cross-sectional view of fungal hypha penetration through the epithelium across the porous membrane into the stromal compartment. The image was taken at 25 hours post-infection. Scale bar, 5 μm. **j.** Measurement of barrier function at 25 hours post-infection. Control represents an uninfected model. **k.** Quantification of inflammatory cytokines in the epithelial and vascular effluent samples. Data are presented as mean ± SD with n = 5. ns: not significant, *P < 0.05, **P < 0.01, ***P < 0.001, and **** P < 0.0001.

Spatiotemporal imaging at the HA-epithelium interface demonstrated species-specific behaviors during biofilm development. Both species co-colonized the saliva-conditioned surface and formed structured biofilms in which *S. mutans* clusters were interwoven with *C. albicans* hyphal forms, with both fungal and bacterial biomass increasing over time (**Fig. 2b**). *C. albicans* rapidly transitioned to filamentous growth from the HA surface, with hyphae extending over 300 μm within 360 minutes (**Fig. 2c**). Notably, time-lapse imaging also captured an unexpected event. *Candida* cells shed from the HA surface and seeded the adjacent gingival epithelium, followed by rapid clonal expansion of the newly deposited cells across the epithelial surface (**Fig. 2d**). *S. mutans* were consistently detected on both extending hypha and dispersed fungal cells (**Fig. 2e**), indicating sustained physical association during both biofilm growth and dispersal. These findings reveal a dispersal-and-seeding mechanism for interkingdom biofilm propagation to the tissue surface, consistent with clinical evidence of *C. albicans*-*S. mutans* co-localization in human plaque and saliva^51^.

Over longer time periods, the biofilm expanded radially from the infected HA bead, covering approximately 15% of the epithelial surface within 25 hours of infection (**Fig. 2f**). This expansion was driven by the extension of *C. albicans* filaments, which preceded the advancing biofilm front (**Fig. 2f**). Meanwhile, *S. mutans* accumulated as clustered cells closely associated with fungal filaments as total biomass increased (**Figs. 2f, 2g**). When we performed single-species infection experiments using saliva-conditioned HA beads coated with either *C. albicans* or *S. mutans* biofilms (**Extended Data Fig. 2**), *C. albicans* cells spread radially from the beads within 10 hours and progressively increased epithelial surface coverage at rates comparable to those observed during co-infection, whereas *S. mutans* remained largely confined to the initial deposition site (**Extended Data Fig. 2a-2c**). Suggesting that fungal hyphae provide as adhesive scaffolds for spatial expansion of *S. mutans*, these patterns of spatial organization are consistent with proximity-dependent interactions reported for *C. albicans* and *S. mutans*, including co-adhesion, metabolite exchange, and chemical signaling, that can reinforce bacterial persistence on fungal structures while promoting fungal filamentation and outward biofilm expansion ^51, 56–57^.

Importantly, regions of extensive fungal growth and hypha formation were associated with displacement of gingival epithelial cells and exposure of the underlying surface (**Fig. 2h**, **2i**). Confocal microscopy revealed hyphal penetration through the porous membrane into the stromal compartment (**Fig. 2i**), consistent with fungal filamentous invasion observed in vivo^58^. Supporting these indicators of compromised tissue integrity, at 25 hours post-infection, we observed an approximately 35% decrease in electrical resistance and a nearly 5-fold increase in barrier permeability relative to uninfected controls (**Fig. 2j**). ELISA of device effluent collected at the same time point indicated increased inflammatory responses in both the epithelial and stromal compartments, as evidenced by significant upregulation of inflammatory mediators (**Fig. 2k**). Notably, comparison of these data to those from single-species infection experiments indicated significantly greater contributions of *C. albicans* to biofilm-induced epithelial dysfunction (**Extended Data Fig. 2d-2f**).

Collectively, these results demonstrate that our platform supports microbial colonization of the microengineered gingival-tooth interface to emulate interkingdom biofilm development in a physiologically relevant manner and recapitulate biofilm-mediated epithelial barrier disruption and inflammatory responses.

### Emergence of hybrid epithelial-mesenchymal state due to infection

Next, we conducted scRNA-seq to examine transcriptomic profiles of our model following infection with *C. albicans* and *S. mutans* and investigate host-microbe interactions at the gingival interface. As the most notable finding, UMAP analysis revealed the emergence of a cell cluster that was nearly absent in the uninfected control model (demarcated in **Fig. 3a**). Positioned between the epithelial and fibroblast clusters in the UMAP plot, this population was identified by its expression of both epithelial (e.g., KRT5, KRT14 and KRT16) and mesenchymal (e.g., VIM, COL1A1, FN1) markers (**Fig. 3b**), which was further supported by differential gene expression analysis (**Fig. 3c**). Pseudotime trajectory analysis indicated that this population originated from basal epithelial cells and transitioned through the spinous and granular epithelial states before acquiring concurrent epithelial and mesenchymal features (**Fig. 3d**). Gene expression profiles along pseudotime showed upregulation of VIM, FN1, and other mesenchymal markers with the emergence of this population, whereas the expression of epithelial markers, such as KRT14 and KRT5, was sustained throughout cell type transition (**Fig. 3e**).

**Fig. 3.**
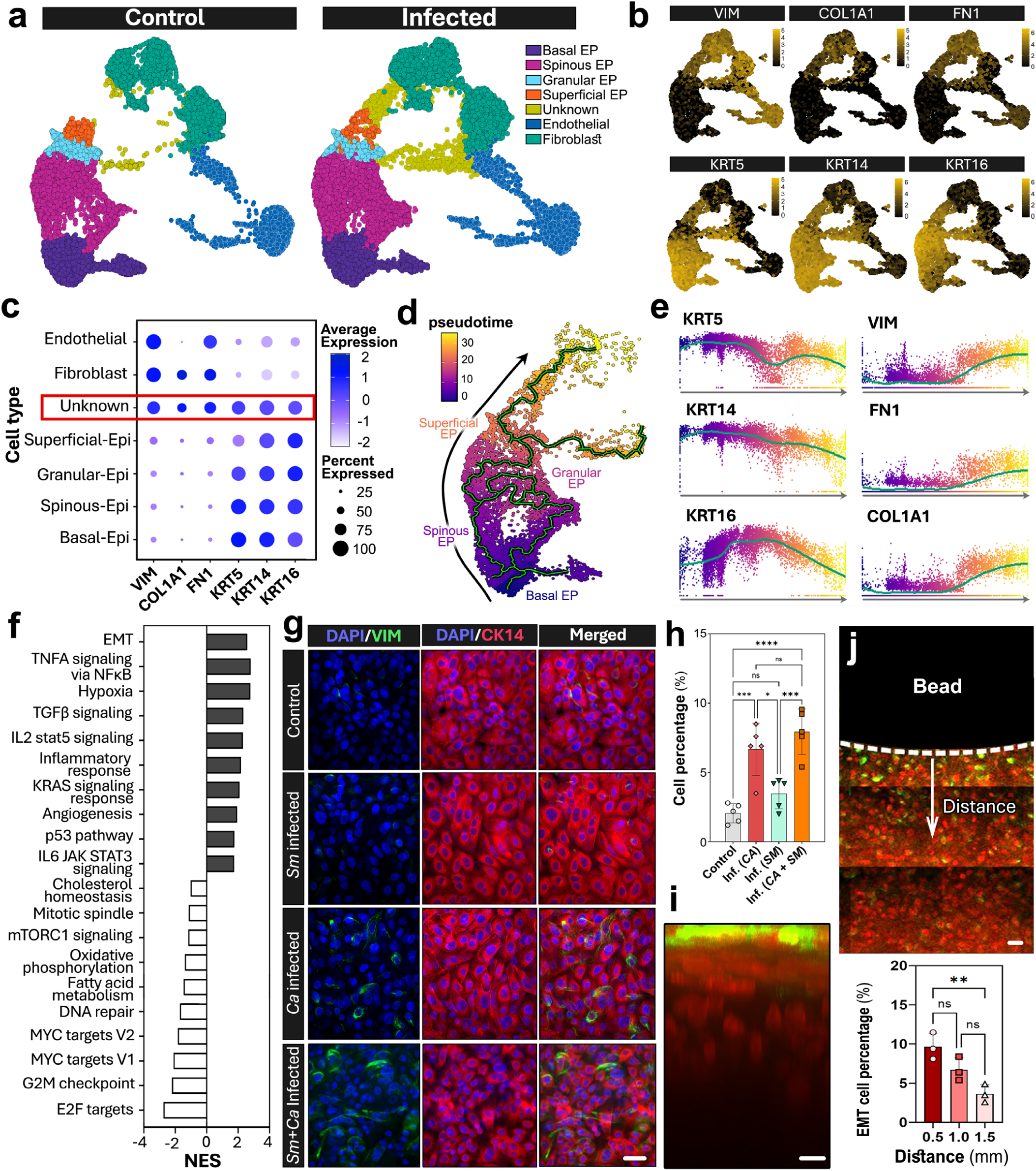
Interkingdom infection-induced EMT-like epithelial state in the mouth-on-a-chip. **a,** UMAP visualization and comparison of distinct cell types in the control and infected groups. The epithelial cell population is outlined. **b,** Feature plots showing the expression of mesenchymal (top row) and epithelial (bottom row) markers on the UMAP. **c,** Dot plot of mesenchymal (VIM, COL1A1, FN1) and epithelial (KRT5, KRT14, KRT16) marker expression across cell types in the engineered gingival tissue. Dot size indicates the percentage of cells expressing each gene, whereas color indicates scaled average expression. The outlined population (red box) co-expresses both mesenchymal and epithelial markers. **d,** Pseudotime trajectory of epithelial cells, colored by inferred pseudotime with the fitted trajectory shown in green. Black arrow indicates the direction of inferred progression. **e,** Expression dynamics of epithelial and mesenchymal along pseudotime in the infected model. Each point represents a cell colored by pseudotime. The fitted curves indicate smoothed expression trends. **f,** GSEA of the infected model. Bars indicate normalized enrichment scores (NES) for each gene set. Dark grey and white denote positive and negative enrichment, respectively. **g,h,** Immunofluorescence staining of CK14 and vimentin (VIM) expression in the gingival epithelium (**g**) with quantification (**h**). Scale bar, 30 µm. **i,** Representative cross-sectional image showing the vertical distribution of CK14/Vimentin-positive epithelial cells within the infected epithelium. Red and green show immunofluorescence of CK14 and vimentin, respectively. Scale bar, 10 µm. **j,** Spatial analysis of CK14/Vimentin-positive epithelial cell abundance relative to the colonized bead. Red and green show immunofluorescence of CK14 and vimentin, respectively. Scale bar, 40 µm. Data are presented as mean ± SD; n = 5 biological replicates. Statistical significance was determined by one-way ANOVA with Tukey’s multiple-comparisons test. ns, not significant; *P < 0.05, **P < 0.01, ***P < 0.001 and ****P < 0.0001.

These data suggested that interkingdom biofilm infection of our model with *C. albicans* and *S. mutans* induced transcriptional reprogramming of gingival epithelial cells toward what is often described as a hybrid epithelial-mesenchymal state or partial epithelial-mesenchymal transition (EMT) in recent literature^59–61^. Indeed, gene set enrichment analysis (GSEA) using the Hallmark gene sets from the Molecular Signatures Database (MSigDB) identified EMT as one of the most significantly enriched pathway in the infected tissues (**Fig. 3f**). Given that the capacity of interkingdom biofilms to induce EMT-like epithelial plasticity has not been well-defined, we next examined this phenotype within the integrated tissue environment of our model.

Notably, immunofluorescence analysis of the gingival epithelium at 25 hours post-infection showed a subset of cells expressing not only cytokeratin-14 (CK14) but also vimentin (**Fig. 3g**), a canonical mesenchymal marker widely recognized as a key feature of cells undergoing epithelial to mesenchymal transition^62–64^. The CK14/VIM double-positive cells accounted for approximately 10% of the epithelial population and were detected only in less than ∼3% of the cells in the uninfected control group (**Figs. 3g, 3h**). The significant increase in the abundance of this cell type was only observed in the presence of *C. albicans* (**Figs. 3g, 3h**). In the infected model, the majority of the vimentin-positive epithelial cells were localized to superficial layers of the stratified epithelium, with only few cells observed in deeper epithelial strata (**Fig. 3i**). Moreover, the abundance of these cells was highest at and immediately adjacent to the gingival-tooth junction and decreased with distance across the epithelial surface (**Fig. 3j**).

### Molecular interrogation of EMT-like phenotype

Our data indicating the emergence of a hybrid epithelial-mesenchymal phenotype in the infected model led us to investigate the molecular basis of this response. To gain a high-level overview of infection-induced signaling activities, we first performed pathway enrichment analysis using the Kyoto Encyclopedia of Genes and Genomes (KEGG) ^65–66^.

Among the most enriched pathways were signaling modules downstream of growth factor receptor tyrosine kinases (GF-RTKs), including GF-RTK-PI3K, GF-RTK-RAS-ERK, and GF-RTK-RAS-PI3K (**Fig. 4a**). Components of these cascades have been implicated in EMT in oral epithelia, including activation of PI3K/AKT and ERK signaling in single-species infection of traditional monolayer gingival cultures, which can reduce epithelial adhesion and induce EMT-associated transcription factors, such as ZEB1, SLUG, and SNAIL ^67–69^. Extending these observations to an interkingdom biofilm context, our findings suggest that co-infection with *C. albicans* and *S. mutans* in the tissue microenvironment of our model may have the capacity to trigger coordinated activation of GF-RTK signaling programs associated with EMT-like transcriptional reprogramming of human gingival epithelial cells.

**Fig. 4.**
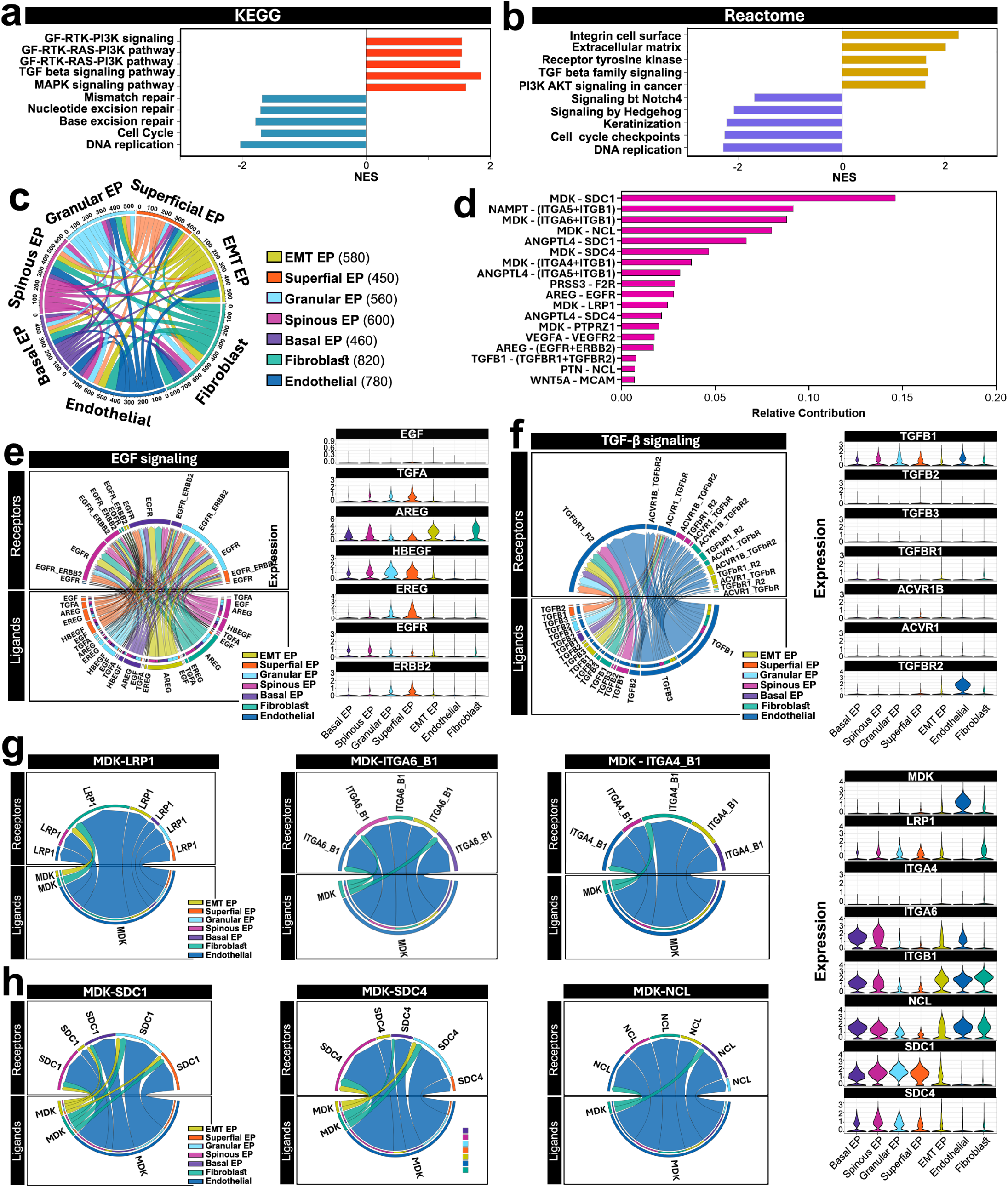
Analysis of molecular pathways involved in infection-associated EMT state. **a,b,** Gene set enrichment analysis (GSEA) of differentially expressed genes (control vs. infected) against the KEGG (**a**) and Reactome (**b**) pathway databases. Bars show the normalized enrichment score (NES) for pathways meeting an FDR q-value < 0.05. **c, c,** Global cell–cell communication network inferred using the CellChatDB ligand–receptor interaction database and visualized as a chord diagram. Each sector represents a cell cluster, with the outer scale indicating the number of inferred ligand–receptor interactions involving that cell type. The width of each chord is proportional to the number of interactions between the connected sender and receiver pair, and chord color denotes the sender cell type. Values in parentheses in the legend represent the total number of inferred interactions in which each cell type participates as sender or receiver. **d,** Top ligand–receptor pairs ranked by relative contribution to all inferred signaling in the infected model. **e,f,** Inferred EGF (**e**) and TGF-β (**f**) signaling networks among cell types shown as chord diagrams in which arcs split into ligand- and receptor-expressing partners to indicate direction. Chord width is proportional to interaction strength. Adjacent violin plots show log-normalized expression per cell across cell types for EGF-family ligands (EGF, TGFA, AREG, HBEGF, EREG) and receptors (EGFR, ERBB2) in **e**, and TGF-β ligands (TGFB1–3) and receptors/co-receptors (TGFBR1, ACVR1B, ACVR1, TGFBR2) in **f**. **g,h,** MDK-centered signaling decomposed by receptor. MDK–LRP1, MDK–ITGA6/ITGB1, MDK–ITGA4/ITGB1 in (**g**) and MDK–SDC1, MDK–SDC4, MDK–NCL in (**h**), each with corresponding violin plots of log-normalized expression of MDK and the indicated receptors across cell types.

We next conducted further interrogation of our sequencing data using the Reactome Pathway Database^70–71^. Importantly, among the pathways positively enriched in the infected model relative to the uninfected control were *PI3K-AKT Signaling in Cancer* and *TGF-β Family Signaling*, both of which are established drivers of EMT (**Fig. 4b**). The PI3K-AKT pathway promotes cell survival, growth, and metabolism and can stabilize EMT transcription factors^72–74^. Binding of TGF-β to its receptor triggers phosphorylation of SMAD proteins, and this signaling upregulates EMT transcription factors and mesenchymal genes while downregulating epithelial junction proteins^75–76^. These pathways reinforce each other to drive EMT^77–78^. Consistent with engagement of an EMT program, the infected model also showed positive enrichment of *Extracellular Matrix* and *Integrin Cell Surface signaling*, which represent downstream signatures of the matrix remodeling and altered cell–matrix adhesion that accompany the mesenchymal transition, together with negative enrichment of *DNA Replication* and *Cell Cycle Checkpoints* (**Fig. 4b**). This suppression of proliferative programs was reproducible across analyses, consistent with the negative enrichment of DNA replication, cell cycle, and DNA repair pathways observed by KEGG (**Fig. 4a**).

Positive enrichment and convergence of these pathways provide a molecular basis for classifying the infection-associated hybrid epithelial-mesenchymal state as an EMT-like phenotype. Notably, this integrative analysis indicates that transcriptional reprogramming of the infected epithelial cells does not arise from the activation of a single isolated pathway but rather from the coordinated engagement of GF-RTK-PI3K and TGF-β family signaling responsive programs, accompanied by transcriptional signatures of matrix remodeling and suppressed epithelial proliferation. This systems-level insight offers a broader mechanistic framework for understanding infection-induced EMT in the human gingival epithelium. To our knowledge, the capacity of interkingdom biofilms to engage such integrated signaling networks in human gingival tissue models has not been described previously.

### Intercellular crosstalk driving infection-induced EMT

To better understand biological regulation of infection-induced EMT in our model, we used CellChatDB^79^ to examine ligand-receptor interactions that mediate the crosstalk between the hybrid epithelial-mesenchymal cell population (termed “EMT cells” hereafter) and the other cell types. Of the 4250 ligand-receptor pairs identified across all cell populations, 580 involved EMT cells (**Fig. 4c**), illustrating their important role in orchestrating infection-induced biological signaling.

The top-ranked ligand-receptor pairs among these interactions included those implicated in EMT of oral epithelial cells, such as AREG-EGFR, AREG-(EGFR+ERBB2), and TGFB1-(TGFBR1+TGFBR2) ^80–82^ (**Fig. 4d, Supplementary Fig. 4**). Mapping and visualization of these data in a cell type-specific manner provided further insight into these interactions. For example, AREG-EGFR signaling occurs within the epithelium in an autocrine manner as previously described ^83–84^, but our analysis showed that the EMT cells receive AREG not only from the other types of epithelial cells but also from gingival fibroblasts in the stromal compartment (**Fig. 4e**). For TGFB1-(TGFBR1+TGFBR2), the vascular endothelium was predicted to be the primary source of TGFB1 in the subepithelial stroma (**Fig. 4f**).

Notably, our data revealed ligand-receptor interactions not previously linked to oral EMT. In particular, several ligand-receptor pairs containing midkine (MDK), a heparin-binding growth factor, ranked among the most significant interactions involving EMT cells with higher levels of statistical significance than EGF and TGF-β signaling described above. These included MDK-(ITGA6+ITGB1), MDK-(ITGA4+ITGB1), MDK-LRP1(**Fig. 4g**). In non-gingival settings, MDK is known to be significantly upregulated in many types of cancer, and its binding to low-density lipoprotein receptor-related protien 1 (LRP1) and integrin receptors has been reported to activate PI3K–AKT, ERK/MAPK, and NF-κB pathways to drive EMT ^85–89^; however, such MDK-based signaling have not been described in the context of infection-associated EMT in the human gingiva.

MDK expression was enriched in endothelial cells and fibroblasts, with comparatively lower expression in EMT epithelial cells. In contrast, EMT cells preferentially expressed several MDK-binding receptors and co-receptors, including β1-integrin heterodimers such as ITGA6–ITGB1, SDC1/SDC4, and NCL. This expression pattern suggests predominantly paracrine and potentially partially autocrine MDK signaling, in which stromal and vascular-derived MDK engages receptors on EMT cells to promote pro-survival and pro-migratory programs during infection (**Fig. 4g**).

Although sydecan 1 (SDC1) and SDC4 have been shown as negative and positive regulators of EMT in oral and other types of cancer, respectively ^90–94^, the role of their interactions with MDK in gingival EMT remains unknown. In our infected model, SDC signaling was predicted to occur through the binding of MDK produced by endothelial and fibroblasts to SDC1 or SDC4 expressed by EMT cells and other epithelial cell types (**Fig. 4h**). Similarly, nucleolin (NCL) can localize to the cell surface and function as a non-canonical receptor for MDK and other ligands. While the MDK-nucleolin (NCL) axis has been implicated in promoting cancer cell survival and metastatic progression ^95–97^, its contribution to EMT in the infected human gingiva has not been established. Our data suggest that interkingdom biofilm infection may activate this signaling by inducing MDK expression in endothelial cells and fibroblasts, enabling its engagement with NCL on the surface of EMT cells (**Fig. 4h**).

In summary, our analysis shows that the EMT cell population in the infected model receives biological inputs from both epithelial and stromal cells to engage in complex intercellular signaling that integrates classical EMT regulators with MDK-based ligand-receptor interactors as previously unrecognized drivers of infection-associated epithelia plasticity in human gingival tissue.

### Metabolic signatures of infection and infection-induced EMT

EMT-like epithelial plasticity has been associated with not only transcriptional and phenotypic alterations but also metabolic remodeling that supports cell survival, proliferation, and migration in various epithelial contexts ^100–103^. It remains poorly understood, however, how infection changes the metabolic state of human gingival tissue and whether infection-associated EMT-like tissue remodeling can generate distinct metabolic signatures.

Motivated by this knowledge gap, we profiled the metabolome of our microengineered model following *C. albicans* biofilm infection, given our finding that *C. albicans* is responsible for driving EMT-like responses (**Fig. 3**). To capture metabolic alterations in a tissue compartment-specific manner, we performed untargeted global metabolomics analysis on device effluent sampled from the epithelial and vascularized stromal compartments at 25 hours post-infection. Partial least squares discriminant analysis (PLS-DA) and hierarchical clustering heatmaps showed clear separation between control and infected samples in both the epithelial and stromal compartments (**Figs. 5a, 5b; Supplementary Figs. 5-8**), indicating broad fungal infection-associated metabolic remodeling.

**Fig. 5.**
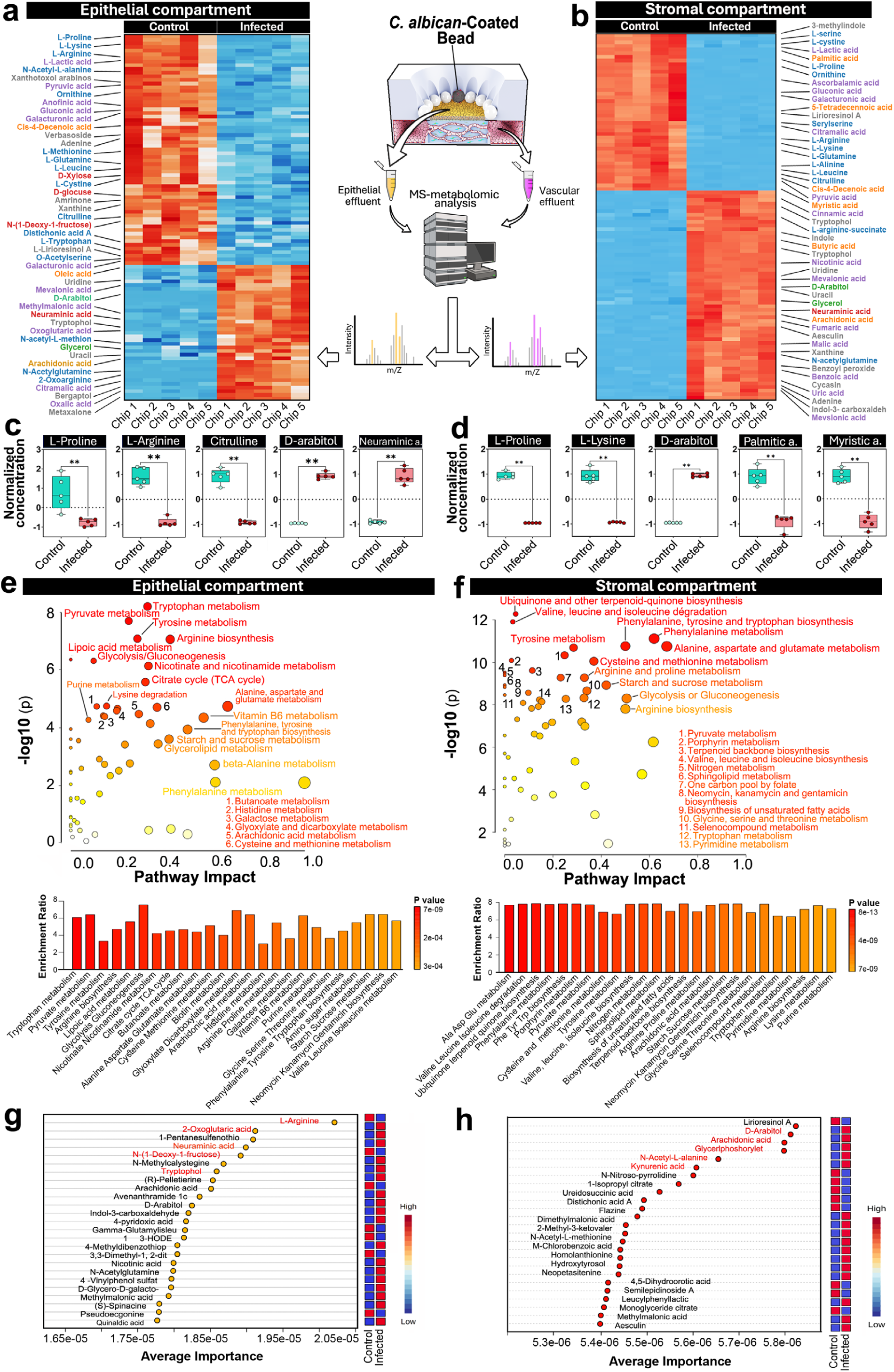
Metabolomic analysis of *C. albicans*–infected gingival tissue model. **a,b,** Heatmaps of untargeted LC–MS metabolomics profiles measured from the epithelial (**a**) and stromal (**b**) chambers. Metabolite classes are color-coded as follows – blue: amino acids and amino-acid derivatives, red: carbohydrates (sugars and sugar derivatives), green: sugar alcohols/polyols, beige: fatty acids/lipids, purple: organic acids. **c,d,** Representative epithelial (**c**) and stromal (**d**) metabolites significantly altered by infection. Individual points represent biological replicates. Box plots show minimum, 25th percentile, mean, and 75th percentile. **e, f,** Pathway impact and enrichment analysis of significantly altered metabolites in the epithelial (**e**) and stromal (**f**) compartments. **g, h,** VIP-based feature importance from PLS-DA identifying the top discriminatory epithelial (**g**) and stromal (**h**) metabolites that separate infected from control groups. Statistical significance was determined by two-tailed unpaired Student’s *t*-test for two-group comparisons. Data are presented as mean ± SD with n = 5. ns, not significant, *P < 0.05, **P < 0.01, ***P < 0.001, and ****P < 0.0001.

Among the significantly changed metabolites, infection significantly depleted epithelial L-proline (**Fig. 5c**), a metabolite critical for the synthesis of collagen and other ECM proteins at the gingival-tooth interface^105–106^. Given the established role of proline in maintaining the structural integrity of the junctional epithelium, this decrease is consistent with infection-induced impairment of epithelial barrier function. Also reduced were citrulline and L-arginine (**Fig. 5c**), indicating the perturbation of the citrulline-arginine axis in the infected tissue microenvironment. This may reflect the suppression of host nitric oxide (NO) pathways used for antimicrobial defense^107–108^ and/or increased consumption of these amino acids by *C. albicans*.

Epithelial metabolites elevated by fungal infection included D-arabitol and tryptophol (**Fig. 5c, Supplementary Fig. 9**). D-arabitol is a fungal polyol not synthesized in substantial quantities by mammals under normal physiological conditions^109^, making it a well-established biomarker of fungal metabolic activity. Tryptophol is an aromatic alcohol derived from tryptophan by *C. albicans*, and its elevation in our model is consistent with increased fungal tryptophan catabolism during infection^110–111^. Epithelial inflammatory responses to infection were supported by elevated levels of arachidonic acid (**Supplementary Fig. 9**), a precursor to pro-inflammatory eicosanoids implicated in gingival inflammation^112^. Increased levels of neuraminic acid were another notable feature of the epithelial metabolic profile (**Fig. 5c**). Considering that sialic acid is a key component of glycoproteins and glycolipids constituting the protective epithelial glycocalyx^113^, its elevation in the effluent may reflect infection-induced remodeling or degradation of the host glycocalyx. This finding is consistent with clinical evidence demonstrating increased salivary levels of sialic acid as potential biomarkers of periodontal disease^114^.

Several metabolites, including L-proline, L-lysine, and D-arabitol were also detected in the stromal effluent and exhibited similar infection-associated trends (**Fig. 5d, Supplementary Fig. 10**). Of note, the stromal compartment displayed a distinct signature characterized by significantly decreased levels of several fatty acids, including palmitic acid, myristic acid, and 5-tetradecenoic acid (**Fig. 5d, Supplementary Fig. 10**). This pattern may reflect increased fatty acid utilization or altered lipid handling (e.g., reduced lipolysis, altered membrane incorporation) in the infected stromal micro-environment. Given that fatty-acid metabolism has been linked to EMT programs in other tissue contexts^115–117^ and that we independently observe an infection-associated EMT-like state in this model, these metabolic changes may represent a complementary indicator of infection-induced EMT-like responses.

Metabolic pathway analysis provided further evidence for infection-induced metabolic reprogramming in the epithelial compartment. Glycolysis/gluconeogenesis and pyruvate metabolism were among the most significantly enriched and high-impact pathways (**Fig. 5e**), indicating perturbation of central carbon metabolism following infection. Such central carbon reprogramming is often associated with EMT, in which altered bioenergetic demand accompanies the transition of epithelial cells towards a migratory, mesenchymal phenotype^118^. Amino acid-related pathways, including tryptophan, tyrosine, and arginine metabolism, were also significantly impacted (**Fig. 5e**). These pathways can generate bioactive metabolites that can activate EMT-associated transcription factors. For example, tryptophan derivatives contribute to EMT in skin epithelial cells by downregulating E-cadherin while increasing the expression of SNAIL, SLUG, and TWIST^119^. Although arginine metabolism has not been directly linked to EMT, arginine availability can influence arginine methylation, which represents a post-translational modification implicated in EMT regulation^120^.

The infected stroma showed enrichment of metabolic pathways spanning a broader range of amino acids, including cysteine, alanine, valine, and leucine (**Fig. 5f**), providing a pathway signature distinct from that of the epithelial compartment. Pathways uniquely enriched in the stroma included ubiquinone and other terpenoid-quinone biosynthesis (**Fig. 5f**). Ubiquinone, also known as coenzyme Q10 (CoQ10), functions as an essential electron carrier in the mitochondrial electron transport chain^121^, and enrichment of this pathway may reflect altered mitochondrial activity in the infected stromal microenvironment. Since ubiquinone also acts as a lipid-soluble antioxidant and links fatty acid beta-oxidation to the respiratory chain^122^, its increased biosynthesis could alternatively represent a compensatory response to replenish CoQ10 pools consumed under infection-induced oxidative stress.

Finally, we performed variable importance in projection (VIP) and receiver operating characteristic (ROC) analyses to explore the feasibility of identifying candidate biomarkers of infection-induced metabolic changes in our model (**Supplementary Figs. 11, 12**). This analysis revealed i) downregulation of L-arginine and N-(1-Deoxy-1-fructosyl)glycine and ii) upregulation of neuraminic acid, 2-oxoglutaric acid, and tryptophol as the top discriminatory epithelial metabolites separating the infected model from uninfected controls (**Fig. 5g, Supplementary Fig. 13**). In the stromal compartment, the top discriminatory metabolites were D-arabitol, arachidonic acid, glycerylphosphorylethanolamine, N-acetyl-L-alanine, and Kynurenic acid, all of which were elevated by infection (**Fig. 5h, Supplementary Fig. 14**). Apart from L-arginine, neuraminic acid, D-arabitol, and arachidonic acid, the infection-associated behavior of these metabolites have not been characterized in the context of fungal gingival infection or EMT-like remodeling of the human gingiva.

### Modeling gingival infection with SARS-CoV-2

Next, we explored whether our microphysiological system could be used to model viral infection in the human oral cavity. Motivated by the global health threat posed by coronaviruses^123–124^, we used SARS-CoV-2 as a model pathogen for this proof-of-concept study (**Fig. 6a**). Although oral manifestations of SARS-CoV-2 infection, such as altered taste perception and dry mouth, have been well-documented^125^, accumulating evidence suggests that the oral cavity is more than a site of secondary symptoms. The gingival epithelium expresses angiotensin-converting enzyme 2 (ACE2) and associated entry factors, including transmembrane protease serine 2 (TMPRSS2) and furin, which facilitate viral entry and transmission^126^. Based on recent reports of SARS-CoV-2 detected in human gingival tissues and gingival crevicular fluid in COVID-19 patients^127–128^, it has been suggested that the periodontal pockets at the gingival-tooth interface may act as viral reservoirs that enable local persistence and systemic dissemination^129^. Despite these insights, the direct effects of SARS-CoV-2 on human gingival tissues remain poorly understood.

**Fig. 6.**
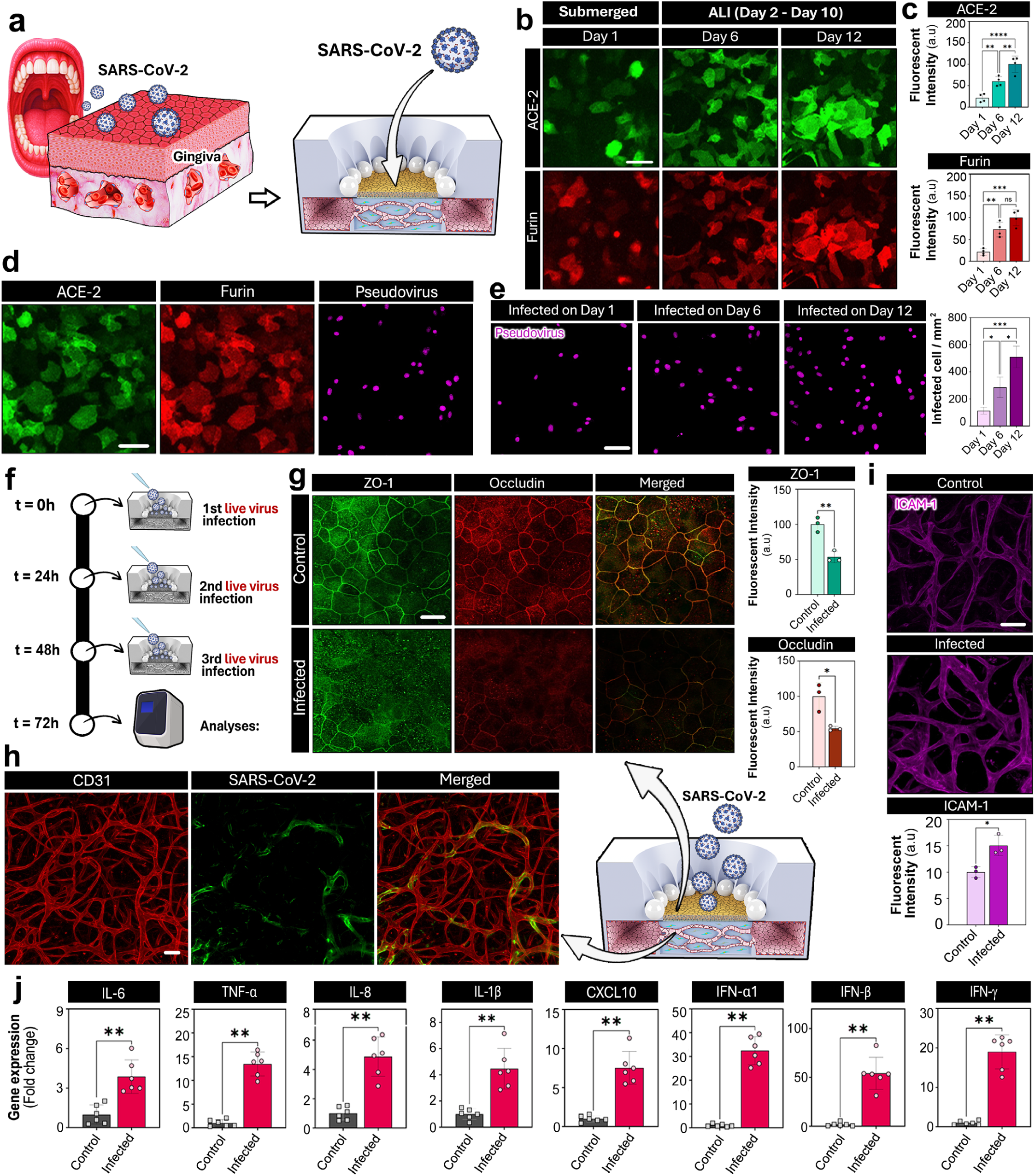
Infection of microengineered gingival tissue with SARS-CoV-2. **a,** In vitro modeling of gingival infection with SARS-CoV-2 in the mouth-on-a-chip. **b,c,** Immunofluorescence analysis of ACE2 and furin expression in the gingival epithelium at different stages of tissue production. Scale bars, 50 µm. **d,** Images of the gingival epithelium treated with SARS-CoV-2 pseudovirus (magenta) at 24 hours post-infection. Scale bars, 50 µm. **e,** Pseudovirus (magenta) on the epithelial surface imaged in tissues infected at different time points during culture. Scale bar, 50 µm. **f,** Timeline for live, wild-type SARS-CoV-2 infection experiments. **g,** Immunofluorescence analysis of epithelial ZO-1 and occludin following wild-type SARS-CoV-2 exposure. The images were taken 24 hours after the 3^rd^ infection shown in **f**. Scale bars, 50 µm. **h,** Immunofluorescence detection of SARS-CoV-2 in the blood vessels of the stromal compartment. Scale bar, 50 µm. **i,** Immunofluorescence analysis of ICAM-1 expression in the microvascular network. Scale bar, 50 µm. **j,** RT–qPCR of epithelial lysates showing induction of inflammatory cytokines and antiviral response genes following SARS-CoV-2 infection. Data are presented as mean ± SD with n = 3–5 biological replicates per condition. Statistical significance was determined by two-tailed unpaired Student’s *t*-test for two-group comparisons and one-way ANOVA with Tukey’s multiple-comparisons test for comparisons among three or more groups. ns, not significant; *P < 0.05, **P < 0.01, ***P < 0.001 and ****P < 0.0001.

Immunofluorescence analysis confirmed the expression of both ACE2 and furin in confluent monolayers of gingival epithelial cells produced by submerged culture for 2 days in our model (**Fig. 6b**). During subsequent ALI culture for additional 10 days, the expression of these markers increased by more than 4-fold (**Figs. 6b, 6c**), suggesting progressive acquisition of viral susceptibility with epithelial maturation. To assess tissue infectivity, we challenged the epithelial compartment with a replication-deficient SARS-CoV-2 pseudovirus carrying a fluorescent protein reporter. Examination at 24 hours post-infection revealed pseudovirus particles distributed across the gingival surface and retained within the epithelium (**Fig. 6d**), indicating viral entry into the cells. Consistent with the temporal profiles of ACE2 and furin expression, longer ALI culture resulted in significantly higher numbers of infected cells (**Fig. 6e**), illustrating the importance of epithelial maturity for properly modeling viral entry at the gingival mucosa.

When our model was challenged with live, wild type SARS-CoV-2 at MOI = 0.3 every 24 hours for 3 consecutive days (**Fig. 6f**), the infected epithelium appeared intact without apparent tissue injury, but our analysis showed significantly reduced expression of tight junction proteins (**Fig. 6g**). Notably, the virus was also detected in the blood vessels of the subepithelial stroma and their vicinity (**Fig. 6h**). Compared to the uninfected control, the endothelium of the infected vasculature expressed significantly higher levels of intercellular adhesion molecule-1 (ICAM-1) (**Fig. 6i**).

Further investigation using RT-qPCR revealed inflammatory responses of our model to infection. The epithelium upregulated expression of pro-inflammatory cytokines characteristic of classical innate immune responses at mucosal surfaces, including IL-6, IL-8, TNF-α, and IL-1β (**Fig. 6j**), which was accompanied by elevated levels of IFN-α1, IFN-β, and IFN-γ (**Fig. 6j**), indicative of an antiviral program involving both Type I (IFN-α1, IFN-β) and Type II (IFN-γ) interferon signaling. Notably, the epithelial response also included significantly increased expression of CXCL10 (**Fig. 6j**). As an interferon-induced potent chemoattractant for activated T cells and NK cells, CXCL10 has been consistently identified as one of the most elevated cytokines in severe COVID-19^130–131^ and shown to play a key role in mediating epithelial-endothelial crosstalk during SARS-CoV-2 infection^132^.

Supporting the emerging view of the human gingiva as an active port of entry for SARS-CoV-2^133–134^, these results demonstrate the capacity of SARS-CoV-2 to compromise the integrity of the gingival epithelial barrier, translocate into the subepithelial stroma, and induce spatially distinct, physiologically relevant inflammatory and antiviral responses that may be useful for investigating immunopathology of COVID-19 in the human oral cavity.

### Modeling hyposalivation and its effect on gingival infection

Saliva is a complex biofluid that plays a crucial role in oral homeostasis by modulating host responses to pathogenic microbes^135^. Under conditions of reduced salivary flow (hyposalivation), thinning of the salivary film on mucosal surfaces compromises these protective mechanisms and increases susceptibility to fungal overgrowth and biofilm-associated infection^136^. While the antimicrobial properties of saliva provide a well-established biological basis for protection against fungal infection^137^, the direct effects of hyposalivation on gingival responses to fungal pathogens remain incompletely understood, particularly in human-relevant contexts. This knowledge gap is clinically relevant given the increasing prevalence of hyposalivation with aging and demographic shifts toward older populations worldwide^138–139^.

Recognizing the inability to recapitulate salivation as an important limitation of our microphysiological system, we next modified the design of our platform to enable in vitro reproduction and precise control of saliva flow, with the goal of investigating the effect of hyposalivation on gingival infection. To model salivation, we integrated a set of microfluidic channels into the upper layer of the device to generate pressure-driven, controlled inflow and outflow of saliva across the epithelial compartment (**Fig. 7a, Supplementary Fig. 15**). Pooled human saliva was delivered into the epithelial compartment through two symmetrically positioned inlet microchannels with a diameter of 100 μm (**Figs. 7a, 7b**). The flow was driven at 120 μl/h, which was determined by our scaling analysis to approximate physiological conditions of saliva flow^140^. Upon entering the open-top epithelial chamber, saliva spread to cover the entire epithelial surface and exited through a dedicated outlet microchannel, permitting continuous renewal of the salivary layer (**Fig. 7c**). Under steady-state flow, the liquid layer over the epithelium reached an average thickness of approximately 110 μm (**Fig. 7d**), comparable to the reported thickness of the salivary film in the human mouth (70-100 μm)^141^. This thickness could be tuned by adjusting channel dimensions and flow parameters. For example, changing the inlet and outlet channel diameter from 100 μm to 40 μm while reducing the flow rate from 120 μl/h to 60 μl/h decreased the average film thickness to 40 μm (**Fig. 7e**).

**Fig. 7.**
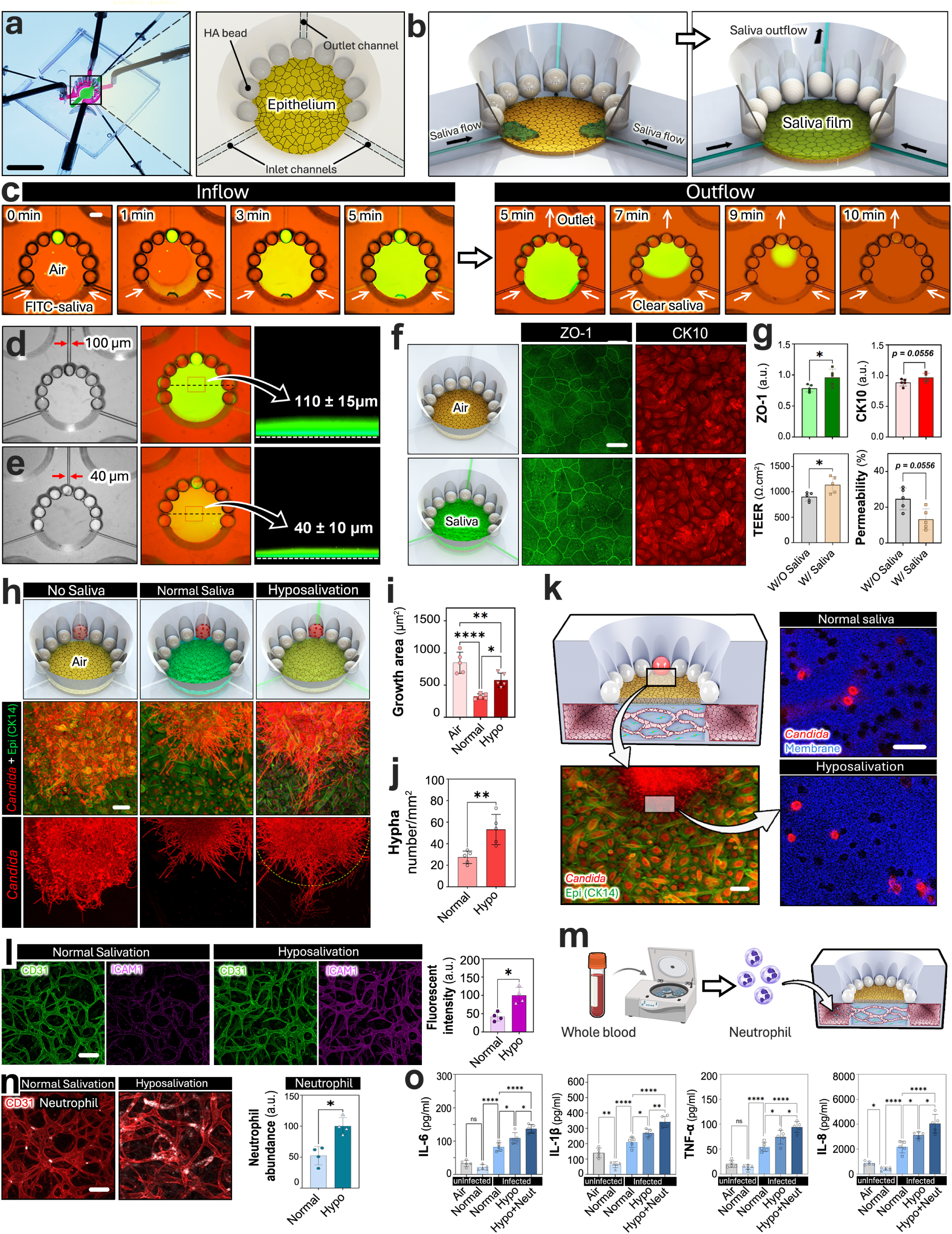
In vitro modeling of salivary flow and hyposalivation in the mouth-on-a-chip. **a,** Photograph and illustration of the mouth-on-a-chip device equipped with inlet and outlet flow channels in the epithelial chamber. Scale bar,1 cm. **b,** Schematic illustration of saliva delivery into the epithelial chamber to form a saliva film on the epithelium. **c,** Visualization of fluorescently labeled saliva entering from two lateral inlets (0-5 minutes) and clearance through the outlet channel (5-10 minutes). Scale bars, 500 µm. **d,e,** Demonstration of changing saliva film thickness to model conditions of normal salivation (**d**) and hyposalivation (**e**). Fluorescently labeled saliva was used for visualization. The thickness val-ues in the cross-sectional images represent mean±S.D. (n = 3). **f,** Immunofluorescence staining of epithelial ZO-1 and CK10 expression with and without saliva. Scale bars, 50 μm. **g,** Quantification of ZO-1 and CK10 expression and barrier function. **h,** Visualization of *C. albicans* growth under no-saliva, normal salivation, and hyposalivation conditions at 25 hours post-infection. Scale bar, 50 μm. **i,j,** Quantification of *C. albicans* growth area (**i**) and hyphal density (**j**). **k,** Immunofluorescence imaging and comparison of hyphal penetration through the porous membrane. Scale bars,100 μm (k1) and 10 μm (k3). **l**, Immunofluorescence staining of endothelial CD31 and ICAM-1 and quantification of ICAM-1 fluorescence intensity at 25 hours post-infection. Scale bar, 100 μm. **m,** Workflow for neutrophil isolation from whole blood and introduction into the vascular chamber of the mouth-on-a-chip. **n,** Representative images and quantification of neutrophil recruitment in *C. albicans-*infected tissue. Scale bar, 100 μm. **o,** Proinflammatory cytokine concentrations in epithelial effluent. Data are presented as mean ± SD, with n = 3–5 biological replicates per condition. Statistical significance was determined by two-tailed unpaired Student’s *t*-test for two-group comparisons and one-way ANOVA with Tukey’s multiple-comparisons test for comparisons among three or more groups; ns, not significant; *P < 0.05, **P < 0.01, ***P < 0.001 and ****P < 0.0001.

Notably, saliva produced measurable effects on epithelial phenotype in our model. The differentiated stratified gingival epithelium maintained under the salivary film for 3 days showed significantly upregulated ZO-1 expression compared to the ALI control without saliva, while CK10 showed a trend toward increased expression (**Figs. 7f, 7g**). Consistent with these results indicating enhanced junctional maturation and epithelial differentiation, the saliva-containing model was characterized by higher electrical resistance and lower barrier permeability (**Fig. 7g**).

Using this system, we then investigated how saliva modulates gingival responses to fungal infection with *C. albicans*. Compared with air-exposed conditions, inclusion of saliva resulted in significantly decreased fungal surface coverage and overall epithelial colonization (**Figs. 7h, 7i**). However, when the saliva layer thickness was reduced to 40 μm to simulate the thinning of the salivary film associated with decreased saliva production in the human mouth^142^, *C. albicans* spread more extensively across the epithelial surface (**Figs. 7h, 7i, 7j**) and exhibited increased invasiveness, as indicated by higher numbers of hyphae traversing into the subepithelial space (**Fig. 7k**).

Relative to the normal salivation model, hyposalivation also led to increased endothelial activation, as evidenced by significantly elevated ICAM-1 expression (**Fig. 7l**). Consistent with this pro-inflammatory phenotype, TNF-α, IL-1β, IL-6, and IL-8 were detected in significantly higher levels in epithelial effluent collected from the hyposalivation model (**Fig. 7m**). To incorporate an immune component, we perfused the stromal vasculature with primary human neutrophils isolated from the peripheral blood to create a more physiologically relevant representation of gingival infection. Under hyposalivation, neutrophil adhesion to the vascular endothelium increased significantly compared to the normal salivation condition (**Fig. 7n**), which was accompanied by further upregulation of IL-1β, IL-6, and IL-8 in epithelial effluent samples (**Fig. 7o**).

Taken together, these results support the established protective role of saliva against fungal infection^137^ and provide in vitro evidence that the altered salivary microenvironment due to hyposalivation may have direct adverse effects on fungal infection outcomes in the human oral cavity by promoting fungal growth and invasiveness, endothelial activation, immune cell infiltration, and acute inflammatory responses.

## Discussion

As one of the most common chronic conditions worldwide, biofilm-associated oral diseases are driven by microbial communities that exploit the unique anatomy and transport environment of the gingival–tooth interface. Despite their prevalence and clinical importance, mechanistic studies in laboratory settings remain challenging due to the complexity of the oral cavity in which stratified epithelium, mineralized surfaces, saliva, vascular transport, resident microbes, and immune surveillance are structurally and functionally integrated under dynamic physical and chemical conditions. The challenge of emulating this complexity motivated us to develop human-relevant microphysiological models that capture the salient features of the gingival–tooth interface under controlled conditions and enable quantitative, spatially-resolved analysis of microbial behavior and host responses. As the scarcity of physiologically relevant preclinical models is increasingly recognized as a barrier to progress in oral biology research, our work offers an experimental framework for engineering precisely controlled, human-relevant in vitro conditions to enable mechanistic investigation of gingival infection and host-microbe interactions with a degree of fidelity and analytical resolution not attainable in conventional cell culture models.

The key biological finding of our study is the emergence of a fungal infection-associated hybrid epithelial-mesenchymal state in the human gingival epithelium during *C. albicans* challenge. While bacterial pathogens such as *P. gingivalis* and *F. nucleatum* have been shown to induce EMT in gingival epithelial cells^143–144^, much less is known about fungal contributions to epithelial plasticity, and evidence has largely been limited to barrier disruption, rather than epithelial reprogramming toward a mesenchymal phenotype. *C. albicans* expresses adhesins, particularly Als3 and Hyr1, that bind to epithelial receptors to trigger internalization or cleavage of E-cadherin, thereby compromising adherens junctions of the oral epithelium^145–147^. *C. albicans* also secretes a peptide toxin called candidalysin that causes a loss or redistribution of E-cadherin and ZO-1 at cell-cell junctions to induce epithelial barrier dysfunction ^148^. However, direct evidence of *C. albicans*-induced EMT has not been reported. Our data reveal that *C. albicans* infection alone is sufficient to drive epithelial reprogramming characterized by the co-expression of canonical epithelial and mesenchymal markers. This transcriptional and phenotypic profile matches a hybrid epithelial-mesenchymal state commonly associated with partial EMT^149^. To our knowledge, the emergence and spatial persistence of this hybrid state have not been previously described in the context of fungal infection of the gingival epithelium.

Partial EMT and related forms of epithelial plasticity are often considered adaptive responses that help epithelial tissues withstand stress and facilitate repair, as demonstrated in the context of wound healing and tissue regeneration^149–150^. In our model, however, the outcome of this epithelial re-programming was predominantly maladaptive. Although the infection period was limited to 24 hours, the emergence of the partial EMT phenotype was accompanied by compromised barrier integrity, increased epithelial permeability, and hyphal outgrowth into the subepithelial compartment (**Fig. 2**).

These results suggest that *C. albicans* may have the capacity to trigger and amplify EMT-associated signaling in ways that rapidly disrupt epithelial homeostasis. We speculate that the high fungal burden and extensive epithelial surface engagement in our model may have induced excessive activation of EMT pathways, converting what might otherwise be a transient, protective stress response to a maladaptive state that promotes barrier failure and tissue invasion. Whether persistent fungal challenge drives sustained inflammation and remodeling of gingival tissue remains an important question for future investigation.

Such maladaptive epithelial responses may have important implications for oral disease. In oral squamous cell carcinoma, EMT and related epithelial plasticity programs are increasingly recognized as key drivers of tumor initiation, progression, metastasis, and therapy resistance^151–152^. Persistent or dysregulated EMT signaling has been shown to allows oral epithelial cells to acquire phenotypes associated with early neoplastic transformation and tumor progression, such as migratory capacity, stemness, and resistance to apoptosis^153–154^. Although our study was not designed to model oncogenic transformation, the induction of a hybrid epithelial-mesenchymal state by *C. albicans* in a human-relevant gingival tissue context raises the possibility that recurrent or poorly controlled fungal challenge could sustain epithelial plasticity programs that, in susceptible settings, may contribute to a pro-tumorigenic microenvironment. This hypothesis warrants further investigation, and the platform described here provides an experimental framework to test the links between fungal burden, epithelial state transitions, barrier disruption, and longer-term tissue remodeling

Providing mechanistic insight into these observations, our ligand-receptor interaction analysis identified MDK-centered signaling, including interactions involving syndecans (SDC) and nucleolin (NCL), as key regulators of the EMT phenotype in the infection model (**Fig. 4**). Previous studies of these pathways have focused predominantly on their role in tumor development and metastasis^155–158^, but our findings suggest that *C. albicans* infection engages signaling axes shared between infection-induced epithelial plasticity and oncogenic transformation. While in vivo relevance of this result remains to be established, further studies to validate and modulate the activity of these pathways may eventually inform the development of therapeutic strategies to prevent, delay, or reverse EMT-related processes implicated in both chronic gingival infection and oral carcinogenesis. More broadly, these findings motivate the exploration of mechanism-based intervention strategies that preserve epithelial homeostasis and limit maladaptive remodeling during gingival infection, complementing current approaches that primarily target microbial burden or broadly suppress inflammation.

The metabolic signatures identified in the infection model (**Fig. 5**) represent another notable outcome of our study with the potential for clinical translation. Many of the top-ranked metabolic pathways and metabolites revealed by this analysis extend beyond those commonly evaluated in periodontal research, prompting the question of whether they may serve as predictive biomarkers of infection-induced EMT and related pathological changes in the human gingiva. Our analysis was performed using effluent samples collected in a compartment-specific manner, distinguishing metabolites released on the epithelial side from those on the vascular side. This spatial resolution provides unique opportunities to compare our in vitro data to clinical measurements of metabolites and signaling factors in saliva or blood. With growing interest in saliva as a non-invasive diagnostic biofluid for monitoring oral and systemic health ^159–160^, such studies may facilitate the development of metabolite-based clinical biomarkers indicative of gingival barrier dysfunction, the severity of infection, or even early signs of malignant transformation in susceptible individuals.

Our study also demonstrated the feasibility of using the mouth-on-a-chip system to model viral infection of the human gingiva with SARS-CoV-2 (**Fig. 6**). Although the experimental burden of live-virus work limited the depth of our analysis, this proof-of-concept lays the groundwork for future research into whether and how viral infection contributes to periodontal pathology. Emerging clinical evidence shows the association of COVID-19 with exacerbated gingival inflammation, dysbiosis of the oral microbiota, and worsened damage of periodontal tissues^161–163^. While systemic immune dysregulation and local amplification of proinflammatory signals during infection are thought to play a critical role in these adverse outcomes^164^, their underlying mechanisms remain poorly understood. Our model may help address this knowledge gap by providing a precisely controlled, human-relevant platform to investigate, for example, whether viral infection primes the gingiva for secondary bacterial or fungal colonization that leads to sustained inflammation, whether it promotes EMT-like epithelial remodeling, how it alters epithelial-immune and host-microbial interactions, and so on. Such mechanistic studies may allow us to better understand the contribution of oral SARS-CoV-2 infection to periodontal biology and potentially inform the development of targeted therapies to prevent or mitigate infection-induced gingival dysfunction and post-COVID periodontal complications^165–166^.

To advance our proof-of-concept infection model into rigorous, clinically relevant investigation, it is essential to recognize that our laboratory system was designed to reproduce select features of the native oral environment, and that its utility will depend on continued improvements in biological and physiological realism. Among the most critical issues is the limited capacity of our model to capture the complexity of the oral microbiota in vivo. Although co-culture of human gingival tissues with *S. mutans* and *C. albicans* provides a controllable experimental platform for dissecting specific host-microbial interactions, generating reliable, predictive preclinical data will require increased microbial diversity, and ultimately, reconstitution of stable host-microbiome homeostasis and its perturbation toward dysbiosis. Despite advances in mammalian tissue culture and microbial cultivation, maintaining complex communities in long-term co-culture under physiologically relevant conditions remains a major challenge. Therefore, advancing our technology will require identifying biochemical and biophysical signals that stabilize oral symbiosis and implementing these regulatory features in vitro.

One promising direction for future investigation is to reproduce physiological environmental cues that shape microbial behavior and host responses in vivo, and to test whether they can be leveraged to stabilize host-microbe co-culture and suppress pathogenic microbial activity. The oral cavity is among the most mechanically active organs in which cells are continuously exposed to physical forces generated by mastication, salivary flow, and facial movements during breathing and speaking. Although work remains to be done to better understand the effects of these forces on oral tissues, studies using gut-on-a-chip models have shown that mechanical forces mimicking those produced by peristaltic motion and luminal flow contribute to establishing stable co-culture of human intestinal epithelial tissues with microbial cells^167–168^. Consistent with this finding, our saliva-containing model showed reduced fungal virulence and overgrowth (**Fig. 7**), which may be attributed to the convective motion of saliva, saliva-derived antimicrobial activity, or a combination of both. Future studies should systematically evaluate additional environmental variables present in the oral cavity, including oxygen gradients, mucins, enzymes, and antimicrobial proteins (e.g., lactoferrin, defensins). These efforts will be instrumental not only for improving co-culture stability but also for in vitro investigation of whether and how perturbations in the local microenvironment contribute to dysbiosis and pathological host responses.

Another important limitation of the current mouth-on-a-chip model is the absence of immune components. In vivo, the gingival mucosa is constantly patrolled by both resident and recruited immune cells, which play a critical role in maintaining microbial balance and coordinating inflammatory and repair responses^169–170^. While we demonstrated the feasibility of simulating neutrophil recruitment in the infection model (**Fig. 7**), this experiment is an incomplete representation of the full spectrum of host defense, which encompasses immune surveillance, pathogen clearance, and resolution of inflammation. The lack of the fully reconstituted immune microenvironment also limits mechanistic interrogation of how immune dysregulation contributes to periodontal immunopathology. Therefore, incorporating additional tissue-resident and circulating immune cell populations with controllable recruitment and activation cues represents an important next step towards improving the physiological realism of the model and its capacity to generate predictive preclinical data.

In conclusion, this work demonstrates the possibility of emulating the integrated physiological context of the human oral cavity in a microengineered 3D culture system to generate human-relevant, high-dimensional preclinical data that capture important aspects of host-microbe interactions at the gingival interface. With rapidly growing efforts to establish and adopt new approach methodologies (NAMs) across life science research, we believe that the mouth-on-a-chip platform described here will contribute to advancing in vitro technologies for mechanistic discovery and preclinical evaluation of preventive and therapeutic strategies in oral biology and dental medicine.

## Materials and Methods

### Device Fabrication

Mouth-on-a-chip devices were fabricated using standard soft lithography techniques (**Supplementary Fig. 1**). A prepolymer mixture of poly(dimethylsiloxane) (PDMS; Sylgard 184, Dow Corning, USA) was prepared by combining the monomer base and curing agent at a 10:1 (w/w) ratio. The mixture was poured into 3D-printed molds (Protolabs, USA), degassed under vacuum for 1 hour to remove air bubbles, and cured in an incubator at 65 °C for 4 hours. After curing, the PDMS components were demolded. To assemble the chip, the top PDMS structure was bonded to a thin intermediate PDMS layer (10:3 ratio) that had been spin-coated onto a polystyrene disc at 2,500 rpm for 5 minutes. A porous polystyrene membrane was affixed at the interface. The bottom PDMS layer was then aligned and sealed to complete the assembly. Fully assembled chips were incubated overnight at 65 °C to promote irreversible bonding between layers. Devices were fabricated either as individual units or in multiplexed 15-chip arrays (**Fig. 1**) and mounted in custom rectangular single-well cell culture plates for downstream experiments.

### Production of vascularized stroma

Prior to cell seeding, all devices were sterilized by ultraviolet (UV) exposure for 30 minutes using a UV curing system (ELC-500, Electro-lite). To generate a perfusable vascular network within the device, 10 μl of fibrin-based hydrogel precursor solution containing human vascular endothelial progenitor cells (3.5 × 10⁶ cells/ml), human gingival fibroblasts (5.5 × 10⁶ cells/ml), thrombin (1 U/ml; T7513, Sigma-Aldrich, USA), and aprotinin (1 U/ml; A1153, Sigma-Aldrich, USA) was injected into the central vascular chamber via the designated inlet (**Fig. 1**). Devices were then incubated at 37 °C with 5% CO₂ for 15 minutes to allow for gel polymerization. Following gelation, endothelial growth medium (EGM-2, Lonza) was introduced into the side channels and media reservoirs. After 24–48 hours of initial culture, the medium was aspirated, and the side channels were incubated with fibronectin (10 μg/ml in EGM-2) for 1 hour at 37 °C to promote endothelial adhesion. The fibronectin solution was then removed, and a suspension of human vascular endothelial progenitor cells (5 × 10⁶ cells/mL in EGM-2) was introduced into the side channels. The cells were allowed to attach for 1 hour before replenishing the reservoirs with fresh medium .This seeding strategy led to the formation of a confluent endothelial monolayer along the side channels, which facilitated anastomosis with the microvascular structures formed within the fibrin gel, ultimately establishing a perfusable vascular network (**Fig. 1**).

### Production of gingival epithelium

Following vascular network formation, the upper (epithelial) compartment of the device was prepared by coating the porous polystyrene membrane with fibronectin (10 μg/ml in EGM-2 medium; F1141, Sigma-Aldrich, USA) for 1 hour at 37 °C. Primary human gingival epithelial progenitor cells (Lifeline Cell Technology, USA) at passage 2 were then seeded onto the membrane surface at a density of 3 × 10⁵ cells/ml. The cells were initially cultured in growth medium (DermaLife K, Lifeline Cell Technology) under submerged conditions for 2–3 days to allow for the formation of a confluent monolayer.

Once confluency was achieved, the growth medium was removed from the upper compartment, and a mixture of gingival epithelial differentiation medium (3D Barrier Medium, ZenBio, USA) and EGM-2 (Lonza, USA) was added to the reservoirs connected to the stromal compartment. The medium was replenished every 2 days. The epithelial tissue was cultured at an air-liquid interface for 10-12 days to promote differentiation and stratification. This process resulted in the formation of a multilayered, keratinized gingival epithelium supported by a perfusable vascular network (**Fig. 1**).

### Characterization of blood vessels

Vascular morphology was quantitatively assessed over time using Angiotool software (Angiotool version 0.6a, https://ccrod.cancer.gov/confluence/display/ROB2/Home). Metrics including total vessel length, total vessel area, number of junctions, and average vessel diameter were calculated for each device. At each time point, five independent devices were analyzed, and the results were reported as mean ± standard deviation (SD). To evaluate the perfusability of the microvascular network, medium containing either 1 μm fluorescent microspheres (FluoSpheres, F-8815, Thermo Fisher Scientific, USA) or FITC-labeled dextran (70 kDa; Sigma-Aldrich, USA) was introduced into the side channels. After removing media from all reservoirs and side channels, the two reservoirs connected to the same side channel were filled with the test solution. Hydrostatic pressure generated by the fluid height difference between inlet and outlet drove perfusion of the solution through the vascular network embedded in the fibrin matrix. Perfusion dynamics were monitored and recorded over time using a laser scanning confocal microscope (LSM 800, Carl Zeiss, Germany).

### Immunostaining of gingival epithelium

For immunofluorescence staining, medium was removed from the devices, and samples were gently washed twice with phosphate-buffered saline (PBS). Tissues were fixed with 4% paraformaldehyde (Electron Microscopy Sciences, USA) for 30 minutes at room temperature, followed by permeabilization with 0.1% Triton X-100 (Sigma-Aldrich, USA) in PBS for 15 minutes. Samples were then incubated in blocking buffer (PBS containing 3% bovine serum albumin (BSA); Sigma-Aldrich) for 1 hour at room temperature. After triple washing with PBS, samples were incubated overnight at 4 °C with primary antibodies targeting epithelial markers (cytokeratin-10/14, ZO-1, Occludin, Vimentin, Fibronectin, ACE2, and Furin) and endothelial markers (CD31, ICAM-1). On the following day, samples were washed twice with PBS and incubated overnight at 4 °C with fluorophore-conjugated secondary antibodies, including Goat anti-Mouse IgG H&L (Alexa Fluor 488 and 647), Goat anti-Rabbit IgG H&L (Alexa Fluor 555), Donkey anti-Rabbit IgG (Alexa Fluor 488), and Donkey anti-Mouse IgG (Alexa Fluor Plus 555) (all from Thermo Fisher Scientific, USA). Nuclei were counterstained using Hoechst 33342 (D1306, Thermo Fisher Scientific, USA). Stained tissues were imaged using a laser scanning confocal microscope (LSM 800, Carl Zeiss, Germany). Image acquisition and processing were performed using ZEN (Zeiss, Germany) and ImageJ software (NIH).

### Hematoxylin and Eosin (H&E) staining of gingival epithelium

For histological evaluation of gingival epithelial tissues, devices were first rinsed with cold PBS and fixed with 4% paraformaldehyde (Electron Microscopy Sciences, USA) at room temperature. Following fixation, the top and bottom PDMS layers were separated, and the polystyrene membrane containing the epithelial cell layer was carefully excised for processing. Samples were subjected to a graded ethanol dehydration series (50%, 60%, 70%, 80%, 90%, and 100%) and subsequently immersed in a xylene-based clearing agent for 5 minutes. Tissues were then transferred to embedding cassettes and infiltrated with molten paraffin. Paraffin blocks were sectioned at a thickness of 5-8 μm using a microtome, and sections were mounted onto glass slides. For staining, tissue sections were deparaffinized in clearing agent and rehydrated through a reverse ethanol gradient (100%, 90%, 80%, 70%) followed by distilled water. Samples were then incubated in a protein blocking solution for 1 hour. Hematoxylin staining was performed by immersing sections in hematoxylin solution for 1 minute, followed by three rinses in distilled water. Sections were then stained in eosin for 30 seconds, rinsed again, dehydrated through ascending ethanol concentrations, cleared, and mounted with a coverslip. Brightfield images were acquired using a standard upright microscope under consistent exposure and magnification settings.

### Analysis of epithelial barrier function

To assess epithelial barrier function, trans-epithelial electrical resistance (TEER) measurements were performed at defined time points during gingival epithelial cell differentiation. Two nickel electrodes connected to a digital ohmmeter were inserted into the apical and basolateral reservoirs of the device to record electrical resistance across the multilayered tissue. Measurements were performed on five independent devices per time point. Devices without cells were measured in parallel to account for background resistance, and net TEER values were calculated by subtracting the blank device values from total resistance. To evaluate barrier disruption in response to microbial infection, TEER was measured in each device both before and after inoculation. Changes in electrical resistance were used to quantify alterations in epithelial integrity over time.

### Preparation of HA microbeads

Hydroxyapatite (HA) microbeads (500 μm in diameter) were pre-incubated in filter-sterilized human saliva at 37 °C for 60 minutes to promote the formation of a salivary pellicle, a protein-rich adsorptive layer that mimics the biochemical characteristics of tooth enamel. This step facilitates physiologically relevant microbial attachment. *Streptococcus mutans* UA159 and *Candida albicans* SN250, two well-characterized oral pathogens, were cultured to exponential phase in ultrafiltered tryptone-yeast extract (UFTYE) medium (2.5% tryptone, 1.5% yeast extract, 1% glucose) at 37 °C with 5% CO₂. The HA microbeads were subsequently incubated with *C. albicans* (10⁵ CFU/ml; yeast form) and/or *S. mutans* (10⁷ CFU/ml) in UFTYE supplemented with 1% sucrose for 60 minutes at 37 °C under gentle rocking to enable microbial colonization. Following incubation, microbeads were washed three times with PBS to remove non-adherent cells. Fluorescently tagged strains were used to enable imaging: *C. albicans* SN250 expressing tdTomato and *S. mutan*s UA159 expressing green fluorescent protein (GFP).

### Analysis of fungal-bacterial growth

The growth dynamics of fungal-bacterial biofilms were monitored using time-lapse confocal imaging and quantitative computational analysis, as previously described^47^. Following microbial colonization, saliva-coated HA microbeads harboring *C. albicans* and *S. mutans* were aseptically transferred into the epithelial chamber of the device. Biofilm growth was supported by a medium consisting of UFTYE supplemented with 25% sterile human saliva and 1% sucrose. Three-dimensional time-lapse imaging was performed every 30 min at 37 °C using a Zeiss LSM800 confocal microscope equipped with a 40X water-immersion objective (numerical aperture = 1.2). This setup enabled high-resolution tracking of spatial and temporal biofilm development within the VMOC. Quantitative image analysis was conducted using BiofilmQ software (https://drescherlab.org/data/biofilmQ) ^171^. Biofilm volume was measured over time using the global metric “Biofilm_Volume.” Additionally, surface coverage dynamics were assessed by calculating the total area of segmented biofilm pixels in maximum intensity z-projections. These metrics provided a quantitative readout of microbial expansion and colonization behavior within the VMOC microenvironment.

### Infection with C. albicans and S. mutans

To model mono- and polymicrobial infections of the gingival epithelium, sterile HA microbeads in the device were aseptically replaced with microbeads pre-coated with *C. albicans*, *S. mutans*, or a combination of both, simulating localized microbial plaque. Following placement of the infected microbeads into designated positions, devices were returned to a humidified cell culture incubator at 37 °C with 5% CO₂. Time-lapse imaging was conducted at 5, 10, 15, 20, and 25 hours post-infection to monitor microbial growth and spatial dynamics. At each time point, five independent devices per condition were analyzed. At the 25-hour endpoint, infected tissues were fixed with 4% paraformaldehyde in PBS and stored at 4 °C for downstream histological and immunofluorescence analyses.

### Viral Infection

#### a. Pseudovirus Infection

To assess the susceptibility of gingival epithelial tissues to SARS-CoV-2 entry, a pseudovirus-based infection assay was performed. Gingival epithelial cells were seeded at a density of 3.5 X 10⁵ cells/cm² and cultured under submerged conditions for 1 day, followed by air-liquid interface (ALI) conditions for up to 12 days. Pseudovirus infection was conducted at three time points representing distinct tissue maturity stages: day 1 (submerged), day 6 (ALI), and day 12 (ALI). At each time point, 50 μl of infection medium (comprising 46.9 μl of epithelial growth medium, 2.5 μl of SARS-CoV-2 pseudovirus stock (Montana Molecular, Cat. No. C1110G), and 0.6 μl of sodium butyrate) was added to the epithelial chamber of the device. Infected devices were incubated at 37 °C with 5% CO₂ for 24 hours. After infection, cells were washed three times with PBS and fixed with 4% paraformaldehyde in PBS. Infected cells expressing green fluorescent protein (GFP) in the nucleus were identified according to the manufacturer’s specifications. Fluorescence imaging was conducted using a laser scanning confocal microscope (LSM 800, Carl Zeiss, Germany), and image processing was performed using ZEN software (Zeiss) and ImageJ (NIH).

#### **b.** Wild-type SARS-CoV-2 Infection

To investigate host-virus interactions with infectious SARS-CoV-2, fully differentiated and vascularized gingival epithelial tissues in our model were infected after 12 days of ALI culture. Infection was carried out at a multiplicity of infection (MOI) of 0.3. The apical surface was washed three times with gingival epithelial differentiation medium, followed by the addition of 50 μl of SARS-CoV-2 inoculum for 1 hour. After incubation, the inoculum was removed, and fresh medium was added to the basolateral reservoirs. This infection regimen was repeated three times over a 72-hour period to enhance infection efficiency. Total RNA was extracted from epithelial tissues using TRIzol reagent (Invitrogen, USA), and cDNA was synthesized using the iScript cDNA Synthesis Kit (Bio-Rad, USA) according to the manufacturer’s protocol. Quantitative RT-PCR was performed using TaqMan® Gene Expression Assays to quantify the mRNA expression levels of proinflammatory cytokines and chemokines, including IL6, TNFA, IL8, IL1B, CXCL10, IFNA1, IFNB1, and IFNG, which are associated with host antiviral response and COVID-19 pathogenesis.

### Modeling hyposalivation

To model hyposalivation in the mouth-on-a-chip, modified version of the device was engineered to include three additional microchannels for saliva flow regulation (**Fig. 7a**). One channel served as the saliva outlet, while the other two were configured as inlets. In devices mimicking normal salivary flow, these channels had a diameter of 100 μm; in the hyposalivation condition, the diameter was reduced to 40 μm. Saliva was perfused through the epithelial compartment at a flow rate of 120 μl/h for the normal condition and 60 μl/h for the hyposalivation model. This configuration resulted in a stable saliva film with an average thickness of approximately 110 ± 15 μm under normal flow and 40 ± 10 μm under hyposalivation.

To examine the impact of salivary flow on *C. albicans* colonization and invasion, pooled sterile human saliva (Innovative Research, USA) was introduced into the apical chamber. After establishing flow conditions, a control (saliva-only) microbead was replaced with a *Candida*-infected microbead as described previously (**Fig. 2a**). Devices were maintained in a humidified cell culture incubator at 37 °C with 5% CO₂ for 24 hours. At the endpoint, saliva and medium were collected from the apical and basolateral compartments and stored at −80 °C for further biochemical analysis. Devices were then washed three times with PBS, fixed in 4% paraformaldehyde, and stored at 4 °C for subsequent imaging. To monitor fungal growth and epithelial colonization, a tdTomato-expressing *C. albicans* SN250 strain was used. Gingival epithelial cells were stained with cytokeratin 14 antibody (CK14; Abcam, USA) and Hoechst 33342 (Thermo Fisher Scientific, USA). Fluorescence imaging was performed using an upright AxioZoom microscope (Zeiss, Germany), and *Candida* coverage area was quantified using ImageJ software (NIH, USA).

To evaluate fungal invasion across the epithelial tissue, five independent regions per sample were imaged to capture *Candida*-colonized zones. Imaging was focused on the epithelial-membrane interface to visualize cross-sections of hyphae penetrating through the porous membrane. The number of hyphae traversing from the epithelial to the vascular compartment per unit area was defined as the invasion index.

### Neutrophil infiltration assay

Human peripheral blood neutrophils were isolated from apheresis products obtained through the Human Immunology Core at the University of Pennsylvania. To examine neutrophil recruitment under microbial infection conditions, neutrophils were labeled with CellTracker™ Deep Red (Thermo Fisher Scientific, USA) and resuspended in culture medium at a final concentration of 2 × 10⁶ cells/ml. The labeled cell suspension was introduced into the vascular compartment via one of the sides microchannels and allowed to circulate through the microvascular network for 4 hours at 37 °C in a humidified cell culture incubator. Following perfusion, devices were gently washed three times with fresh culture medium to remove non-adherent cells. Neutrophil adhesion patterns were assessed qualitatively by fluorescence imaging and visually compared between healthy and infected conditions.

### Analysis of proinflammatory cytokines and chemokines

To quantify the secretion of proinflammatory cytokines and matrix remodeling enzymes by vascularized gingival epithelial tissues in our model, enzyme-linked immunosorbent assays (ELISAs) were performed on effluent media collected under healthy and infection conditions (*C. albicans*, *S. mutans*, and *Ca*+*Sm* co-infection). Five independent devices were prepared per condition. Medium samples were collected from both the epithelial and vascular chambers prior to infection and 24 hours post-infection. Collected media were centrifuged at 2,000 × *g* for 15 minutes to remove cellular or extracellular matrix debris and stored at −80 °C until analysis. The following commercially available ELISA kits were used according to manufacturers’ instructions: IL-6(Sigma-Aldrich, RAB0306), TNF-α (Abcam, ab181421), IL-1β (Sigma-Aldrich, RAB0273), IL-8/CXCL8 (Sigma-Aldrich, RAB0319), IL-18 (Sigma-Aldrich, RAB0543-1KT), MMP-2 (Sigma-Aldrich, RAB0365-1KT), LDH (cytotoxicity control) (Roche, 04744926001).

Briefly, 100 μl of either standard solution or conditioned medium was added to each well. After a 2-hour incubation at room temperature, wells were washed five times with 300 μl of manufacturer-supplied wash buffer and incubated with the respective biotinylated detection antibody for 1 hour.

Wells were then washed and incubated with 100 μl of TMB substrate for 20 minutes in the dark. Reactions were stopped with 100 μl of stop solution, and optical density (O.D.) was measured using a plate reader (M200, Tecan, Switzerland). Cytokine concentrations were calculated from standard curves generated by plotting O.D. versus known concentrations of standards.

### Single-cell RNA sequencing

For single-cell transcriptomic analysis, devices were first washed three times with PBS to remove residual media. Gingival epithelial and stromal compartments were enzymatically dissociated using trypsin, and the resulting cell suspension was filtered through a 30 μm cell strainer to eliminate matrix debris and cellular aggregates. Live cells were resuspended in DMEM containing 5% fetal bovine serum (FBS) at a final density of 800 live cells/μl. Single-cell suspensions from each sample were loaded into individual channels of a Chromium Single Cell 3′ Reagent Kit v2 microfluidic chip (10X Genomics, USA) and processed according to the manufacturer’s protocol. During gel bead-in-emulsion (GEM) generation, mRNA transcripts were uniquely barcoded and reverse transcribed. Complementary DNA (cDNA) was then amplified, and sequencing libraries were prepared following the 10X Genomics user guide. Libraries were sequenced on an Illumina NovaSeq 6000 platform using an S1 100-cycle flow cell (v1.5 chemistry). Library quality was assessed using the Agilent TapeStation (for fragment size distribution) and KAPA Library Quantification Kit (for concentration). Raw sequencing reads were de-multiplexed and aligned to the GRCh38 human transcriptome using the Cell Ranger pipeline (v5.0.0, 10X Genomics). Data from individual libraries were aggregated and normalized for sequencing depth using the Cell Ranger `“aggr”` function prior to downstream analysis.

### Single-cell sequencing data analysis

To analyze single-cell RNA sequencing data, the Seurat package (v5.0.1) was used. Low-quality cells were initially filtered out based on gene and mitochondrial content thresholds. Raw gene count data were normalized to account for sequencing depth variation and scaled for uniformity. Highly variable genes were identified to guide downstream analysis. For data integration across experimental conditions, the `merge()` function was applied. Dimensionality reduction was performed using Latent Semantic Indexing (LSI), and hierarchical clustering was followed by visualization with Uniform Manifold Approximation and Projection (UMAP). Cells expressing fewer than 200 genes or more than 20% mitochondrial transcripts were excluded from further analysis. Integration anchors across conditions were calculated using `FindIntegrationAnchors(),` and datasets were integrated using `IntegrateData()`. Linear dimensionality reduction was performed with principal component analysis (PCA), and the number of significant principal components was determined using the `ElbowPlot()` function. Cells were clustered using a shared nearest neighbor (SNN)-based algorithm. Clusters were visualized in two-dimensional space using UMAP. Differential expression analysis was performed using the `FindAllMarkers()` function. Genes were considered differentially expressed if they had a log2 fold change > 0.25, were expressed in at least 25% of cells in the compared groups, and had an adjusted p-value < 0.05. Clusters were annotated based on the expression of top differentially expressed genes (DEGs) and canonical cell type markers.

### Pseudotime and trajectory analysis

Monocle 3 (v1.0.0) was employed to investigate the temporal progression of cellular transcriptional states through pseudotime analysis. Cells were ordered based on transcriptional similarity to reflect their advancement along biological processes. Epithelial cell clusters were extracted from the integrated Seurat object and converted into a Monocle-compatible format. Pseudotime trajectories were constructed using principal graph learning. The root node of the gingival epithelial cell trajectory was defined based on cells expressing the highest levels of basal epithelial markers (*TP63*, *KRT15*). Gene expression dynamics along the trajectory were assessed using Moran’s I test to identify genes whose expression varied significantly with pseudotime. Genes exhibiting consistent temporal trends, including monotonic increases, decreases, or biphasic patterns, were identified and prioritized for down-stream analysis.

### Gene set enrichment analysis

Gene set enrichment analysis (GSEA) was performed to identify biological pathways enriched in gingival epithelial cells from the *Ca+Sm* infected model compared to healthy controls. Differentially expressed genes were identified from single-cell RNA-seq data and ranked by log₂ fold change. Ranked gene lists were analyzed using the clusterProfiler R package (v4.6.0) to evaluate enrichment across three gene set collections from the Molecular Signatures Database (Mysid v7.5.1): **Hallmark**, **KEGG**, and **Reactome** pathways. Enrichment scores were computed using the `GSEA()` function with 1,000 permutations. Pathways with a false discovery rate (FDR) q-value < 0.05 were considered statistically significant. Normalized enrichment scores (NES) were used to rank pathways, and key signaling cascades associated with epithelial-to-mesenchymal transition (EMT) were further analyzed. Visualization of NES rankings and pathway components was performed using the `ggplot2` and `enrichplot` packages in R.

### Analysis of ligand-receptor interactions

To elucidate enriched ligand-receptor interactions mediating soluble or contact-dependent inter-lineage signaling within the *Ca+Sm* infected model, we employed **CellChatDB v2.079**^68^. This manually curated database of ligand-receptor pairs is integrated with a statistical framework designed to infer cell-cell communication networks from single-cell transcriptomic data.

Normalized count matrices from *Ca+Sm* infected samples were used as input, along with cell identity metadata derived from scRNA-seq analysis. Potential interactions between cell types were quantified by computing the mean of the average expression levels of ligands and their cognate receptors. Statistical significance was assessed using a permutation-based test, in which cell type labels were randomly shuffled 1,000 times to generate a null distribution. Interactions with mean expression levels greater than 0.1 and permutation *p*-values below 0.05 were considered significant. To focus on intercellular communication, direct integrin-extracellular matrix interactions were excluded. Interaction networks were visualized using the **Circlize** R package. Global interaction strength between cell types was represented using aggregate chord diagrams, while directional chord diagrams highlighted individual ligand-receptor interactions with a total mean expression above 0.35. In directional diagrams, each chord reflects the maximum of all significant means across interacting cluster pairs, with chord width corresponding to the overall interaction strength for a given ligand-receptor pair.

### Metabolomic analysis

To assess infection-induced changes in metabolic activity of vascularized gingival tissue, media from the apical and basolateral chambers of each device were collected 24 hours after infection with *C. albicans* (*Ca*), *S. mutans* (*Sm*), or the *Ca+Sm* combination. A total of 50 μl of medium was harvested per chamber and stored at −80 °C until processing. Metabolite extraction was initiated by adding 5 μl of each sample to 120 μl of pre-chilled (−20 °C) solvent mixture (25:25:10, v/v/v acetonitrile:methanol:water), followed by vertexing for 10 seconds and incubation on ice for at least 5 minutes. Samples were centrifuged at 16,000 × *g* for 20 minutes at 4 °C, and the supernatant was transferred to new tubes for LC–MS analysis. A procedure blank (solvent only) was processed in parallel to identify and subtract background signals.

Metabolites were analyzed using a Vanquish Horizon UHPLC system (Thermo Fisher Scientific, USA) coupled to an Orbitrap Exploris 480 mass spectrometer (Thermo Fisher Scientific, USA). Separation was performed using a Waters XBridge BEH Amide XP column (2.5 μm, 150 mm X 2.1 mm) under hydrophilic interaction chromatography (HILIC) conditions at 25 °C. Mobile phase A consisted of 20 mM ammonium acetate and 22.5 mM ammonium hydroxide in 95:5 (v/v) water:acetonitrile (pH 9.45), and mobile phase B was 100% acetonitrile. Samples were analyzed in both positive and negative ESI modes using a linear gradient elution profile, with a total runtime of 25 minutes and a flow rate of 0.15 ml/min. The injection volume was 5 μl.

ESI source parameters were as follows: spray voltage, 3200 V (positive) or −2800 V (negative); sheath gas, 35 arb; auxiliary gas, 10 arb; sweep gas, 0.5 arb; ion transfer tube temperature, 300 °C; vaporizer temperature, 35 °C. LC–MS data were acquired in full scan polarity-switching mode with an Orbitrap resolution of 120,000 at *m/z* 200, AGC target of 1e7, maximum injection time of 200 ms, and scan range of *m/z* 60–1000.

Raw LC–MS files (.raw) were converted to mzXML format using ProteoWizard (v3.0.20315). Peak detection was performed in El-MAVEN (v0.12.0) with modified parameters: mass resolution, 5 ppm; time domain resolution, 10 scans; minimum peak intensity, 10,000; and minimum peak width, 5 scans. The resulting peak table was exported as a .csv file. Peak annotation for untargeted metabolomics was conducted using NetID with default parameters. Statistical analysis was performed using MetaboAnalyst 5.0, including data normalization, multivariate analysis, and pathway enrichment analysis.

### Statistical analysis

Sample sizes were determined based on *n* = 3-5 independent devices per experimental group. Data was analyzed using GraphPad Prism 10 (GraphPad Software, USA). For comparisons between two groups, Student’s *t*-test was used. For comparisons involving more than two groups, one-way ANOVA was performed followed by appropriate post hoc tests. Results are presented as mean ± SD. Statistical significance was defined as follows: \**P* < 0.05*, **P< 0.01,***P < 0.001*, \*\*\*\**P* < 0.0001.

## Acknowledgements

This work was supported by the National Institute of Dental and Craniofacial Research (NIDCR) (T90-DE-030854). M.Y. and Z.R. were supported by the NIDCR T90/R90 Mentored Scientist Training Award from the Center for Innovation and Precision Dentistry at the University of Pennsylvania and the NIDCR Postdoctoral Training Program (R90DE031532 to H.K.), respectively. Z.R. is supported by the National Institute of Dental & Craniofacial Research (NIDCR) of the National Institutes of Health under Award Number 1K99DE033428.

## Author Contributions

M.Y. designed the research, performed the experiments, wrote the manuscript. M.Y and P.F. analyzed the data and performed bioinformatics analyses. M. K. and R.Z. designed the research and performed the experiments. In collaboration with W. D. L. and J. D. R. developed and executed the metabolomic analysis at the Princton University. D.D.H. and H.K. designed the research, analyzed the data, and wrote the manuscript.

---

**Extended Data Fig. 1.**
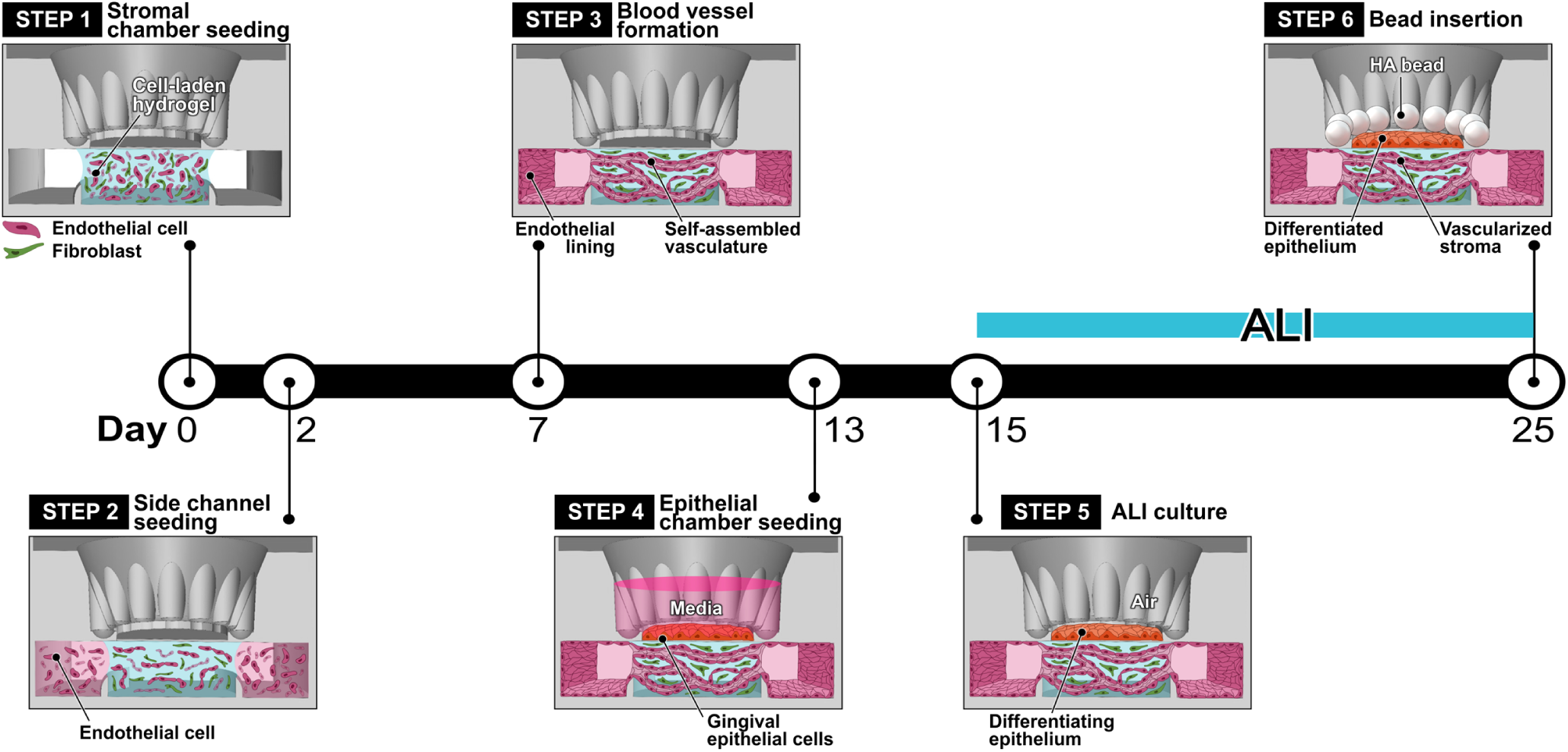
Timeline for model production. A human gingival tissue construct is generated through 13 days of endothelial-fibroblast co-culture in the stromal compartment and the resultant vasculogenic self-assembly of blood vessels, followed by submerged and ALI culture of gingival epithelial cells for additional 12 days.

**Extended Data Fig. 2.**
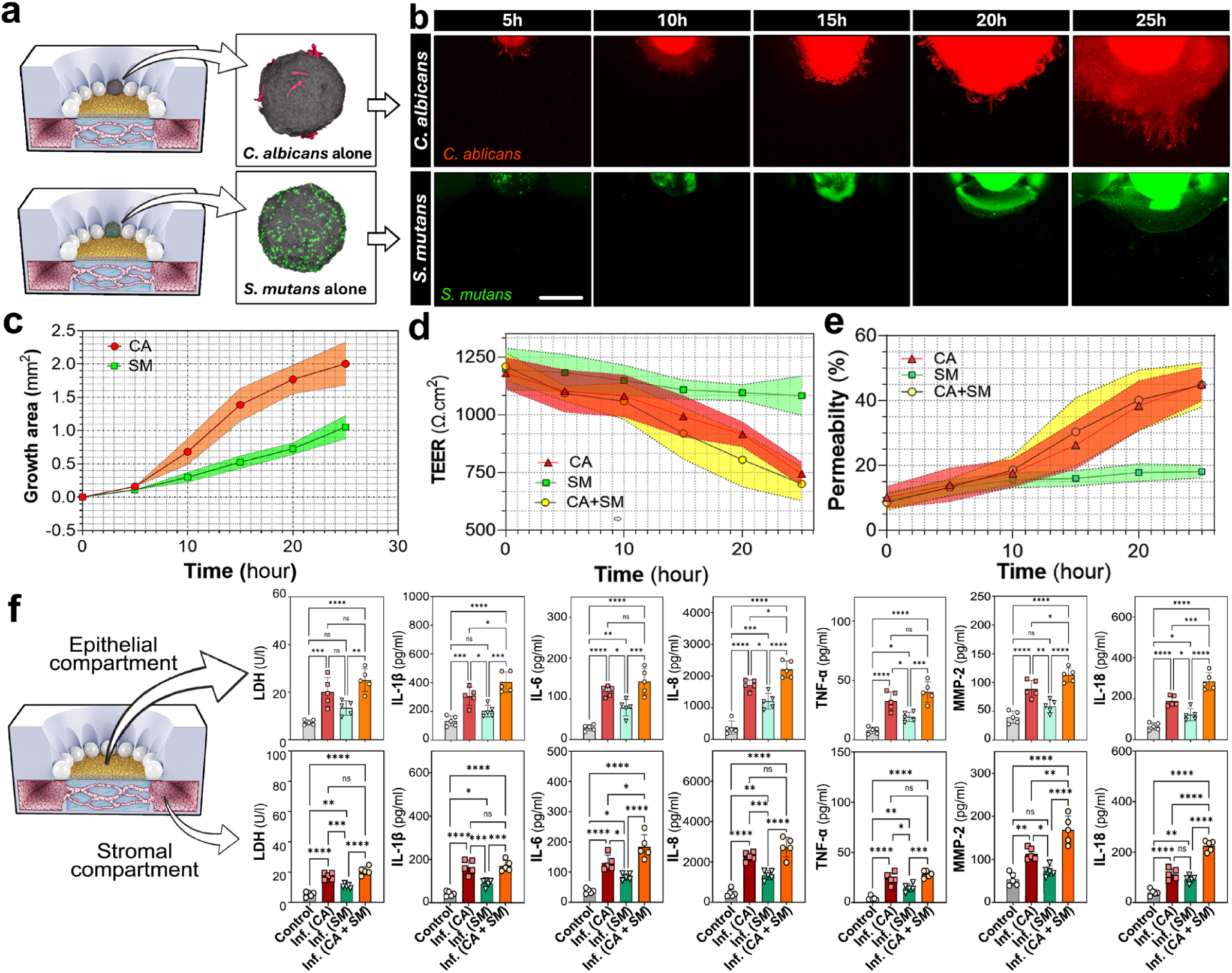
Single-species infection. **a,** Micrographs of HA beads coated with either *C. albicans* (red) or *S. mutans* (green) used in the single-species infection experiments. **b,** Representative time-lapse fluorescence images of microbial growth over the epithelium over 25 hours. Scale bar, 500 µm. **c,d,e,** Quantification and comparison of microbial growth area (**c**), transepithelial electrical resistance (TEER; **d**), barrier permeability to 4 kDa-FITC dextran (**e**) over time in response to *C. albicans* (*CA*), *S. mutans* (*SM*), or mixed *C. albicans* + *S. mutans* infection (*CA+SM*). **f,** Quantification of LDH and inflammatory cytokine levels in device effluent collected from the epithelial (top row) and stromal (bottom row) compartment. Data are presented as mean ± SD. Individual points represent independent devices or biological replicates, with n = 5 per condition. Statistical significance was determined by two-tailed unpaired Student’s *t*-test for two-group comparisons and one-way ANOVA with Tukey’s multiple-comparisons test for comparisons among three or more groups. ns, not significant; *P < 0.05, **P < 0.01, ***P < 0.001 and ****P < 0.0001.

