## Supplementary material for "Emulating the gingival-tooth interface during bacterial, fungal, and viral infection in a microphysiological model of the human oral cavity": Supplmentary Information

**Supplementary Information includes:**

**Supplementary Figures 1 to 15:**
**Supplementary Figure 1.** Fabrication workflow and dimensions of mouth-on-a-chip device.
**Supplementary Figure 2.** Effect of ALI culture on epithelial barrier function.
**Supplementary Figure 3.** Epithelial markers for cell-type annotation.
**Supplementary Figure 4.** Analysis of ligand-receptor pairs mediating intercellular interactions.
**Supplementary Figure 5.** Plot of partial least squares discriminant analysis (PLS-DA) of metabolites in the epithelial compartment.
**Supplementary Figure 6.** Heatmap of epithelial metabolites.
**Supplementary Figure 7.** Plot of partial least squares discriminant analysis (PLS-DA) of metabolites in the vascular/stromal epithelial compartment.
**Supplementary Figure 8.** Heatmap of stromal metabolites.
**Supplementary Figure 9.** Top-ranked differentially regulated epithelial metabolites.
**Supplementary Figure 10.** Top-ranked differentially regulated stromal metabolites.
**Supplementary Figure 11**. Metabolite biomarker analysis of the infected gingival epithelium.
**Supplementary Figure 12.** Metabolite biomarker analysis of the infected gingival stroma.
**Supplementary Figure 13**. Top five discriminatory epithelial metabolites.
**Supplementary Figure 14**. Top five discriminatory stromal metabolites.
**Supplementary Figure 15.** Device design to model salivary flow.

**Supplementary Tables:**

**Supplementary Table 1.** Key resources
**Supplementary Table 2.** List of primers

**Supplementary Fig. 1 | Fabrication workflow and dimensions of mouth-on-a-chip device.**

**a,** Schematic showing the process of fabricating micropatterned PDMS layers to build a mouth-on-a-chip device. **b,** Device illustrations showing the overall device footprint, microchannel geometry, and the dimensions of key structural components.

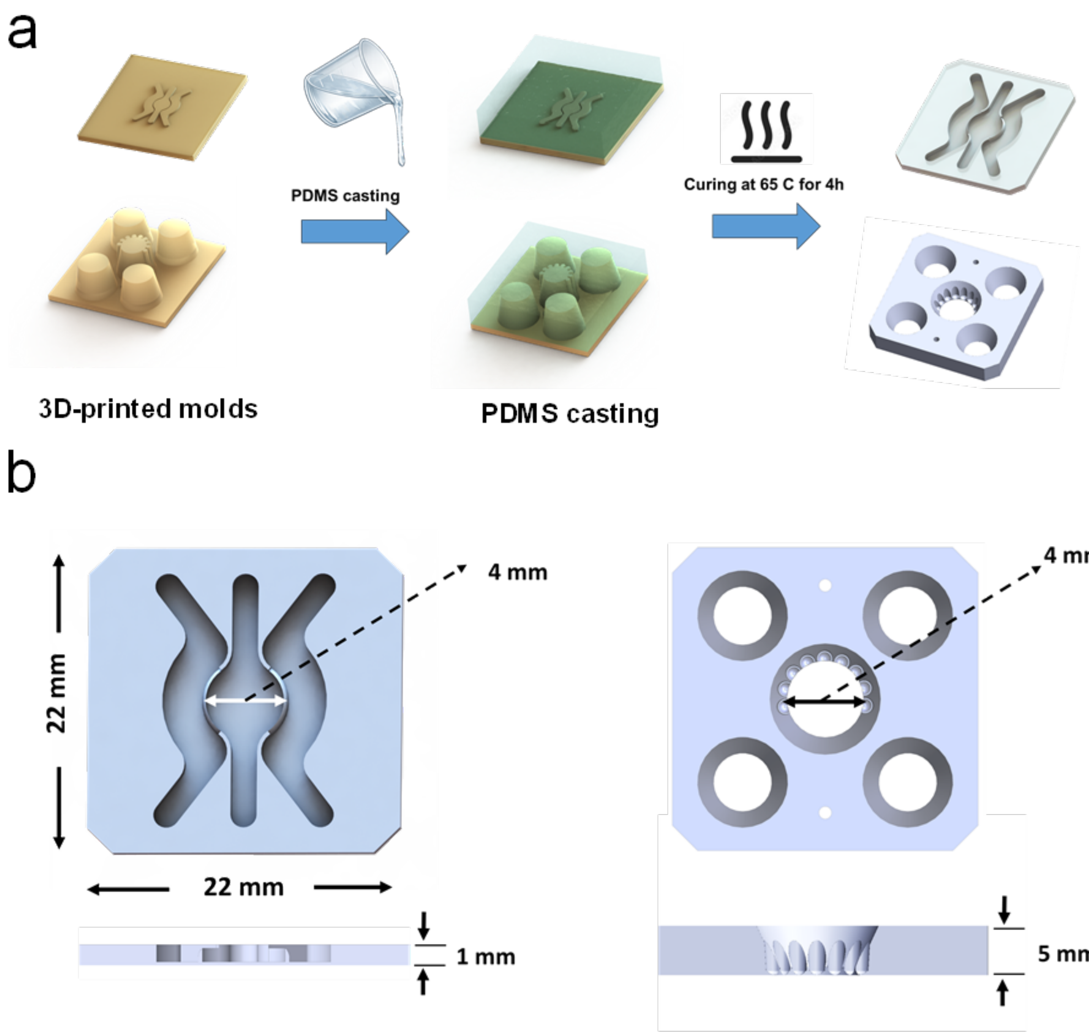

### Supplementary Figure 2. Effect of ALI culture on epithelial barrier function

Time-dependent decrease in epithelial permeability and corresponding increase in transepithelial electrical resistance (TEER) after ALI exposure, demonstrating progressive epithelial barrier maturation in the MOC device. Data are presented as mean ± S.D.

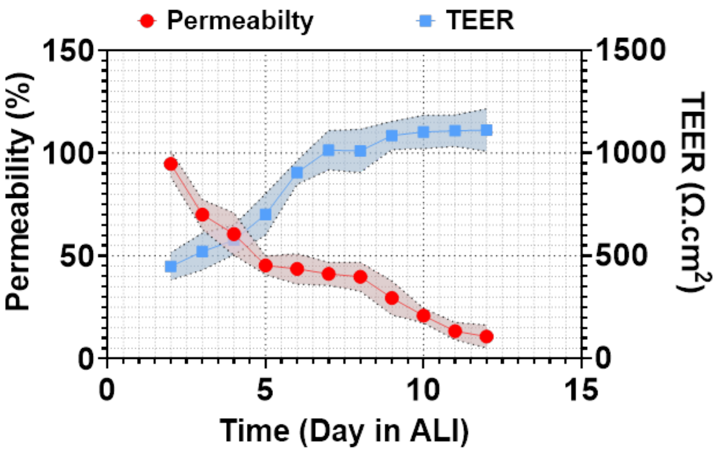


### Supplementary Figure 3. Epithelial markers for cell-type annotation

Dot plot showing the expression of i) canonical epithelial differentiation markers for basal, spinous, granular, superficial epithelial cells and ii) stromal markers for endothelial cells and fibroblasts. These markers were used for identification and annotation of cell types shown in Figure 1i. Dot size represents the fraction of cells expressing each gene, and color intensity indicates average scaled expression within each cell type.

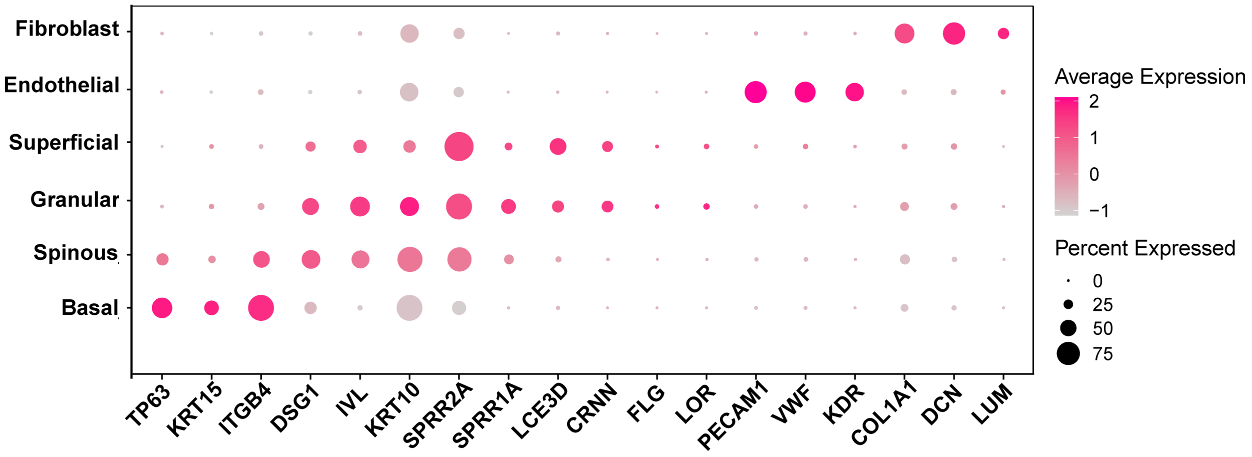

### Supplementary Figure 4. Analysis of ligand-receptor pairs mediating intercellular interactions

Bar plot showing the relative contribution of individual ligand–receptor (LR) pairs to cell–cell communication in the infected model. LR pairs are ranked by their contribution score.

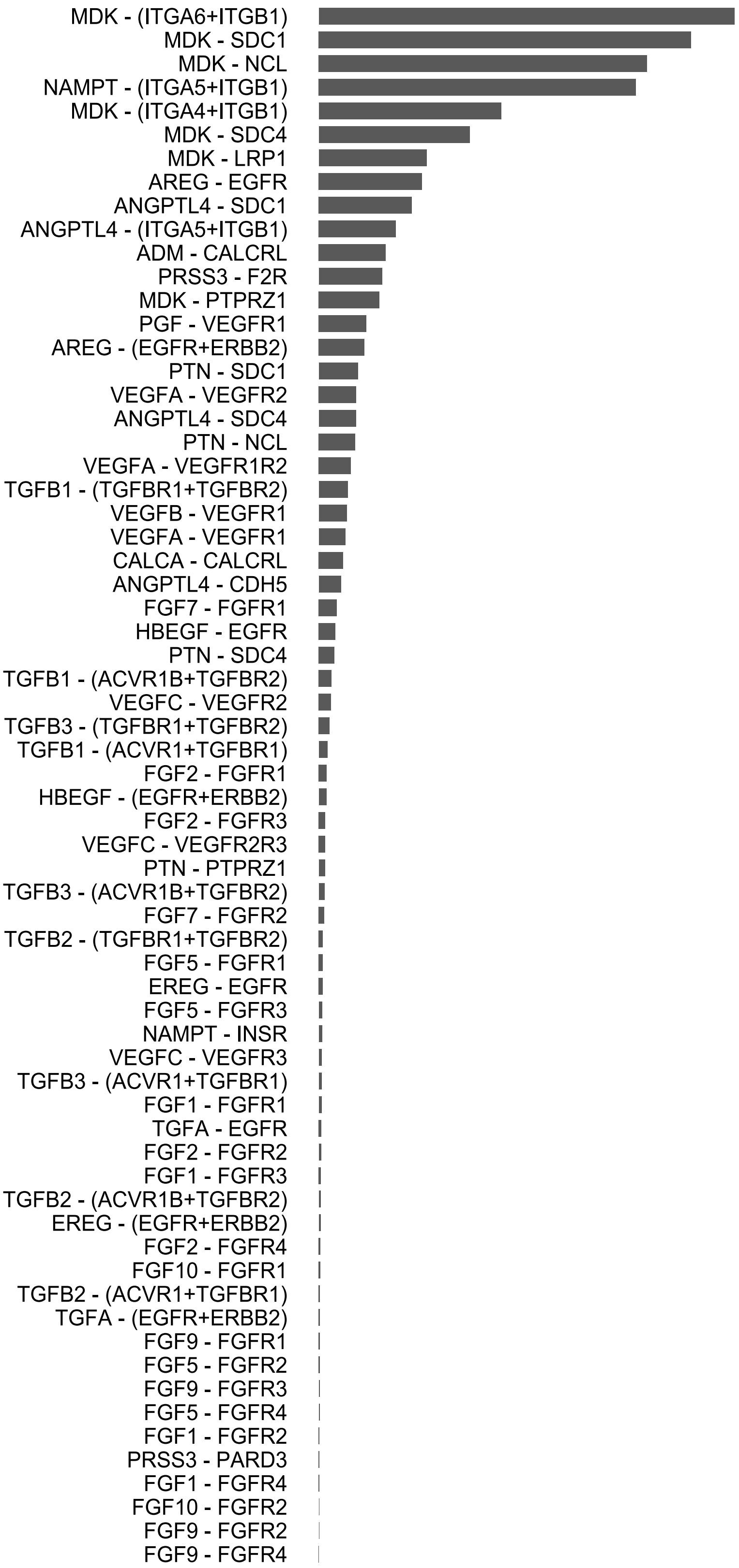

**Supplementary Figure 5. Plot of partial least squares discriminant analysis – epithelial**

PLS-DA scores plot of metabolomic profiles from epithelial effluent samples (n = 5 per group). The plot shows clear separation between uninfected control (green) and infected (red) groups along the first three components. Shaded regions denote 95% confidence ellipsoids for each group.

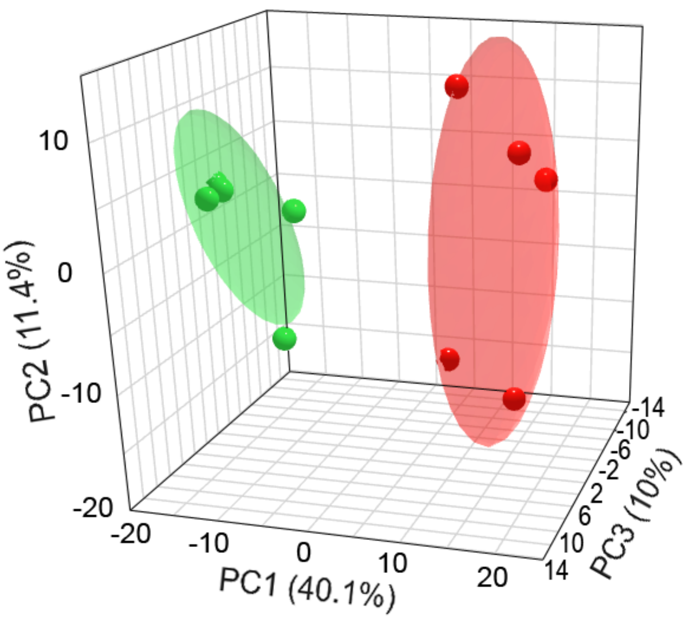

### Supplementary Figure 6. Heatmap of epithelial metabolites

Heatmap of 198 significantly altered metabolites in gingival epithelial cells due to infection. Metabolite abundances are row-scaled (z-score) across samples (blue = lower, red = higher) to highlight infection-associated shifts. Hierarchical clustering shows consistent metabolite signatures that distinguish infected from control epithelial cells.

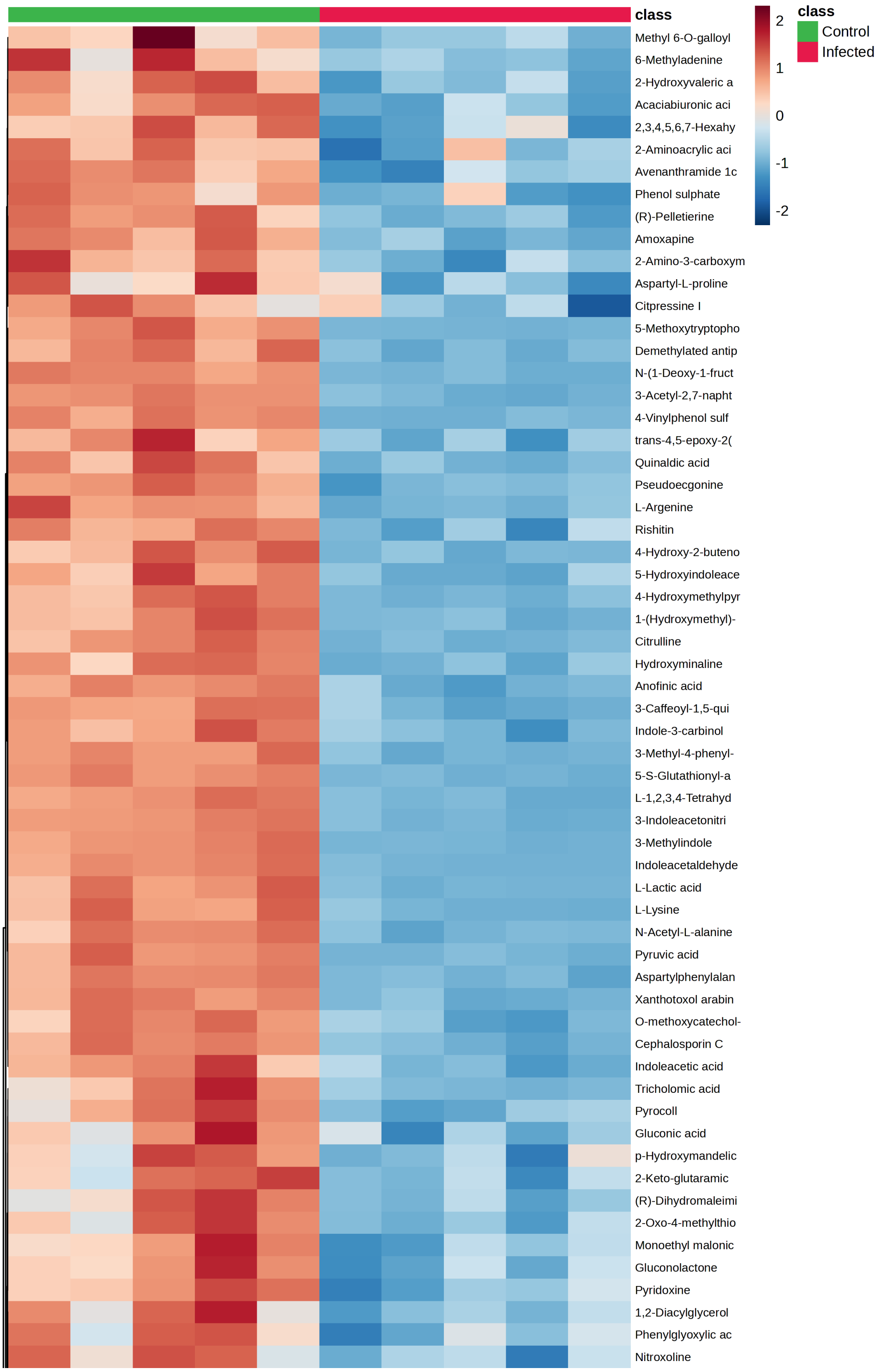

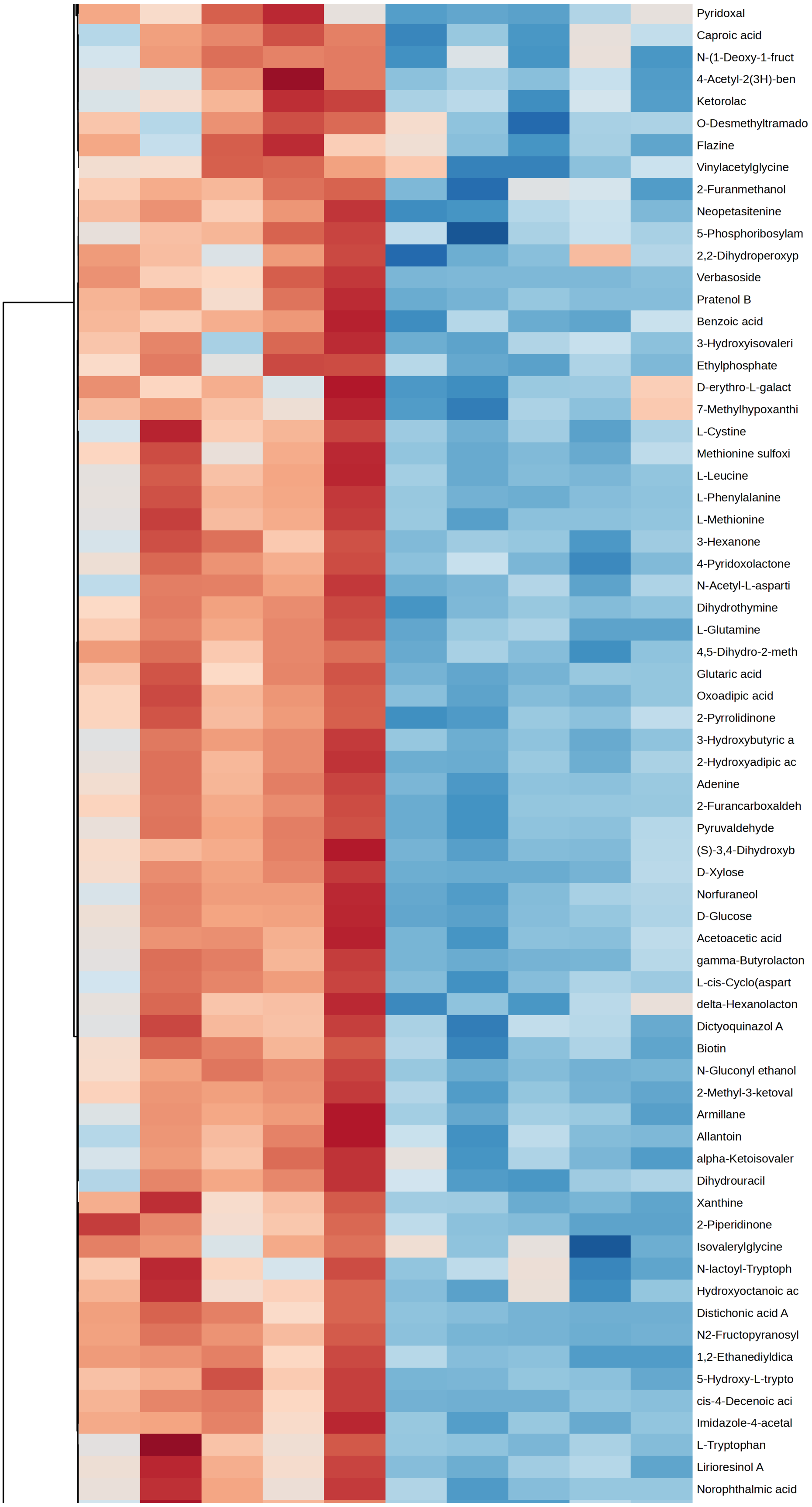

Pyridoxal
Caproic acid
N-(1-Deoxy-1-fruct
4-Acetyl-2(3H)-ben
Ketorolac
O-Desmethyltramado
Flazine
Vinylacetylglycine
2-Furanmethanol
Neopetasitenine
5-Phosphoribosylam
2,2-Dihydroperoxyp
Verbasoside
Pratenol B
Benzoic acid
3-Hydroxyisovaleri
Ethylphosphate
D-erythro-L-galact
7-Methylhypoxanthi
L-Cystine
Methionine sulfoxi
L-Leucine
L-Phenylalanine
L-Methionine
3-Hexanone
4-Pyridoxolactone
N-Acetyl-L-asparti
Dihydrothymine
L-Glutamine
4,5-Dihydro-2-meth
Glutaric acid
Oxoadipic acid
2-Pyrrolidinone
3-Hydroxybutyric a
2-Hydroxyadipic ac
Adenine
2-Furancarboxaldeh
Pyruvaldehyde
(S)-3,4-Dihydroxyb
D-Xylose
Norfuraneol
D-Glucose
Acetoacetic acid
gamma-Butyrolacton
L-cis-Cyclo(aspart
delta-Hexanolacton
Dictyoquinazol A
Biotin
N-Gluconyl ethanol
2-Methyl-3-ketoval
Armillane
Allantoin
alpha-Ketoisovaler
Dihydrouracil
Xanthine
2-Piperidinone
Isovalerylglycine
N-lactoyl-Tryptoph
Hydroxyoctanoic ac
Distichonic acid A
N2-Fructopyranosyl
1,2-Ethanediyldica
5-Hydroxy-L-trypto
cis-4-Decenoic aci
Imidazole-4-acetal
L-Tryptophan
Lirioresinol A
Norophthalmic acid

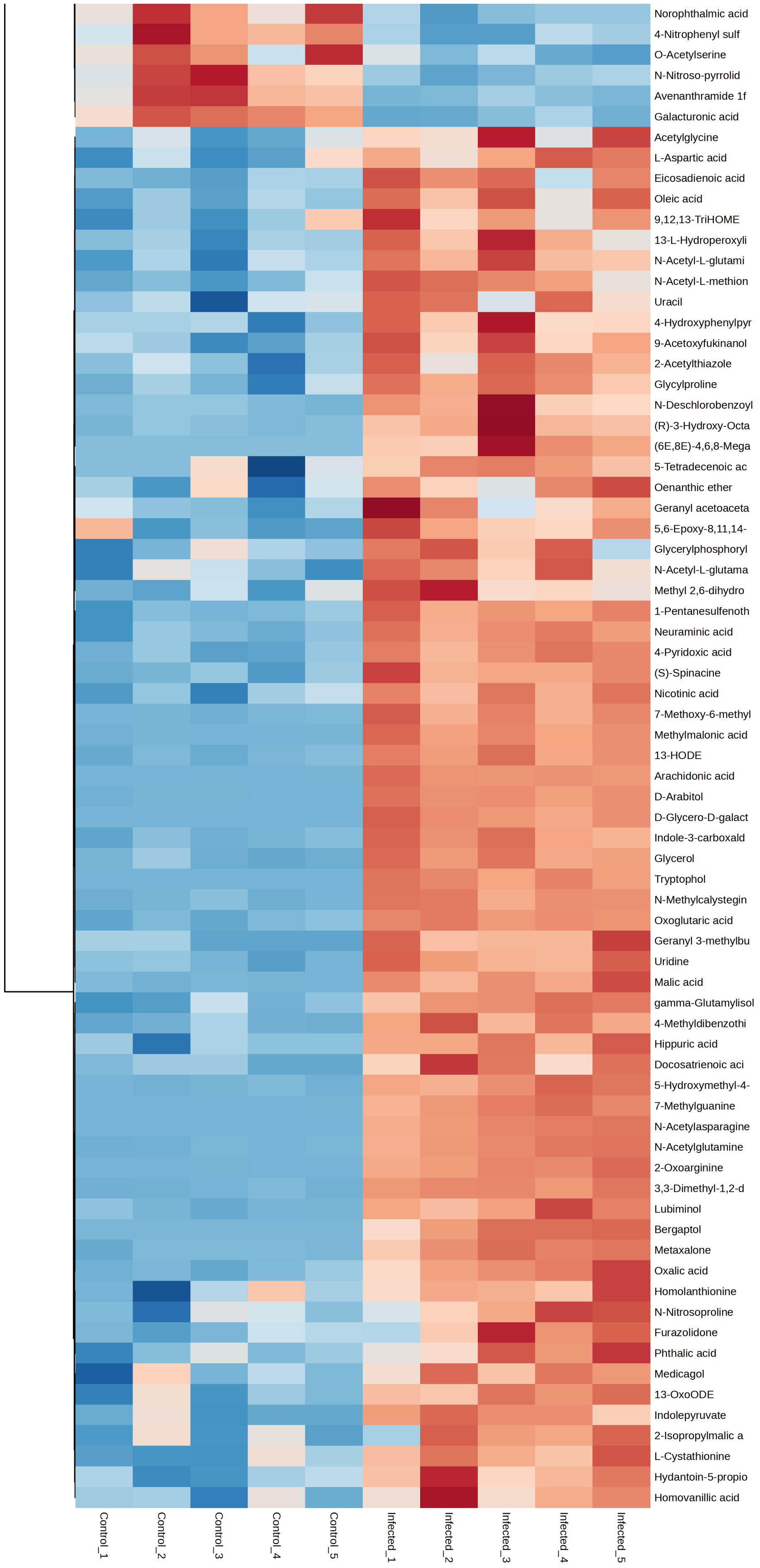
Norophthalmic acid
4-Nitrophenyl sulf
O-Acetylserine
N-Nitroso-pyrrolid
Avenanthramide 1f
Galacturonic acid
Acetylglycine
L-Aspartic acid
Eicosadienoic acid
Oleic acid
9,12,13-TriHOME
13-L-Hydroperoxyli
N-Acetyl-L-glutami
N-Acetyl-L-methion
Uracil
4-Hydroxyphenylpyr
9-Acetoxyfukinanol
2-Acetylthiazole
Glycylproline
N-Deschlorobenzoyl
(R)-3-Hydroxy-Octa
(6E,8E)-4,6,8-Mega
5-Tetradecenoic ac
Oenanthic ether
Geranyl acetoaceta
5,6-Epoxy-8,11,14-
Glycerylphosphoryl
N-Acetyl-L-glutama
Methyl 2,6-dihydro
1-Pentanesulfenoth
Neuraminic acid
4-Pyridoxic acid
(S)-Spinacine
Nicotinic acid
7-Methoxy-6-methyl
Methylmalonic acid
13-HODE
Arachidonic acid
D-Arabitol
D-Glycero-D-galact
Indole-3-carboxald
Glycerol
Tryptophol
N-Methylcalystegin
Oxoglutaric acid
Geranyl 3-methylbu
Uridine
Malic acid
gamma-Glutamylisol
4-Methyldibenzothi
Hippuric acid
Docosatrienoic aci
5-Hydroxymethyl-4-
7-Methylguanine
N-Acetylasparagine
N-Acetylglutamine
2-Oxoarginine
3,3-Dimethyl-1,2-d
Lubiminol
Bergaptol
Metaxalone
Oxalic acid
Homolanthionine
N-Nitrosoproline
Furazolidone
Phthalic acid
Medicagol
13-OxoODE
Indolepyruvate
2-Isopropylmalic a
L-Cystathionine
Hydantoin-5-propio
Homovanillic acid
Control_1
Control_2
Control_3
Control_4
Control_5
Infected_1
Infected_2
Infected_3
Infected_4
Infected_5

### Supplementary Figure 7. Plot of partial least squares discriminant analysis – stromal

PLS-DA scores plot of metabolomic profiles from stromal effluent samples (n = 5 per group). The plot shows clear separation between uninfected control (green) and infected (red) groups along the first three components. Shaded regions denote 95% confidence ellipsoids for each group.

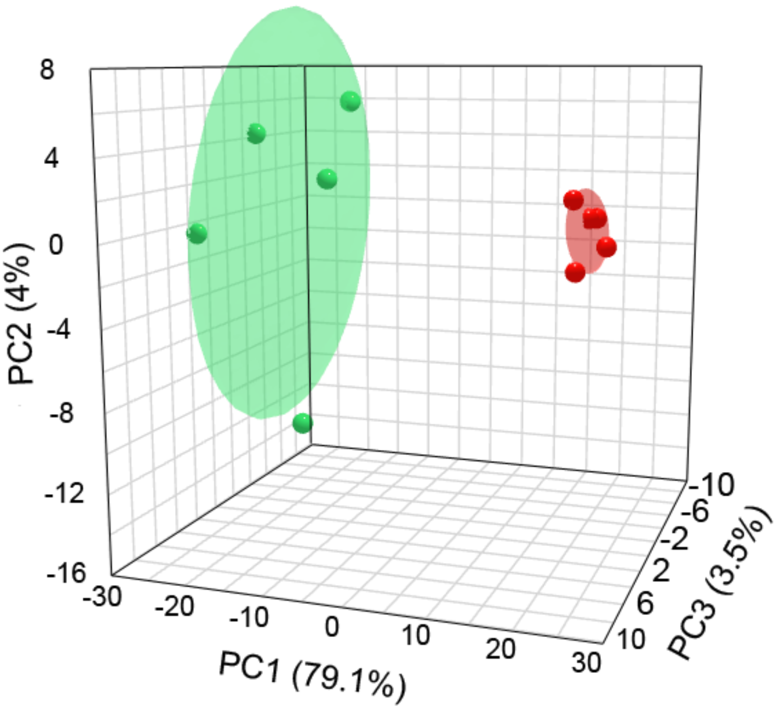

### Supplementary Figure 8. Heatmap of stromal metabolites

Heatmap of 518 significantly altered metabolites in the stromal compartment due to infection. Metabolite abundances are row-scaled (z-score) across samples (blue = lower, red = higher) to highlight infection-associated shifts. Hierarchical clustering shows consistent metabolite signatures that distinguish infected from control epithelial cells.

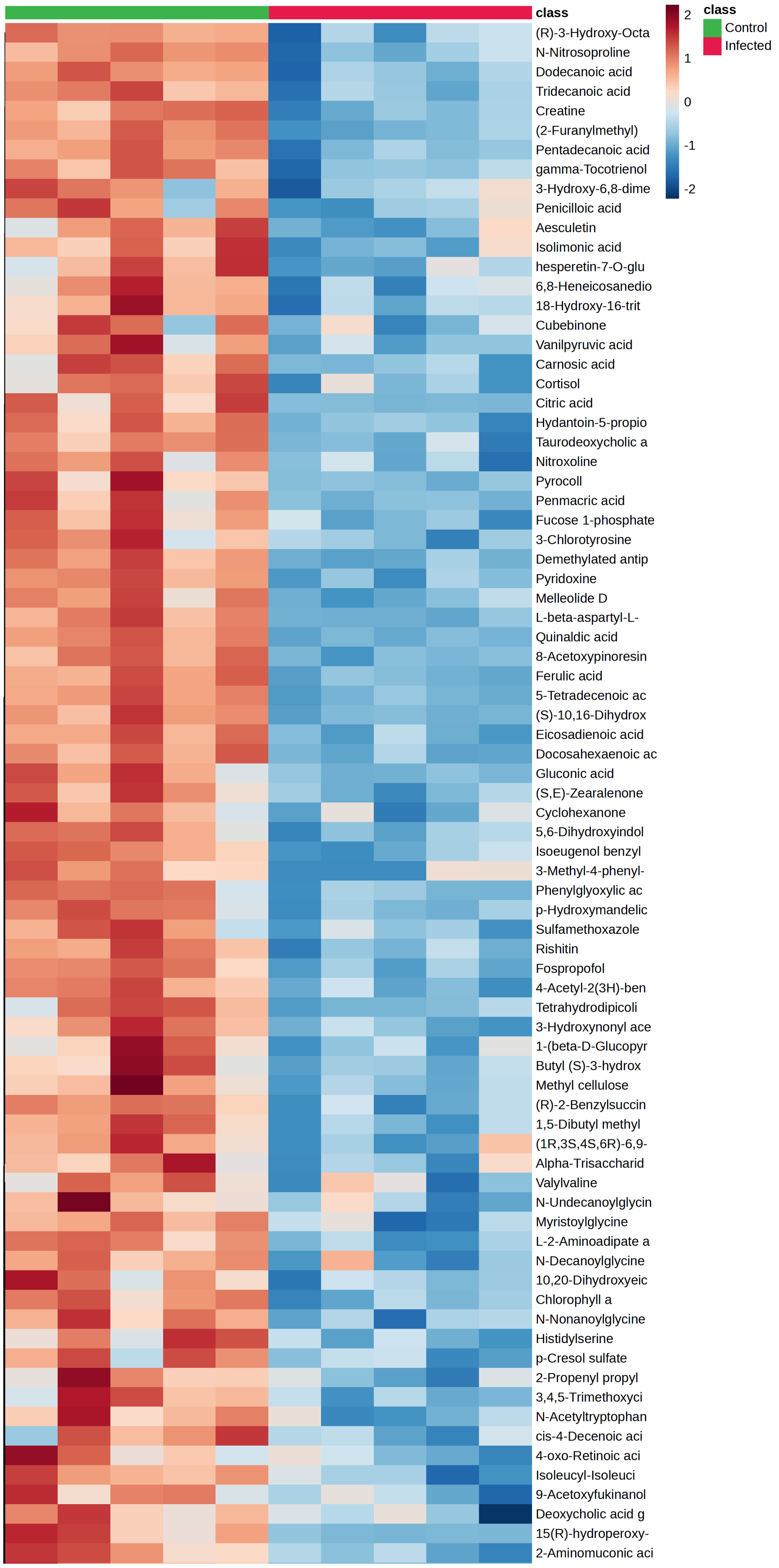

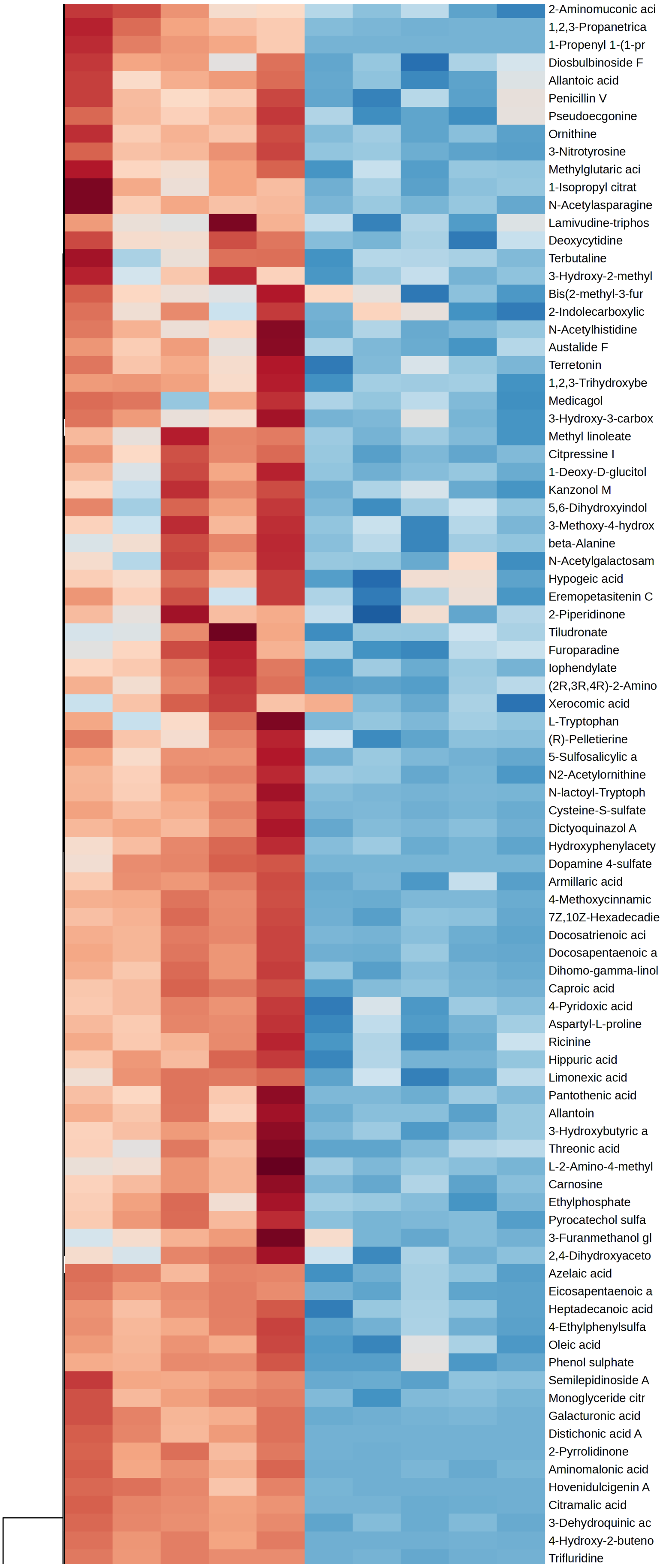
2-Aminomuconic aci
1,2,3-Propanetrica
1-Propenyl 1-(1-pr
Diosbulbinoside F
Allantoic acid
Penicillin V
Pseudoecgonine
Ornithine
3-Nitrotyrosine
Methylglutaric aci
1-Isopropyl citrat
N-Acetylasparagine
Lamivudine-triphos
Deoxycytidine
Terbutaline
3-Hydroxy-2-methyl
Bis(2-methyl-3-fur
2-Indolecarboxylic
N-Acetylhistidine
Austalide F
Terretonin
1,2,3-Trihydroxybe
Medicagol
3-Hydroxy-3-carbox
Methyl linoleate
Citpressine I
1-Deoxy-D-glucitol
Kanzonol M
5,6-Dihydroxyindol
3-Methoxy-4-hydrox
beta-Alanine
N-Acetylgalactosam
Hypogeic acid
Eremopetasitenin C
2-Piperidinone
Tiludronate
Furoparadine
Iophendylate
(2R,3R,4R)-2-Amino
Xerocomic acid
L-Tryptophan
(R)-Pelletierine
5-Sulfosalicylic a
N2-Acetylornithine
N-lactoyl-Tryptoph
Cysteine-S-sulfate
Dictyoquinazol A
Hydroxyphenylacety
Dopamine 4-sulfate
Armillaric acid
4-Methoxycinnamic
7Z,10Z-Hexadecadie
Docosatrienoic aci
Docosapentaenoic a
Dihomo-gamma-linol
Caproic acid
4-Pyridoxic acid
Aspartyl-L-proline
Ricinine
Hippuric acid
Limonexic acid
Pantothenic acid
Allantoin
3-Hydroxybutyric a
Threonic acid
L-2-Amino-4-methyl
Carnosine
Ethylphosphate
Pyrocatechol sulfa
3-Furanmethanol gl
2,4-Dihydroxyaceto
Azelaic acid
Eicosapentaenoic a
Heptadecanoic acid
4-Ethylphenylsulfa
Oleic acid
Phenol sulphate
Semilepidinoside A
Monoglyceride citr
Galacturonic acid
Distichonic acid A
2-Pyrrolidinone
Aminomalonic acid
Hovenidulcigenin A
Citramalic acid
3-Dehydroquinic ac
4-Hydroxy-2-buteno
Trifluridine

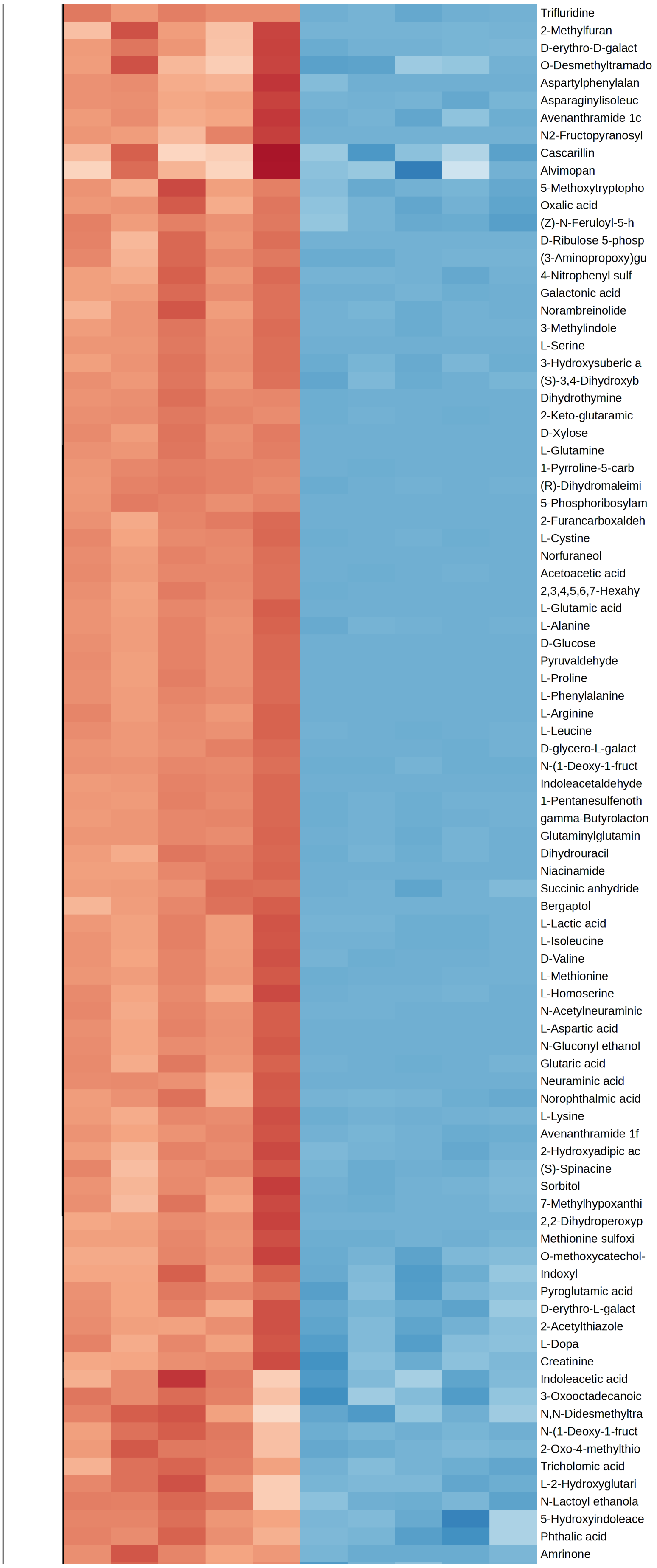
Trifluridine
2-Methylfuran
D-erythro-D-galact
O-Desmethyltramado
Aspartylphenylalan
Asparaginylisoleuc
Avenanthramide 1c
N2-Fructopyranosyl
Cascarillin
Alvimopan
5-Methoxytryptopho
Oxalic acid
(Z)-N-Feruloyl-5-h
D-Ribulose 5-phosp
(3-Aminopropoxy)gu
4-Nitrophenyl sulf
Galactonic acid
Norambreinolide
3-Methylindole
L-Serine
3-Hydroxysuberic a
(S)-3,4-Dihydroxyb
Dihydrothymine
2-Keto-glutaramic
D-Xylose
L-Glutamine
1-Pyrroline-5-carb
(R)-Dihydromaleimi
5-Phosphoribosylam
2-Furancarboxaldeh
L-Cystine
Norfuraneol
Acetoacetic acid
2,3,4,5,6,7-Hexahy
L-Glutamic acid
L-Alanine
D-Glucose
Pyruvaldehyde
L-Proline
L-Phenylalanine
L-Arginine
L-Leucine
D-glycero-L-galact
N-(1-Deoxy-1-fruct
Indoleacetaldehyde
1-Pentanesulfenoth
gamma-Butyrolacton
Glutaminylglutamin
Dihydrouracil
Niacinamide
Succinic anhydride
Bergaptol
L-Lactic acid
L-Isoleucine
D-Valine
L-Methionine
L-Homoserine
N-Acetylneuraminic
L-Aspartic acid
N-Gluconyl ethanol
Glutaric acid
Neuraminic acid
Norophthalmic acid
L-Lysine
Avenanthramide 1f
2-Hydroxyadipic ac
(S)-Spinacine
Sorbitol
7-Methylhypoxanthi
2,2-Dihydroperoxyp
Methionine sulfoxi
O-methoxycatechol-
Indoxyl
Pyroglutamic acid
D-erythro-L-galact
2-Acetylthiazole
L-Dopa
Creatinine
Indoleacetic acid
3-Oxooctadecanoic
N,N-Didesmethyltra
N-(1-Deoxy-1-fruct
2-Oxo-4-methylthio
Tricholomic acid
L-2-Hydroxyglutari
N-Lactoyl ethanola
5-Hydroxyindoleace
Phthalic acid
Amrinone

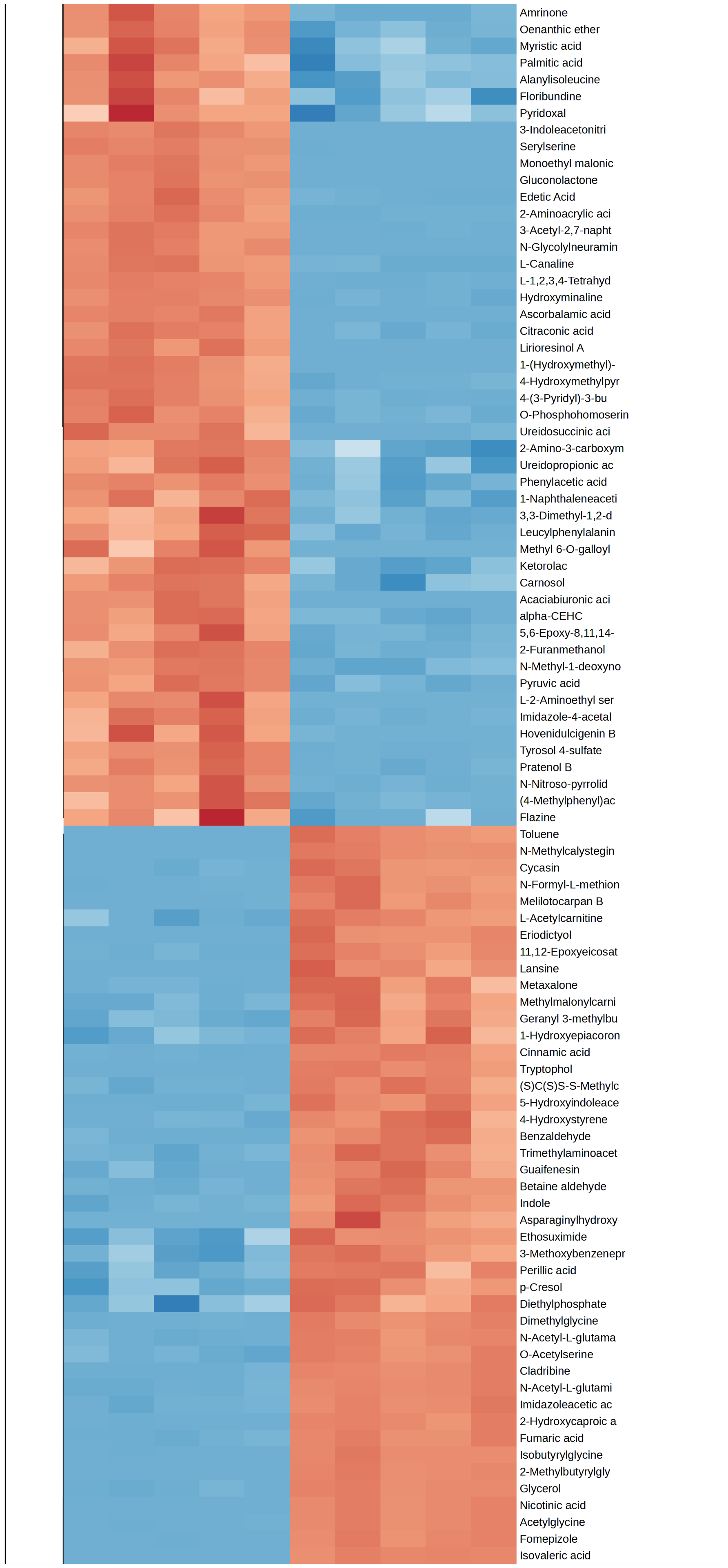
Amrinone
Oenanthic ether
Myristic acid
Palmitic acid
Alanylisoleucine
Floribundine
Pyridoxal
3-Indoleacetonitri
Serylserine
Monoethyl malonic
Gluconolactone
Edetic Acid
2-Aminoacrylic aci
3-Acetyl-2,7-napht
N-Glycolylneuramin
L-Canaline
L-1,2,3,4-Tetrahyd
Hydroxyminaline
Ascorbalamic acid
Citraconic acid
Lirioresinol A
1-(Hydroxymethyl)-
4-Hydroxymethylpyr
4-(3-Pyridyl)-3-bu
O-Phosphohomoserin
Ureidosuccinic aci
2-Amino-3-carboxym
Ureidopropionic ac
Phenylacetic acid
1-Naphthaleneaceti
3,3-Dimethyl-1,2-d
Leucylphenylalanin
Methyl 6-O-galloyl
Ketorolac
Carnosol
Acaciabiuronic aci
alpha-CEHC
5,6-Epoxy-8,11,14-
2-Furanmethanol
N-Methyl-1-deoxyno
Pyruvic acid
L-2-Aminoethyl ser
Imidazole-4-acetal
Hovenidulcigenin B
Tyrosol 4-sulfate
Pratenol B
N-Nitroso-pyrrolid
(4-Methylphenyl)ac
Flazine
Toluene
N-Methylcalystegin
Cycasin
N-Formyl-L-methion
Melilotocarpan B
L-Acetylcarnitine
Eriodictyol
11,12-Epoxyeicosat
Lansine
Metaxalone
Methylmalonylcarni
Geranyl 3-methylbu
1-Hydroxyepiacoron
Cinnamic acid
Tryptophol
(S)C(S)S-S-Methylc
5-Hydroxyindoleace
4-Hydroxystyrene
Benzaldehyde
Trimethylaminoacet
Guaifenesin
Betaine aldehyde
Indole
Asparaginylhydroxy
Ethosuximide
3-Methoxybenzenepr
Perillic acid
p-Cresol
Diethylphosphate
Dimethylglycine
N-Acetyl-L-glutama
O-Acetylserine
Cladribine
N-Acetyl-L-glutami
Imidazoleacetic ac
2-Hydroxycaproic a
Fumaric acid
Isobutyrylglycine
2-Methylbutyrylgly
Glycerol
Nicotinic acid
Acetylglycine
Fomepizole
Isovaleric acid

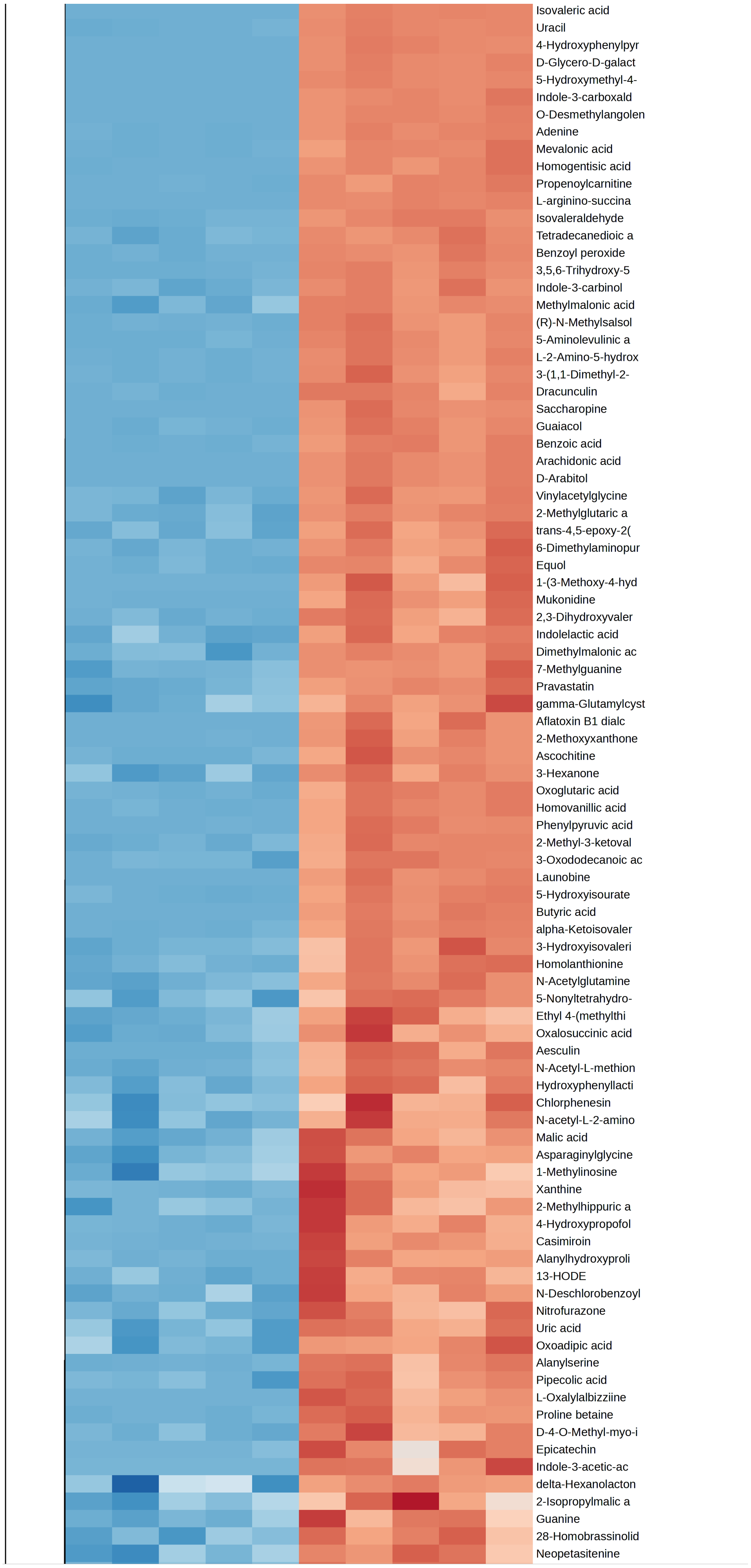

Isovaleric acid
Uracil
4-Hydroxyphenylpyr
D-Glycero-D-galact
5-Hydroxymethyl-4-
Indole-3-carboxald
O-Desmethylangolen
Adenine
Mevalonic acid
Homogentisic acid
Propenoylcarnitine
L-arginino-succina
Isovaleraldehyde
Tetradecanedioic a
Benzoyl peroxide
3,5,6-Trihydroxy-5
Indole-3-carbinol
Methylmalonic acid
(R)-N-Methylsalsol
5-Aminolevulinic a
L-2-Amino-5-hydrox
3-(1,1-Dimethyl-2-
Dracunculin
Saccharopine
Guaiacol
Benzoic acid
Arachidonic acid
D-Arabitol
Vinylacetylglycine
2-Methylglutaric a
trans-4,5-epoxy-2(
6-Dimethylaminopur
Equol
1-(3-Methoxy-4-hyd
Mukonidine
2,3-Dihydroxyvaler
Indolelactic acid
Dimethylmalonic ac
7-Methylguanine
Pravastatin
gamma-Glutamylcyst
Aflatoxin B1 dialc
2-Methoxyxanthone
Ascochitine
3-Hexanone
Oxoglutaric acid
Homovanillic acid
Phenylpyruvic acid
2-Methyl-3-ketoval
3-Oxododecanoic ac
Launobine
5-Hydroxyisourate
Butyric acid
alpha-Ketoisovaler
3-Hydroxyisovaleri
Homolanthionine
N-Acetylglutamine
5-Nonyltetrahydro-
Ethyl 4-(methylthi
Oxalosuccinic acid
Aesculin
N-Acetyl-L-methion
Hydroxyphenyllacti
Chlorphenesin
N-acetyl-L-2-amino
Malic acid
Asparaginylglycine
1-Methylinosine
Xanthine
2-Methylhippuric a
4-Hydroxypropofol
Casimiroin
Alanylhydroxyproli
13-HODE
N-Deschlorobenzoyl
Nitrofurazone
Uric acid
Oxoadipic acid
Alanylserine
Pipecolic acid
L-Oxalylalbizziine
Proline betaine
D-4-O-Methyl-myo-i
Epicatechin
Indole-3-acetic-ac
delta-Hexanolacton
2-Isopropylmalic a
Guanine
28-Homobrassinolid
Neopetasitenine

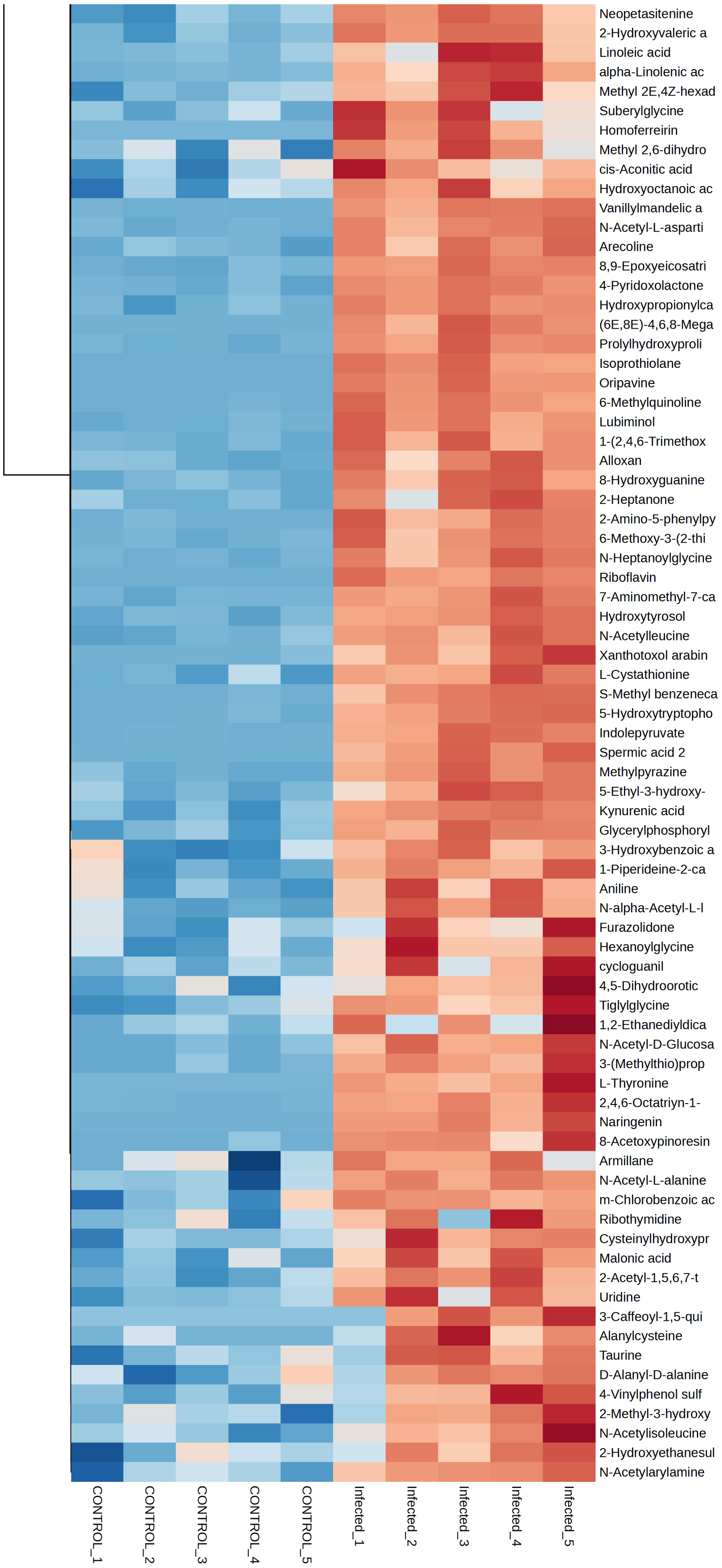
Neopetasitenine
2-Hydroxyvaleric a
Linoleic acid
alpha-Linolenic ac
Methyl 2E,4Z-hexad
Suberylglycine
Homoferreirin
Methyl 2,6-dihydro
cis-Aconitic acid
Hydroxyoctanoic ac
Vanillylmandelic a
N-Acetyl-L-asparti
Arecoline
8,9-Epoxyeicosatri
4-Pyridoxolactone
Hydroxypropionylca
(6E,8E)-4,6,8-Mega
Prolylhydroxyproli
Isoprothiolane
Oripavine
6-Methylquinoline
Lubiminol
1-(2,4,6-Trimethox
Alloxan
8-Hydroxyguanine
2-Heptanone
2-Amino-5-phenylpy
6-Methoxy-3-(2-thi
N-Heptanoylglycine
Riboflavin
7-Aminomethyl-7-ca
Hydroxytyrosol
N-Acetylleucine
Xanthotoxol arabin
L-Cystathionine
S-Methyl benzeneca
5-Hydroxytryptopho
Indolepyruvate
Spermic acid 2
Methylpyrazine
5-Ethyl-3-hydroxy-
Kynurenic acid
Glycerylphosphoryl
3-Hydroxybenzoic a
1-Piperideine-2-ca
Aniline
N-alpha-Acetyl-L-l
Furazolidone
Hexanoylglycine
cycloguanil
4,5-Dihydroorotic
Tiglylglycine
1,2-Ethanediyldica
N-Acetyl-D-Glucosa
3-(Methylthio)prop
L-Thyronine
2,4,6-Octatriyn-1-
Naringenin
8-Acetoxypinoresin
Armillane
N-Acetyl-L-alanine
m-Chlorobenzoic ac
Ribothymidine
Cysteinylhydroxypr
Malonic acid
2-Acetyl-1,5,6,7-t
Uridine
3-Caffeoyl-1,5-qui
Alanylcysteine
Taurine
D-Alanyl-D-alanine
4-Vinylphenol sulf
2-Methyl-3-hydroxy
N-Acetylisoleucine
2-Hydroxyethanesul
N-Acetylarylamine
CONTROL_1
CONTROL_2
CONTROL_3
CONTROL_4
CONTROL_5
Infected_1
Infected_2
Infected_3
Infected_4
Infected_5

### Supplementary Figure 9. Top-ranked differentially regulated epithelial metabolites

Top 10 upregulated and top 10 downregulated metabolites in the gingival epithelial compartment of the mouth-on-a-chip following *C. albicans* infection. Control denotes uninfected gingival tissue. Data represent mean ± SD and were normalized to the median (n = 5 per group). Box plots show minimum, 25th percentile, mean, and 75th percentile. Statistical significance is indicated as ns (not significant), *$P < 0.05$*, and **$P < 0.01$*.

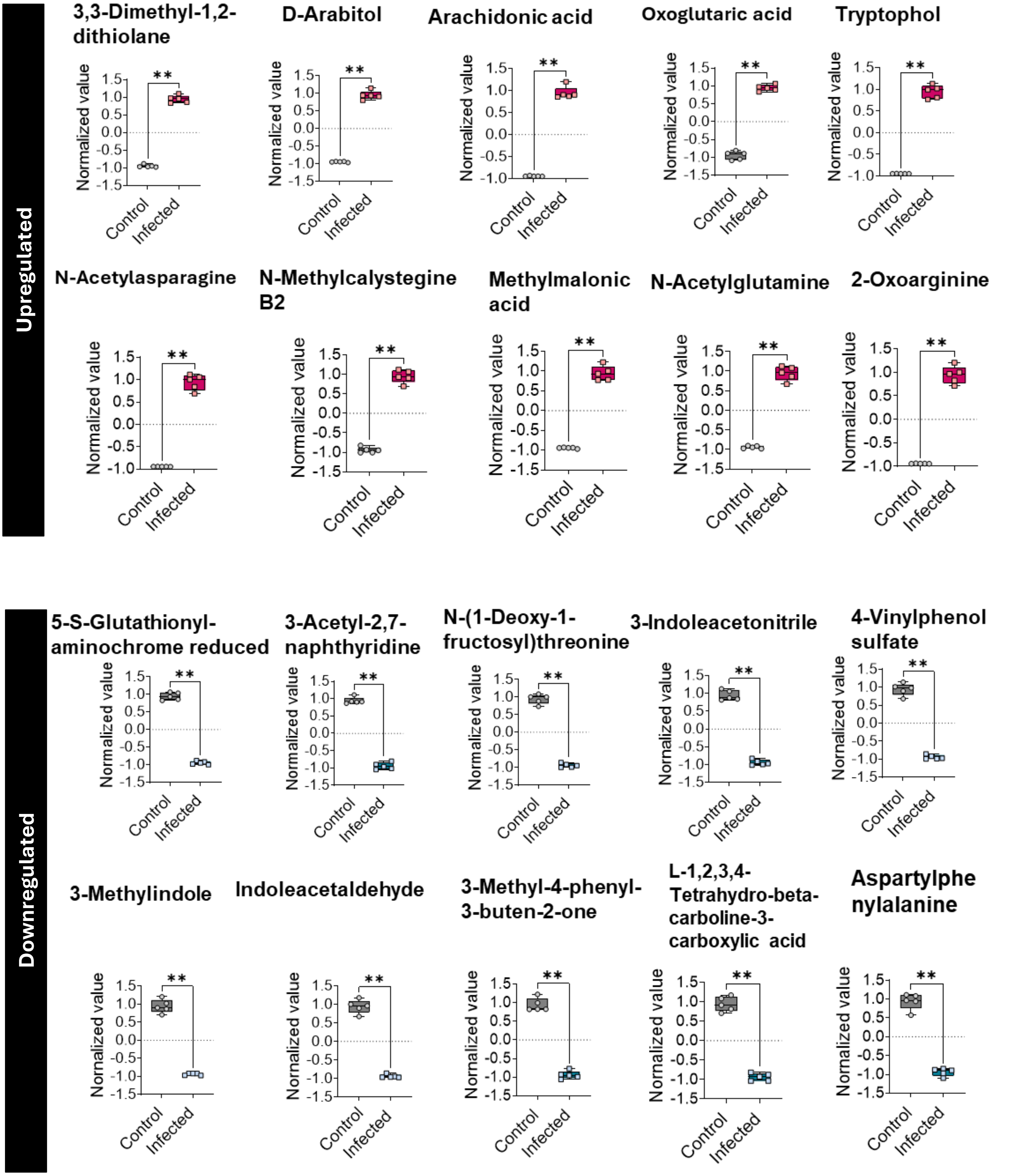

### Supplementary Figure 10. Top-ranked differentially regulated stromal metabolites

Top 10 upregulated and top 10 downregulated metabolites in the gingival stromal compartment of the mouth-on-a-chip following *C. albicans* infection. Control denotes uninfected gingival tissue. Data represent mean ± SD and were normalized to the median (n = 5 per group). Box plots show minimum, 25th percentile, mean, and 75th percentile. Statistical significance is indicated as ns (not significant), *$P < 0.05$*, and **$P < 0.01$*.

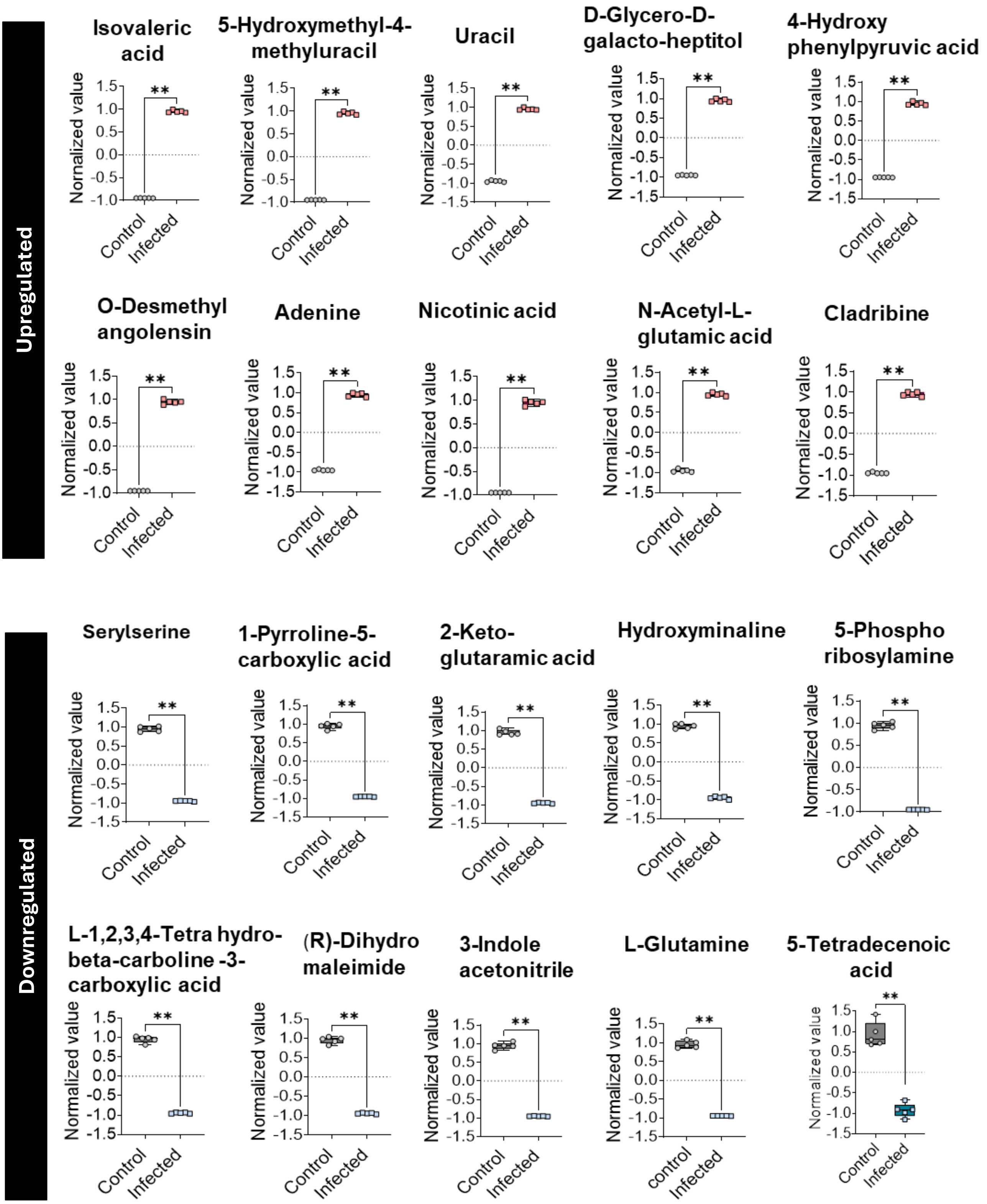

### Supplementary Figure 11. Metabolite biomarker analysis of the infected gingival epithelium

**a.** Receive operating characteristic (ROC) curves derived from PLS-DA models to identify metabolite features that discriminate infected from uninfected conditions. Each curve represents a model constructed with a different number of metabolite features (5-100) with AUC and 95% CI values shown in the legend. **b.** Top discriminatory metabolites ranked by VIP scores. The accompanying heatmap on the right shows relative metabolite abundance.

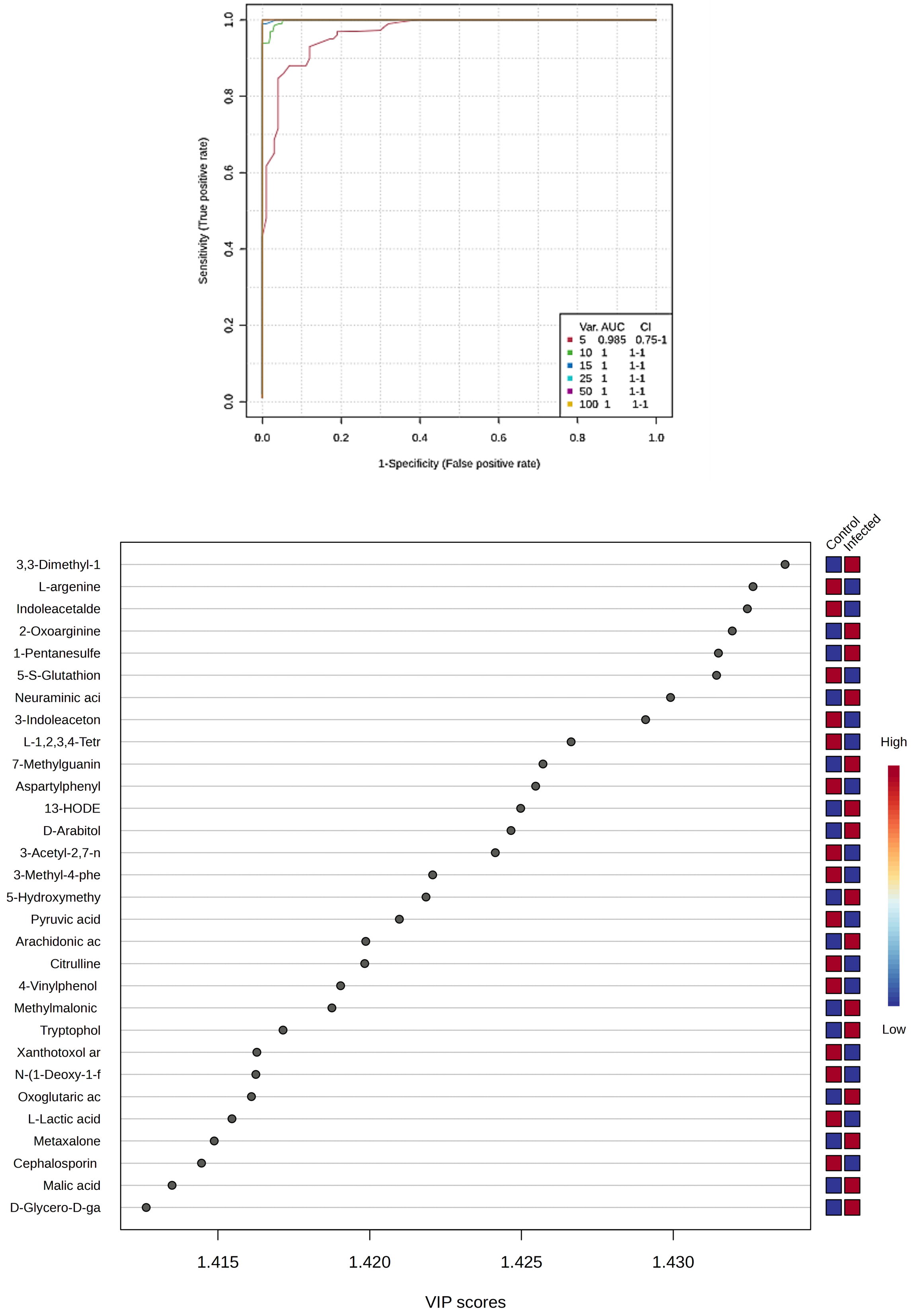

### Supplementary Figure 12. Metabolite biomarker analysis of the infected gingival stroma

**a.** Receive operating characteristic (ROC) curves derived from PLS-DA models to identify metabolite features that discriminate infected from uninfected conditions. Each curve represents a model constructed with a different number of metabolite features (5-100) with AUC and 95% CI values shown in the legend. **b.** Top discriminatory metabolites ranked by VIP scores. The accompanying heatmap on the right shows relative metabolite abundance.

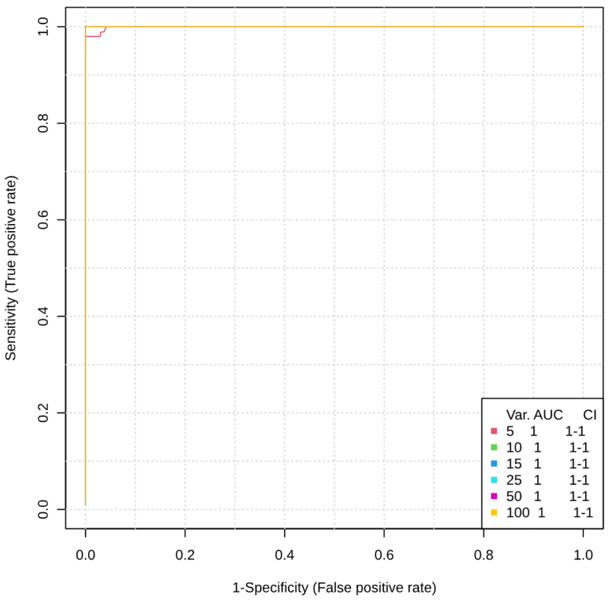


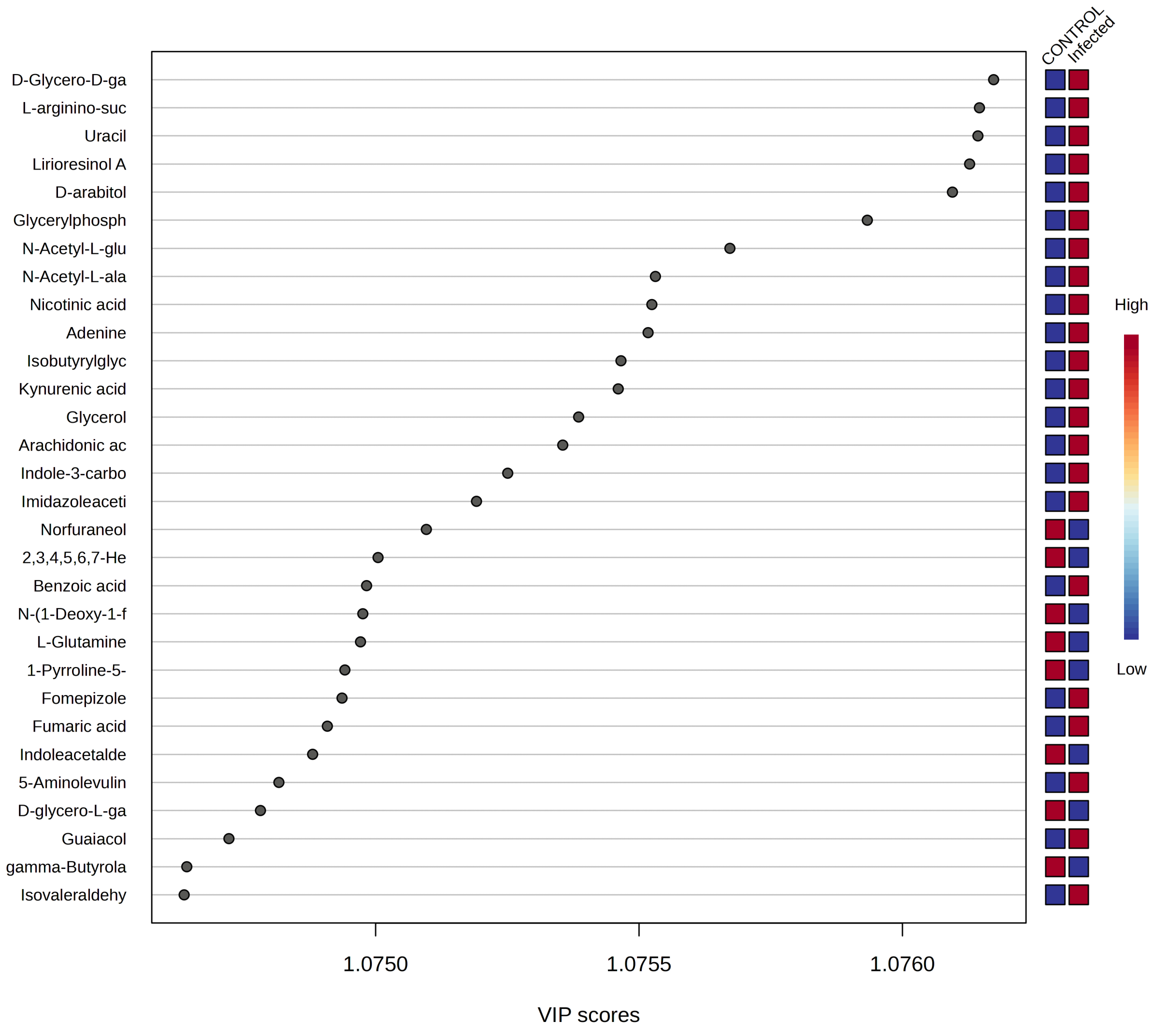

### Supplementary Figure 13. Top five discriminatory epithelial metabolites

Top five metabolite biomarkers identified from the epithelial compartment that discriminate infected from uninfected control conditions with the highest levels of significance. Data represent mean ± SD and were normalized to the median (n = 5 per group). Box plots show minimum, 25th percentile, mean, and 75th percentile. Statistical significance is indicated as: ns (not significant), *$P < 0.05$, **$P < 0.01$.

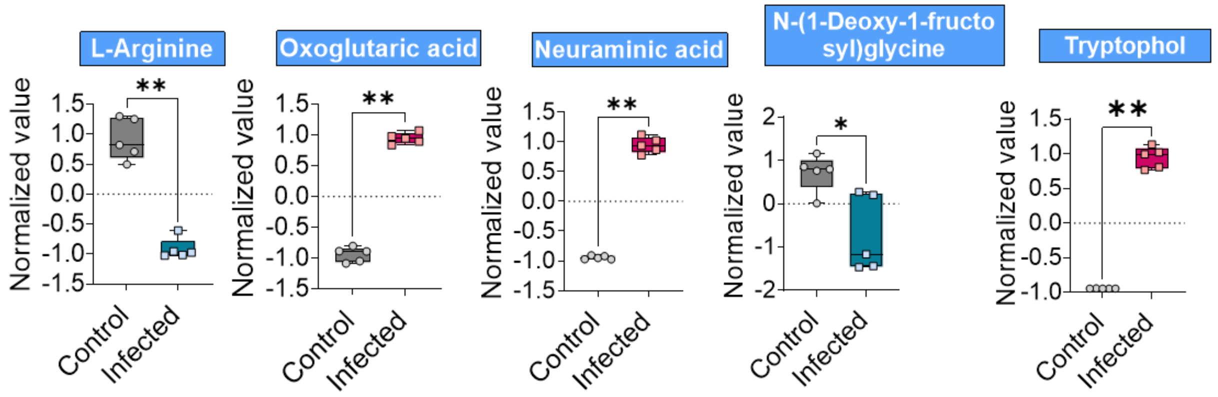

### Supplementary Figure 14. Top five discriminatory stromal metabolites

Top five metabolite biomarkers identified from the stromal compartment that discriminate infected from uninfected control conditions with the highest levels of significance. Data represent mean ± SD and were normalized to the median (n = 5 per group). Box plots show minimum, 25th percentile, mean, and 75th percentile. Statistical significance is indicated as: ns (not significant), *$P < 0.05$, **$P < 0.01$.

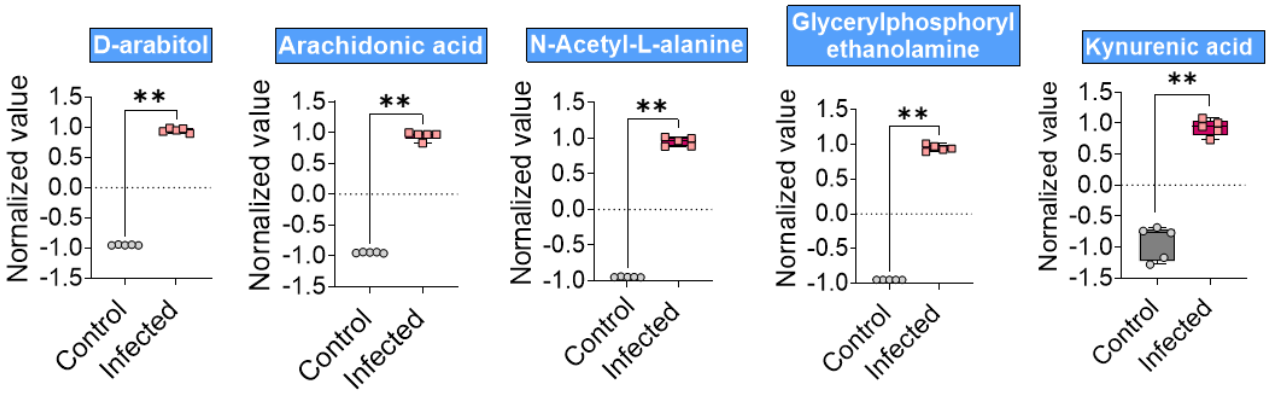

### Supplementary Figure 15. Device design to model salivary flow

**a.** Illustration of the mouth-on-a-chip containing a network of microchannels to generate salivary flow. **b,c,** Top and cross-sectional views of devices used to model normal (**b**) and hyposalivation (**c**) conditions. The difference between the two designs is the diameter of the microchannels used for delivery of human saliva (100 µm for normal and 40 µm for hyposalivation.

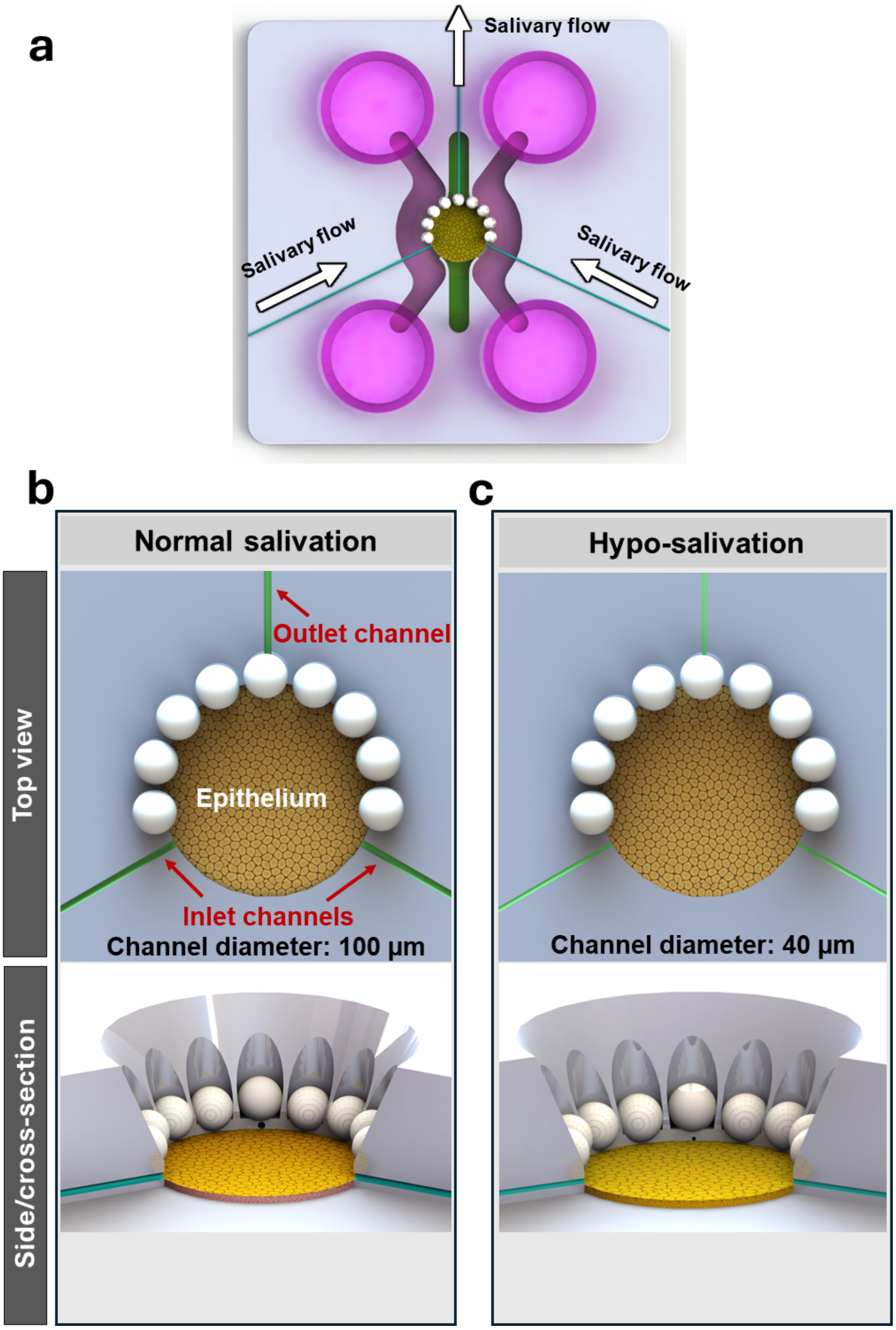

### Supplementary Table 1. Key resources

| Reagent or resource | Source | Identifier/Catalog |
|---|---|---|
| Antibodies | | |
| **Alexa Fluor 488 Anti-CD31** | abcam | ab215911 |
| **Anti-ZO-1** | Invitrogen | 61-7300 |
| **Anti-ZO-1** | Invitrogen | 33-9100 |
| **Alexa Fluor™ 488 Phalloidin** | ThermoFisher Scientific | A-12379 |
| **Anti-cytokeratin-10 (CK10) antibody** | Thermo Fisher Scientific | MA1-06319 |
| **Anti-cytokeratin-14 (CK14) antibody** | Thermo Fisher Scientific | MA5-32214 |
| **Anti-Occludin antibody** | Thermo Fisher Scientific | 40-4700 |
| **Anti-Claudin 1 antibody (2H10D10)** | Thermo Fisher Scientific | 37-4900 |
| **Anti-vimentin antibody** | Thermo Fisher Scientific | CL48880232100UL |
| **Anti-ACE2 antibody** | Thermo Fisher Scientific | MA5-32307 |
| **Anti-furin antibody** | Thermo Fisher Scientific | PA5-96680 |
| **Anti-SARS spike glycoprotein antibody [1A9]** | abcam | ab273433 |
| **Anti-ICAM-1 antibody** | Invitrogen | MA5407 |
| **Donkey anti-mouse secondary antibody, Alexa Fluor Plus 555** | Invitrogen | A-32773 |
| **Goat anti-mouse secondary antibody, Alexa Fluor Plus 488** | Invitrogen | A-32723 |
| **Goat anti-mouse secondary antibody, Alexa Fluor Plus 647** | Invitrogen | A-32728 |
| **Goat anti-rabbit secondary antibody, Alexa Fluor Plus 555** | Invitrogen | 2732 |
| **Donkey anti-rabbit secondary antibody, Alexa Fluor 488** | Invitrogen | A-32790 |
| **Hoechst nuclear stain** | Thermo Fisher Scientific | D1306 |
| **CellTracker Deep Red** | Thermo Fisher Scientific | C34565 |
| **Chemicals, peptides, and recombinant proteins** | | |
| **DMEM/F-12, HEPES, no phenol red** | Gibco/Invitrogen | 11039021 |
| **EGM™-2 BulletKit™** | Lonza | CC-3162 |
| **FGM™-2 BulletKit™** | Lonza | CC-3132 |
| **DermaLife K growth medium** | Lifeline Cell Technology | LL-0007 |
| **Gingival epithelial differentiation medium** | CELLNTEC | CnT-PR-3D |
| **EasySep™ Direct Human Neutrophil Isolation Kit** | STEMCELL Technologies | 100-0404 |
| **Phosphate-buffered saline (PBS)** | gibco | 14190-136 |
| **Paraformaldehyde, 4%** | Thermo Scientific | J19943-K2 |
| **Fibrin** | Sigma-Aldrich | F8630-1G |
| **Thrombin** | Sigma-Aldrich | T7513 |
| **Aprotinin** | Sigma-Aldrich | A1153 |
| **Fibronectin** | Sigma-Aldrich | F1141 |
| **Trypsin** | gibco | 25200-056 |
| **FITC-labeled dextran, 70 kDa** | Sigma-Aldrich | 46945-100MG-F |
| **Lucifer Yellow CH, lithium salt** | Invitrogen | L453 |
| **Mounting medium / coverslip mounting reagent** | Dako | S3023 |
| **Histological clearing agent** | National Diagnostics | HS-200 |
| **Triton X-100** | Sigma-Aldrich | T8787-100ML |
| **Haematoxylin solution** | Electron Microscopy Science | 26030-20 |
| **Eosin solution** | Sigma-Aldrich | HT110280-2.5L |
| **Bovine serum albumin (BSA)** | Sigma-Aldrich | A7906-100G |
| **iScript cDNA Synthesis Kit** | Bio-Rad | 1708891 |
| **TaqMan Gene Expression Assays** | Thermo Fisher Scientific | 4331182 |
| **Human IL-6 ELISA kit** | Sigma Aldrich | RAB0306 |
| **Human IL-8 / CXCL8 ELISA Kit** | Sigma Aldrich | RAB0319 |
| **Human TNF-α ELISA Kit** | abcam | Ab181421 |
| **Human IL-1β ELISA Kit** | Sigma Aldrich | RAB0273 |
| **Human TGF-β1 DueSet ELISA** | R&D Systems | DY240 |
| **Cytotoxicity Detection Kit$^{PLUS}$ (LDH)** | Roche | 04744926001 |
| Cells and biological samples | | |
| **Primary human gingival epithelial progenitor cells** | Lifeline Cell Technologies | FC-0094 |
| **Primary human gingival fibroblasts** | Lifeline Cell Technologies | FC-0095 |
| **Human Cord Blood CD34+ Cells, Frozen** | STEMCELL Technologies | 70008 |

| | | |
|---|---|---|
| **Human peripheral blood neutrophils** | Human Immunology Core, University of Pennsylvania | Apheresis products |
| **Pooled sterile human saliva** | Innovative Research | IRHUSL50ML |
| **SARS-CoV-2 pseudovirus carrying fluorescent reporter** | Montana Molecular | C1110G |
| **Microbial Strain** | | |
| ***C. albicans*** | Biofilm Research Lab at UPENN | SN250-tdTomato |
| ***S. mutans*** | Biofilm Research Lab at UPENN | UA159-GFP |
| **Equipment and instruments** | | |
| **Laser scanning confocal microscope** | Carl Zeiss | LSM 800 |
| **Upright fluorescence microscope** | Zeiss | AxioZoom |
| **Digital ohmmeter for TEER measurements** | FLUKE Corp. | RMS multimeter |
| **UV Sterilization system** | Electro-lite | ELC-500 |
| **Plate reader** | Tecan | M200 |
| **Microtome (Hm325)** | MICROM GmbH | 902120 |
| **Software and algorithms** | | |
| **AngioTool** | National Cancer Institute / AngioTool | v0.6a |
| **ZEN software** | Zeiss | v3.11 |
| **ImageJ** | NIH | v1.54g |
| **BiofilmQ** | Drescher Lab | v1.0.2 |
| **Cell Ranger pipeline** | 10x Genomics | v5.0.0 |
| **Seurat package** | Satija Lab | v5.0.1 |
| **Monocle 3** | Trapnell Lab | v1.0.0 |
| **clusterProfiler R package** | Bioconductor | v4.6.0 |
| **Molecular Signatures Database (MSigDB)** | Broad Institute | v7.5.1 |
| **CellChatDB** | CellChat | v2.0 |
| **ProteoWizard** | ProteoWizard | v3.0.20315 |
| **El-MAVEN** | El-MAVEN | v0.12.0 |
| **MetaboAnalyst** | MetaboAnalyst | v5.0 |
| **GraphPad Prism** | GraphPad Software | v10 |

### Supplementary Table 2. List of primers

| Gene name | Source | Identifier/Catalog |
|---|---|---|
| **GAPDH** | Sino Biological | HP100003 |
| **IL-6** | Sino Biological | HP100427 |
| **TNF/TNFA** | Sino Biological | HP100592 |
| **CXCL8/IL8** | Sino Biological | HP100179 |
| **IL-1B** | Sino Biological | HP100210 |
| **INFA1** | Sino Biological | HP104901 |
| **INFB1** | Sino Biological | HP100883 |
| **INFG** | Sino Biological | HP102720 |
| **CXCL10/IP-10** | Sino Biological | HP100690 |
